# Hypnotic treatment reverses NREM sleep disruption and EEG desynchronization in a mouse model of Fragile X syndrome to rescue memory consolidation deficits

**DOI:** 10.1101/2023.07.14.549070

**Authors:** Jessy D. Martinez, Lydia G. Wilson, William P. Brancaleone, Kathryn G. Peterson, Donald S. Popke, Valentina Caicedo Garzon, Roxanne E. Perez Tremble, Marcus J. Donnelly, Stephany L. Mendez Ortega, Daniel Torres, James J. Shaver, Brittany C. Clawson, Sha Jiang, Zhongying Yang, Sara J. Aton

**Affiliations:** Department of Molecular, Cellular, and Developmental Biology, University of Michigan, Ann Arbor, MI 48109; Undergraduate Program in Neuroscience, University of Michigan, Ann Arbor, MI 48109

## Abstract

Fragile X syndrome (FXS) is a highly-prevalent genetic cause of intellectual disability, associated with disrupted cognition and sleep abnormalities. Sleep loss itself negatively impacts cognitive function, yet the contribution of sleep loss to impaired cognition in FXS is vastly understudied. One untested possibility is that disrupted cognition in FXS is exacerbated by abnormal sleep. We hypothesized that restoration of sleep-dependent mechanisms could improve functions such as memory consolidation in FXS. We examined whether administration of ML297, a hypnotic drug acting on G-protein-activated inward-rectifying potassium channels, could restore sleep phenotypes and improve disrupted memory consolidation in *Fmr1^-/y^* mice. Using 24-h polysomnographic recordings, we found that *Fmr1^-/y^* mice exhibit reduced non-rapid eye movement (NREM) sleep and fragmented NREM sleep architecture, alterations in NREM EEG spectral power (including reductions in sleep spindles), and reduced EEG coherence between cortical areas. These alterations were reversed in the hours following ML297 administration. Hypnotic treatment following contextual fear or spatial learning also ameliorated disrupted memory consolidation in *Fmr1^-/y^* mice. Hippocampal activation patterns during memory recall was altered in *Fmr1^-/y^* mice, reflecting an altered balance of activity among principal neurons vs. parvalbumin-expressing (PV+) interneurons. This phenotype was partially reversed by post-learning ML297 administration. These studies suggest that sleep disruption could have a major impact on neurophysiological and behavioral phenotypes in FXS, and that hypnotic therapy may significantly improve disrupted cognition in this disorder.

## Introduction

FXS is an X-linked neurodevelopmental disorder resulting from silencing of the *FMR1* gene and loss of Fragile X messenger Ribonucleoprotein (FMRP). The leading cause of both heritable intellectual disability and syndromic autism spectrum disorder (ASD)^1, 2^, FXS is characterized by altered sensory processing, hyperactivity, and cognitive impairments^3–5^. Available data suggest that FXS patients have trouble falling asleep and frequent nighttime awakenings^6–9^ – a phenotype shared with other neurodevelopmental disorders^10–13^. However, the relationship between altered sleep and the cognitive and behavioral aspects of FXS (or other ASDs) is unknown. *Fmr1* knockout mice recapitulate behavioral phenotypes seen in FXS patients, as well as disrupted sensory cortex plasticity and hippocampus-dependent memory consolidation^14–19^. However, while limited data using home-cage observational studies suggest alterations in sleep across development in *Fmr1^-/y^* mice^20, 21^, a full characterization of sleep behavior *Fmr1^-/y^* mice (e.g., using continuous polysomnographic recording) has not been done. Because sleep disruption can impair cognitive function and synaptic plasticity in both humans and mice^22–32^, one untested possibility is that disrupted sensory processing and cognition in FXS (and other ASDs) are exacerbated by abnormal sleep. If sleep disruption is a major driver of cognitive disruption in ASD, hypnotic treatment could prove to be an important therapeutic adjuvant for treatment^33–36^.

Recent studies have led to the development of new classes of hypnotic drugs - including orexin receptor antagonists and activators for G-protein inward rectifying potassium (GIRK) channels^37–39^. GIRK1/2-containing GIRK channels are highly expressed in regions such as the hippocampus, neocortex, and cerebellum ^40–42^, and can be directly activated by ML297, leading to neuronal hyperpolarization^43, 44^. Recent studies using ML297 in rodents have shown it suppresses seizure activity, acts as an anxiolytic, and increases NREM and REM sleep^39, 43–46^. Critically, GIRK1/2 channel activity is likely disrupted by loss of FMRP^47, 48^.

We used polysomnographic recordings to fully characterize alterations to sleep architecture and cortical EEG activity in adult *Fmr1^-/y^* mice. We find alterations to sleep amounts, architecture, and EEG oscillations, which are largely reversed by administration of the GIRK1/2 channel agonist ML297. ML297 administration following training on two hippocampus-mediated, sleep-dependent tasks (object location memory and contextual fear memory) led to a rescue of disrupted consolidation of memory in both tasks. These studies set the groundwork for understanding sleep as a therapeutic target for treating patients with FXS and other ASDs.

## Results

### *Fmr1^-/y^* mice have disrupted NREM sleep architecture and altered EEG activity

To fully characterize sleep phenotypes in a FXS mouse model, we first recorded EEG/EMG activity continuously across the rest (light) phase in *Fmr1^-/y^* mice and male wild-type (WT) littermates. 4-5 month old WT and *Fmr1^-/y^* mice were implanted with bilateral EEG electrodes over primary visual cortex (V1) and were recorded across the 12-h light phase (ZT0-12) (**Figure 1A**). *Fmr1^-/y^* mice spent significantly less total time in NREM sleep, more time in wake, and equal time in REM, compared to WT littermates (**Figure 1B**). Mean duration of NREM bouts was significantly shorter in *Fmr1^-/y^* mice (**Figure 1C**), and total number of NREM and wake bouts (but not REM bouts) was significantly greater in *Fmr1^-/y^* mice (**Figure 1D**) – suggesting fragmented NREM sleep architecture.

**Figure 1:**
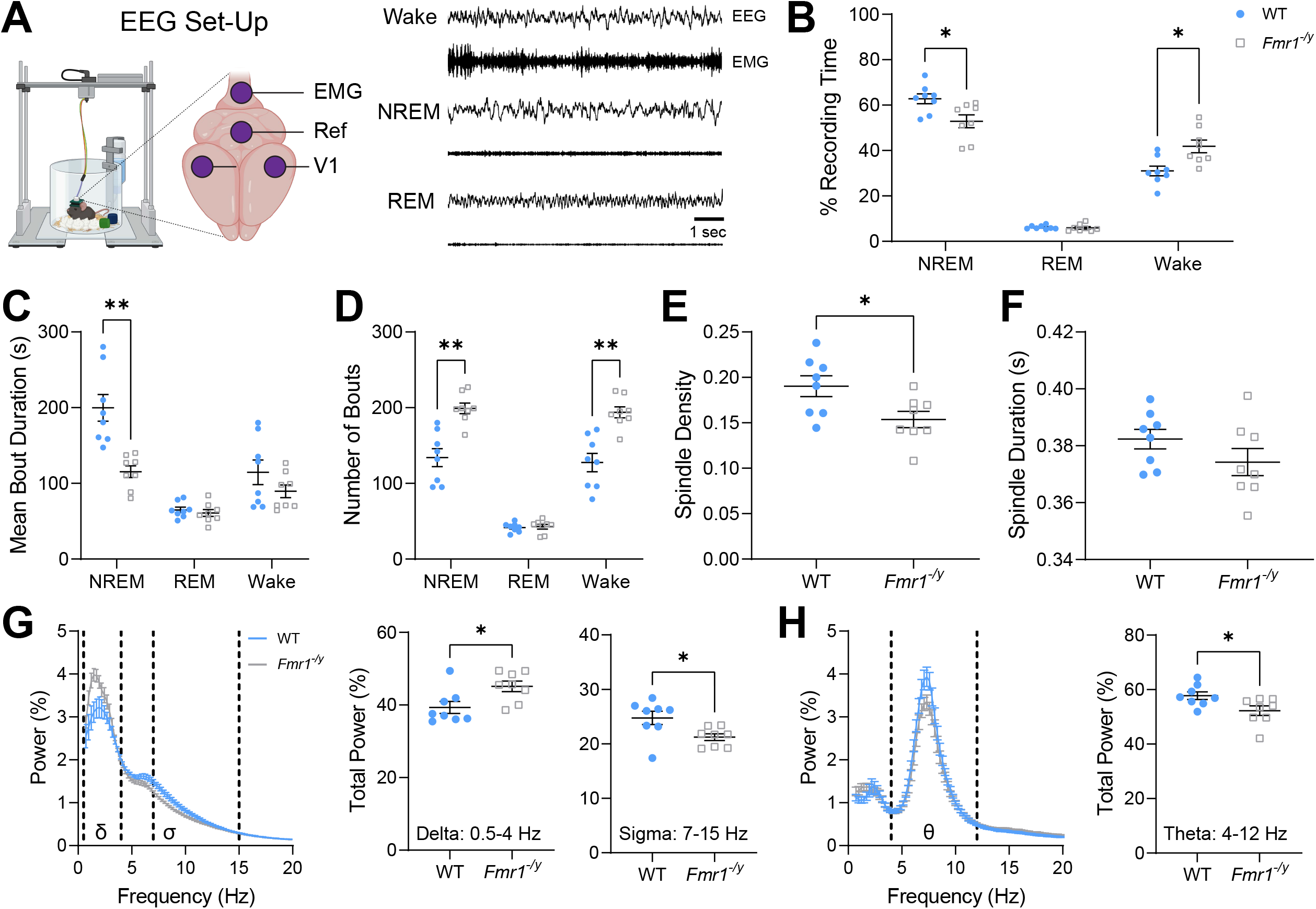
*Fmr1^-/y^*mice exhibit disrupted NREM sleep and altered EEG activity during their rest phase. **(A)** Schematic of EEG/EMG recording configuration, with EEG electrodes positioned over bilateral primary visual cortex (V1) and representative 10-sec epochs are shown for wake, NREM sleep, and REM sleep. **(B)** Percent recording time (for the entire 12-h light phase) spent in NREM sleep, REM sleep, and wake for WT and *Fmr1^-/y^* mice. Two-way RM ANOVA, *p*(state) < 0.0001, *p*(genotype) = 0.19, *p*(state x genotype interaction) = 0.0014. **(C)** Mean bout durations across the light phase for NREM, REM, and wake. Two-way RM ANOVA, *p*(state) < 0.0001, *p*(genotype) = 0.0062, *p*(state x genotype interaction) = 0.0003. **(D)** Number of light-phase bouts of NREM, REM, and wake. Two-way RM ANOVA, *p*(state) < 0.0001, *p*(genotype) = 0.0005, *p*(state x genotype interaction) < 0.0001. **(E)** NREM spindle density (detected events per second) during the 12-h light phase. **(F)** NREM spindle duration during the 12-h light phase. **(G)** NREM EEG power spectrum (***left***) with delta (δ; 0.5-4 Hz) and sigma (σ; 7-15 Hz) frequency bands highlighted via dashed lines. Two-way RM ANOVA, *p*(frequency) < 0.0001, *p*(genotype) = 0.60, *p*(frequency x genotype interaction) < 0.0001. Total NREM delta-band power (***middle***) and total sigma-band power for spindles (***right***) across the 12-h light phase. **(H)** REM EEG power spectrum (***left***), showing the theta (θ; 4-12 Hz) frequency band highlighted via dashed lines. Two-way RM ANOVA, *p*(frequency) < 0.0001, *p*(genotype) = 0.10, *p*(frequency x genotype interaction) < 0.0001. Total REM theta-band power (***right***) across the 12-h light phase. Sample size: *n* = 8 mice/genotype. *, **, and *** indicate *p* < 0.05, *p* < 0.01, and *p* < 0.001, Sidak’s *post hoc* test (**B-D**) or two-tailed, unpaired t-test (**E-H**). Data points and error bars indicate mean ± SEM. Symbols: filled (blue) circles and open (gray) squares = WT and *Fmr1^-/y^* mice, respectively.

To test whether other features of NREM sleep were disrupted in *Fmr1^-/y^* mice, we next compared EEG activity between the genotypes. Density of NREM sleep spindles (waxing-and-waning 7-15 Hz [sigma (σ) band] oscillations ^49–52^), detected in the EEG using a semi-automated method^46, 53^, was significantly decreased in *Fmr1^-/y^* mice compared to WT littermates (**Figure 1E**). Spindle duration was similar between genotypes (**Figure 1F**). With respect to overall state-specific EEG power, *Fmr1^-/y^* mice showed significantly increased delta (δ; 0.5-4 Hz) and decreased sigma (σ; 7-15 Hz) activity during NREM, and significantly reduced theta (θ; 4-12 Hz) power during REM, compared to WT littermates (**Figure 1G-H**). Consistent with previous findings from *Fmr1^-/y^* mice and rats ^54–56^, gamma (γ; 31-100 Hz) power was increased in the EEG during both NREM and wake in *Fmr1^-/y^* mice (**Figure S1A, C**). Delta power during REM and wake was similar between genotypes (**Figure S1B-C**). Thus, multiple state-specific cortical, thalamocortical, and hippocampal oscillations^57, 58^ are disrupted in *Fmr1^-/y^* mice, in association with disruptions in overall sleep architecture.

### Administration of hypnotic compound ML297 renormalizes NREM sleep in *Fmr1^-/y^* mice

Because activation of GIRK channels (via coupling to GABA_B_ receptors) is thought to be disrupted in FXS due to loss of FMRP^47, 48^, we tested whether GIRK1/2 agonist hypnotic ML297 could restore sleep in *Fmr1^-/y^* mice. We measured changes to sleep architecture and oscillations after administering ML297 or vehicle to *Fmr1^-/y^* mice and WT littermates. Mice implanted with EEG recording electrodes over V1 and prefrontal cortex (PFC) underwent 4-day, continuous EEG recording, consisting of a 24-h baseline (starting at ZT0) on day 1 (baseline A), a 24-h period following administration of vehicle (at ZT 0) on day 2, a second 24-h baseline recording (baseline B) on day 3, and a 24-h period following administration of ML297 (30 mg/kg) on day 4 (**Figure 2A**).

**Figure 2:**
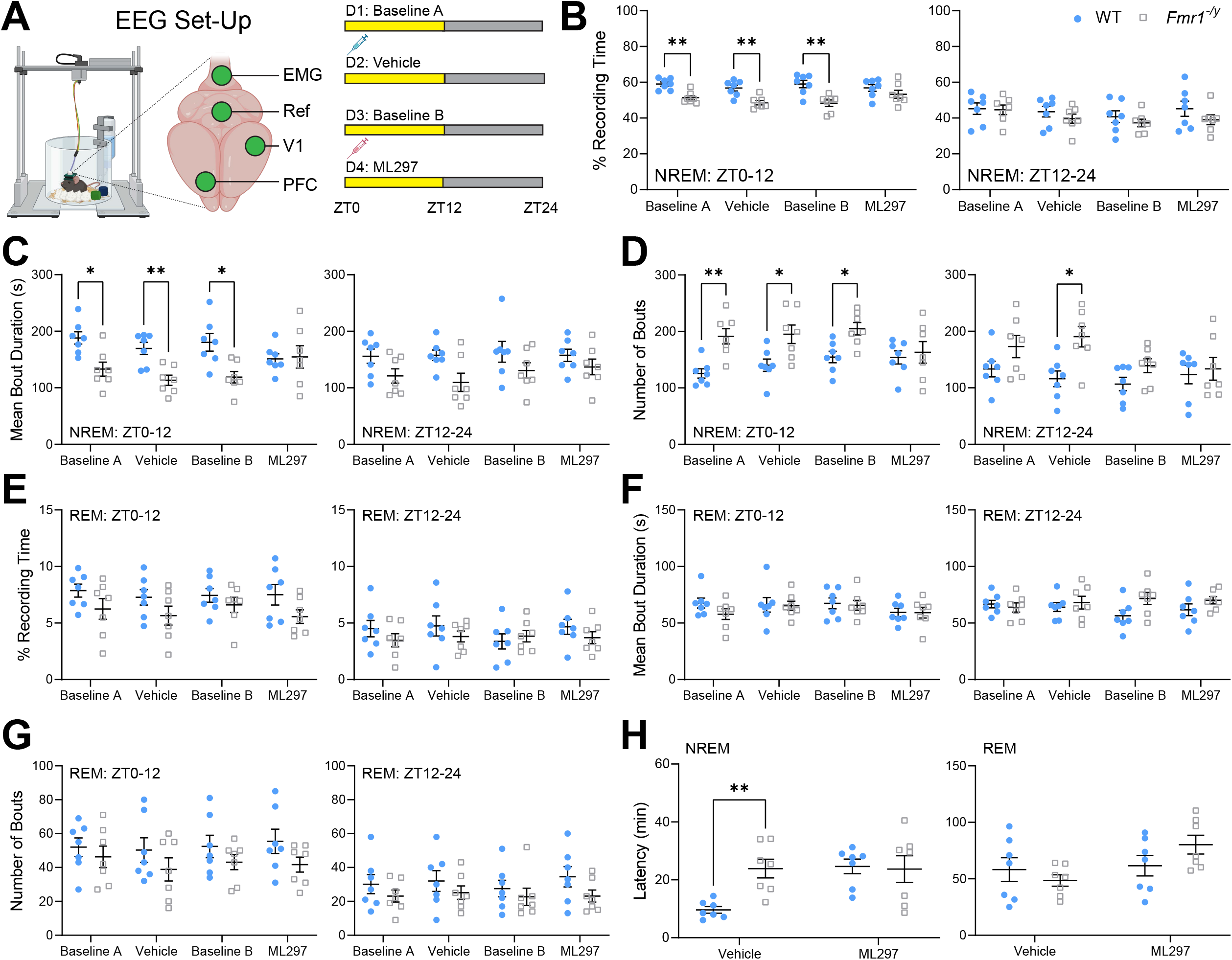
ML297 administration renormalizes rest-phase NREM sleep architecture in *Fmr1^-/y^* mice. **(A)** Schematic of EEG/EMG recording configuration, with EEG electrodes positioned over primary visual (V1) and prefrontal (PFC) cortical regions (***left***). Experimental timeline of 4-day continuous recording in a 12:12 light-dark cycle (***right***). Baseline recordings were carried out on days 1 and 3; vehicle and ML297 i.p. injections were given at lights-on (ZT0) on days 2 and 4, respectively. **(B)** NREM percent recording time for the 0-12 h light phase (***left***) and 12-24 h dark phase (***right***) across all four days for WT and *Fmr1^-/y^* mice. Two-way RM ANOVA for ZT0-12: *p*(condition) = 0.23, *p*(genotype) = 0.0009, *p*(condition x genotype interaction) = 0.083 and ZT12-24: *p*(condition) = 0.02, *p*(genotype) = 0.36, *p*(condition x genotype interaction) = 0.46. **(C)** NREM bout duration for the 0-12 h light phase (***left***) and 12-24 h dark phase (***right***) across all four days for WT and *Fmr1^-/y^* mice. Two-way RM ANOVA for ZT0-12: *p*(condition) = 0.25, *p*(genotype) = 0.0099, *p*(condition x genotype interaction) = 0.0033 and ZT12-24: *p*(condition) = 0.47, *p*(genotype) = 0.035, *p*(condition x genotype interaction) = 0.63. **(D)** Number of NREM bouts for the 0-12 h light phase (***left***) and 12-24 h dark phase (***right***) across all four days for WT and *Fmr1^-/y^*mice. Two-way RM ANOVA for ZT0-12: *p*(condition) = 0.09, *p*(genotype) = 0.0090, *p*(condition x genotype interaction) = 0.013 and ZT12-24: *p*(condition) = 0.018, *p*(genotype) = 0.059, *p*(condition x genotype interaction) = 0.029. **(E)** REM percent recording time for the 0-12 h light phase (***left***) and 12-24 h dark phase (***right***) across all four days for WT and *Fmr1^-/y^* mice. Two-way RM ANOVA for ZT0-12: *p*(condition) = 0.49, *p*(genotype) = 0.095, *p*(condition x genotype interaction) = 0.74 and ZT12-24: *p*(condition) = 0.089, *p*(genotype) = 0.48, *p*(condition x genotype interaction) = 0.021. **(F)** REM bout duration for the 0-12 h light phase (***left***) and 12-24 h dark phase (***right***) across all four days for WT and *Fmr1^-/y^* mice. Two-way RM ANOVA for ZT0-12: *p*(condition) = 0.30, *p*(genotype) = 0.49, *p*(condition x genotype interaction) = 0.62 and ZT12-24: *p*(condition) = 0.94, *p*(genotype) = 0.16, *p*(condition x genotype interaction) = 0.15. **(G)** Number of REM bouts for the 0-12 h light phase (***left***) and 12-24 h dark phase (***right***) across all four days for WT and *Fmr1^-/y^*mice. Two-way RM ANOVA for ZT0-12: *p*(condition) = 0.71, *p*(genotype) = 0.15, *p*(condition x genotype interaction) = 0.84 and ZT12-24: *p*(condition) = 0.41, *p*(genotype) = 0.25, *p*(condition x genotype interaction) = 0.63. **(H)** Latency to NREM (***left***) and REM (***right***) sleep after vehicle or ML297 treatment in WT and *Fmr1^-/y^*mice. Two-way RM ANOVA for NREM latency: *p*(treatment) = 0.0057, *p*(genotype) = 0.10, *p*(treatment x genotype interaction) = 0.005 and REM latency: *p*(treatment) = 0.10, *p*(genotype) = 0.52, *p*(treatment x genotype interaction) = 0.17. Sample size: *n* = 7 mice/genotype. * and ** indicate *p* < 0.05 and *p* < 0.01, Sidak’s *post hoc* test. Data points and error bars indicate mean ± SEM.

Similar to our initial findings, baseline recordings (baselines A and B) in *Fmr1^-/y^* mice were characterized by significantly decreased NREM across the 12-h light phase, while dark (i.e., active) phase (ZT12-24) NREM was similar between genotypes (**Figure 2B**). Disrupted NREM sleep architecture (decreased NREM bout durations and increased NREM bout numbers) was evident in *Fmr1^-/y^* mice across both baseline days’ light phase, but not dark phase (**Figure 2C-D; Figure S2A-C**). Vehicle treatment did not alter NREM sleep phenotypes in *Fmr1^-/y^* mice. However, ML297 administration restored NREM sleep time, bout duration, and bout number to levels seen in WT littermates (**Figure 2B-D**). REM sleep time and architecture was comparable between genotypes across both light and dark phases, and across baseline and treatment recording days (**Figure 2E-G**). Latency to NREM sleep was greater in *Fmr1^-/y^* mice after vehicle treatment, but remained comparable to WT littermates’ NREM latency after ML297 treatment (**Figure 2H**). Together, these data show that sleep disruption in *Fmr1^-/y^* mice is renormalized by ML297 to match sleep patterns WT littermates. This indicates that restoration of GIRK1/2 channel activation, which is thought to be disrupted in FXS ^47, 48^, can rescue NREM sleep deficits present in *Fmr1^-/y^* mice.

### Cortical subregion-specific NREM EEG changes and intracortical desynchrony are partially renormalized by ML297 in *Fmr1^-/y^* mice

Because we noted changes in state-specific EEG spectral power in *Fmr1^-/y^* mice, we next assessed whether these changes were time-of-day and brain region-specific, and how ML297 impacted cortical EEG activity. In V1, across both baseline recordings (baseline A and B), we again observed increased NREM delta (δ) and gamma (γ) power, and reduced sigma (σ) power, in *Fmr1^-/y^* mice – changes which were present across the light phase (**Figure 3A, C; Figure S3A**). In contrast, during the dark phase, V1 NREM delta power remained significantly elevated, while sigma and gamma power were comparable to WT littermates (**Figure S4A, C; Figure S3A**). In simultaneous EEG recordings in PFC, baseline NREM gamma power was elevated in *Fmr1^-/y^* mice during the light phase, but delta and sigma power were comparable between WT littermates and *Fmr1^-/y^* mice (**Figure 3B, D; Figure S3B**). Similar trends in PFC spectral power were observed across the dark cycle (**Figure S4B, D; Figure S3B**). These regional differences in NREM delta power are consistent with reported anteroposterior axis differences in NREM slow wave activity in children and adults with ASD^59, 60^. Similarly, baseline V1 recordings again showed decreased REM theta (θ) power in *Fmr1^-/y^* mice compared to WT littermates across the light and dark phase (**Figure 3E, G; Figure S4E, G**). However, no changes in REM theta were detectable in PFC at any time of day (**Figure 3F, H; Figure S4F, H**). This may reflect a similar brain region-specific change in the distribution of REM theta that has been reported in adult ASD patients^61^. In baseline wake, delta EEG power was similar between genotypes (in both cortical areas across both light and dark phases), while gamma power remained elevated across PFC and V1 in *Fmr1^-/y^* mice (**Figure S5A-D**).

**Figure 3:**
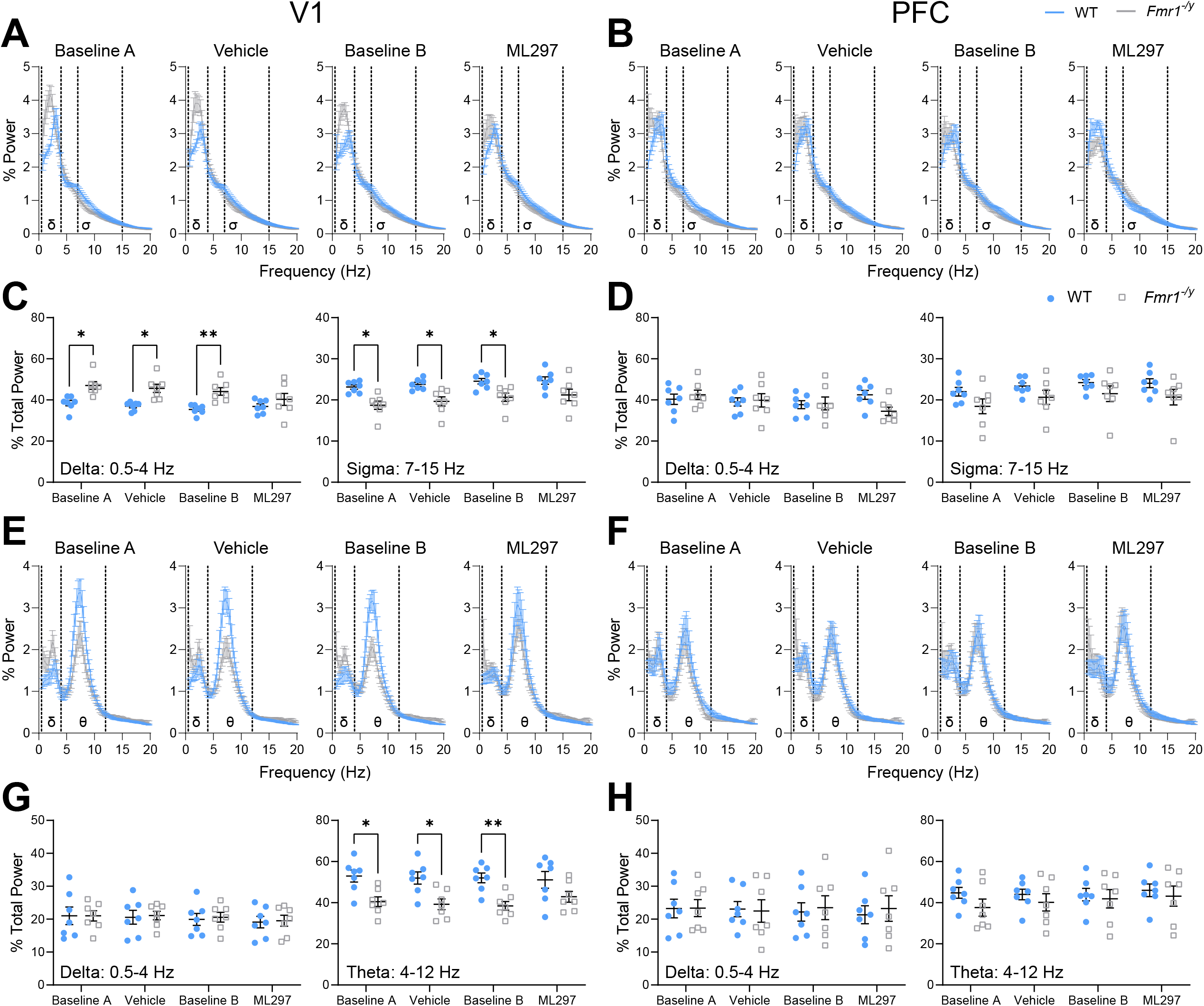
NREM and REM spectral power in V1, but not PFC, is altered in *Fmr1^-/y^* mice and normalized with ML297. (A) NREM EEG power spectra in V1 for baselines and treatment conditions during ZT0-12 (light phase) with delta (δ; 0.5-4 Hz) and sigma (σ; 7-15 Hz) frequency bands highlighted via dashed lines. Two-way RM ANOVA, *p*(frequency) < 0.0001, *p*(frequency x genotype interaction) < 0.0001, for baseline A-ML297 conditions and *p*(genotype) = 0.085 (baseline A), = 0.003 (vehicle), = 0.057 (baseline B), and = 0.42 (ML297). (B) NREM EEG power spectra in PFC for baselines and treatment conditions during ZT0-12 (light phase) with delta (δ; 0.5-4 Hz) and sigma (σ; 7-15 Hz) frequency bands highlighted via dashed lines. Two-way RM ANOVA, *p*(frequency) < 0.0001, *p*(frequency x genotype interaction) < 0.0001, for baseline A-ML297 conditions and *p*(genotype) = 0.088 (baseline A), = 0.21 (vehicle), = 0.26 (baseline B), and = 0.27 (ML297). (C) Total NREM delta-band power (***left***) and total sigma-band power for spindles (***right***) in V1 during ZT0-12. Two-way RM ANOVA for total delta power: *p*(condition) = 0.0009, *p*(genotype) = 0.0052, *p*(condition x genotype interaction) = 0.014 and total sigma power: *p*(condition) = 0.013, *p*(genotype) = 0.0049, *p*(condition x genotype interaction) = 0.83. (D) Total NREM delta-band power (***left***) and total sigma-band power for spindles (***right***) in PFC during ZT0-12. Two-way RM ANOVA for total delta power: *p*(condition) = 0.26, *p*(genotype) = 0.68, *p*(condition x genotype interaction) = 0.031 and total sigma power: *p*(condition) = 0.062, *p*(genotype) = 0.10, *p*(condition x genotype interaction) = 0.94. (E) REM EEG power spectra in V1 for baselines and treatment conditions during ZT0-12 (light phase) with delta (δ; 0.5-4 Hz) and theta (θ; 4-12 Hz) frequency bands highlighted via dashed lines. Two-way RM ANOVA, *p*(frequency) < 0.0001, *p*(frequency x genotype interaction) < 0.0001, for baseline A, vehicle, and baseline B conditions; ML297 condition, *p*(frequency) < 0.0001, *p*(frequency x genotype interaction) = 0.023 and *p*(genotype) = 0.083 (baseline A), = 0.051 (vehicle), = 0.046 (baseline B), and = 0.31 (ML297). (F) REM EEG power spectra in PFC for baselines and treatment conditions during ZT0-12 (light phase) with delta (δ; 0.5-4 Hz) and theta (θ; 4-12 Hz) frequency bands highlighted via dashed lines. Two-way RM ANOVA, *p*(frequency) < 0.0001, *p*(frequency x genotype interaction) > 0.99, for baseline A-ML297 conditions and *p*(genotype) = 0.12 (baseline A), = 0.23 (vehicle), = 0.73 (baseline B), and = 0.57 (ML297). (G) Total REM delta-band power (***left***) and total theta-band power (***right***) in V1 during ZT0-12. Two-way RM ANOVA for total delta power: *p*(condition) = 0.24, *p*(genotype) = 0.86, *p*(condition x genotype interaction) = 0.98 and total theta power: *p*(condition) = 0.54, *p*(genotype) = 0.0046, *p*(condition x genotype interaction) = 0.39. (H) Total REM delta-band power (***left***) and total theta-band power (***right***) in PFC during ZT0-12. Two-way RM ANOVA for total delta power: *p*(condition) = 0.66, *p*(genotype) = 0.87, *p*(condition x genotype interaction) = 0.56 and total theta power: *p*(condition) = 0.23, *p*(genotype) = 0.43, *p*(condition x genotype interaction) = 0.43. Sample size: *n* = 7 mice/genotype. * and ** indicate *p* < 0.05 and *p* < 0.01, Sidak’s *post hoc* test. Data points and error bars indicate mean ± SEM.

Vehicle treatment had no effects on any state-specific EEG features in *Fmr1^-/y^* mice (**Figure 3A, C, E, G; Figure S3A-B, Figure S4A, C, E, G; Figure S5B-D**). ML297 administration normalized V1 NREM delta and sigma power in *Fmr1^-/y^* mice (**Figure 3A, C**), but did not affect NREM gamma power, which remained elevated in both V1 and PFC (**Figure S3A-B**). Interestingly, however, ML297 did normalize wake gamma activity in both V1 and PFC regions – and this effect persisted across both light and dark phases (**Figure S5C-D**). REM theta power in V1 was also normalized in *Fmr1^-/y^* mice after ML297 treatment (**Figure 3E, G; Figure S4E, G**).

Observed region-specific changes in NREM sigma power and sleep spindles in *Fmr1^-/y^* mice suggest that coordination of these thalamocortical oscillations^62^ might be disrupted by loss of FMRP. Because synchronous thalamocortical activity during sleep spindles is thought to be essential for brain development and cognitive function^53, 58, 63–69^, we tested whether NREM spindle density, or spindle coordination between brain regions, could be altered by ML297 administration. In baseline recordings from V1, we again observed reduced spindle density *Fmr1^-/y^* mice compared to WT littermates during the light phase, but not dark phase (**Figure 4A**). In contrast, spindle density in PFC recordings did not differ significantly between genotypes (**Figure 4B**). Vehicle administration did not affect V1 NREM spindle density in *Fmr1^-/y^* mice, but caused an apparent *decrease* in spindle density in PFC. In contrast, ML297 restored V1 spindle density to WT levels in *Fmr1^-/y^* mice, and had no effect on spindle density in PFC (**Figure 4A-B**).

**Figure 4:**
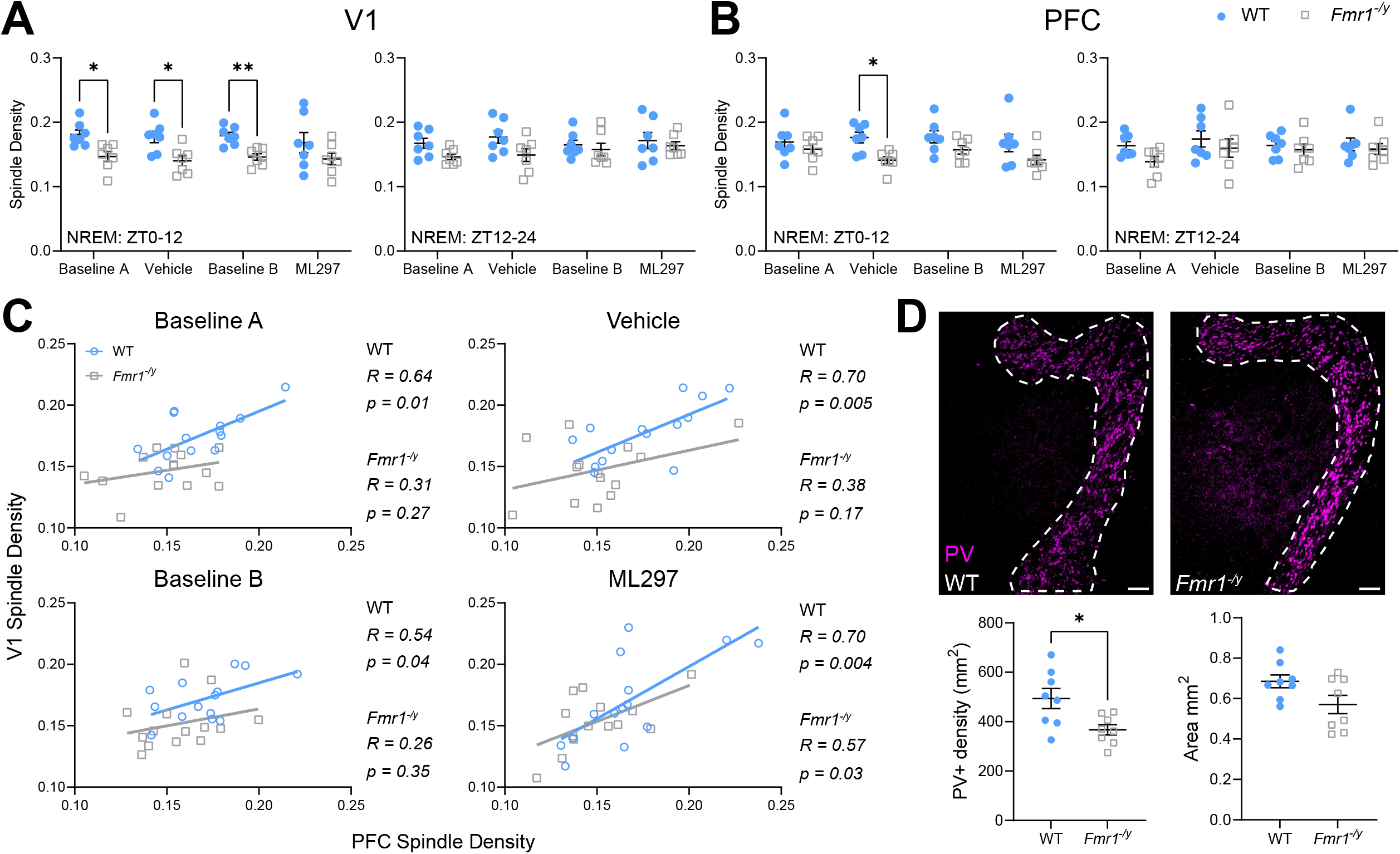
*Fmr1^-/y^*mice show discrepant NREM spindle densities between V1 and PFC due to reduced V1 spindling that is rescued by ML297 administration. **(A)** Spindle density (detected events per second) during NREM sleep for ZT0-12 (***left***) and ZT12-24 (***right***) in V1. Two-way RM ANOVA for ZT0-12: *p*(condition) = 0.63, *p*(genotype) = 0.0019, *p*(condition x genotype interaction) = 0.89 and ZT12-24: *p*(condition) = 0.61, *p*(genotype) = 0.04, *p*(condition x genotype interaction) = 0.53. **(B)** Spindle density (detected events per second) during NREM sleep for ZT0-12 (***left***) and ZT12-24 (***right***) in PFC. Two-way RM ANOVA for ZT0-12: *p*(condition) = 0.38, *p*(genotype) = 0.012, *p*(condition x genotype interaction) = 0.46 and ZT12-24: *p*(condition) = 0.31, *p*(genotype) = 0.18, *p*(condition x genotype interaction) = 0.68. **(C)** Correlations of V1 and PFC spindle densities across the four continuous days of recording between WT and *Fmr1^-/y^* mice. Data points represent NREM spindle density across either the light or dark phase (ZT0-12 or ZT12-24) for each individual mouse in both WT and *Fmr1^-/y^* groups. *R* and *p* values for Pearson correlation of each data set are found alongside corresponding graph. **(D)** Representative images of parvalbumin (PV+) expressing interneurons in the thalamic reticular nucleus (TRN) (***top***) outlined by dashed lines in WT and *Fmr1^-/y^* mice. Scale bar = 250 µm. PV+ interneuron density of the TRN (***left***) and area of TRN measured (***right***). Sample size: *n* = 7 mice/genotype (panels **A-C**) and *n* = 8 (panel **D**). * and ** indicate *p* < 0.05 and *p* < 0.01, Sidak’s *post hoc* test (panels **A-C**) and two-tailed, unpaired t-test (panel **D**). Data points and error bars indicate mean ± SEM.

Because these changes were brain region-specific (similar to other changes in NREM oscillations in *Fmr1^-/y^* mice), we compared NREM spindle densities for simultaneous recordings in V1 vs. PFC of individual mice. At baseline, spindle densities between the two regions were consistently correlated in WT littermates, but not in *Fmr1^-/y^* mice (**Figure 4C**). When WT littermates were treated with either vehicle or ML297, the consistent positive relationship between V1 and PFC spindle densities was unaffected. Vehicle administration had no effect in *Fmr1^-/y^* mice (i.e., V1 and PFC spindle densities remained uncorrelated). However, after ML297 treatment, spindle densities became significantly correlated between V1 and PFC EEG sites in individual mice, suggesting better coordination of these oscillations during NREM sleep (**Figure 4C**). The observed changes in spindle coordination prompted us to assess whether structural changes were present in the thalamic reticular nucleus (TRN) (a structure known to be essential for generation of NREM spindles^63^) of *Fmr1^-/y^* mice. Using quantitative immunohistochemistry (IHC), we found a reduced density of parvalbumin-expressing (PV+) interneurons in the TRN of *Fmr1^-/y^* mice compared to WT littermates (**Figure 4D**). This suggests that ML297 restores coordination of spindle oscillations that is lost in *Fmr1^-/y^* mice due to microcircuit-level changes in the TRN.

To further quantify state-specific EEG oscillatory synchrony, we compared overall V1-PFC spectral coherence for *Fmr1^-/y^* mice and WT littermates. Across both baseline recordings (baseline A and B), we found NREM V1-PFC delta band coherence was significantly lower in *Fmr1^-/y^* mice compared to WT littermates (**Figure 5A, C**). Sigma band coherence was also significantly reduced during the first baseline (baseline A) recording session (**Figure 5C**). NREM V1-PFC gamma coherence (**Figure S6A-B**) and REM delta and theta coherence (**Figure 5B, D**) was similar between genotypes. Vehicle treatment did not affect these baseline differences in coherence. In contrast, ML297 administration normalized V1-PFC NREM delta coherence (**Figure 5A-D**). Together, our data suggests that coordination of NREM delta and spindle oscillations, which is disrupted in *Fmr1^-/y^* mice, can be restored by administration of ML297.

**Figure 5:**
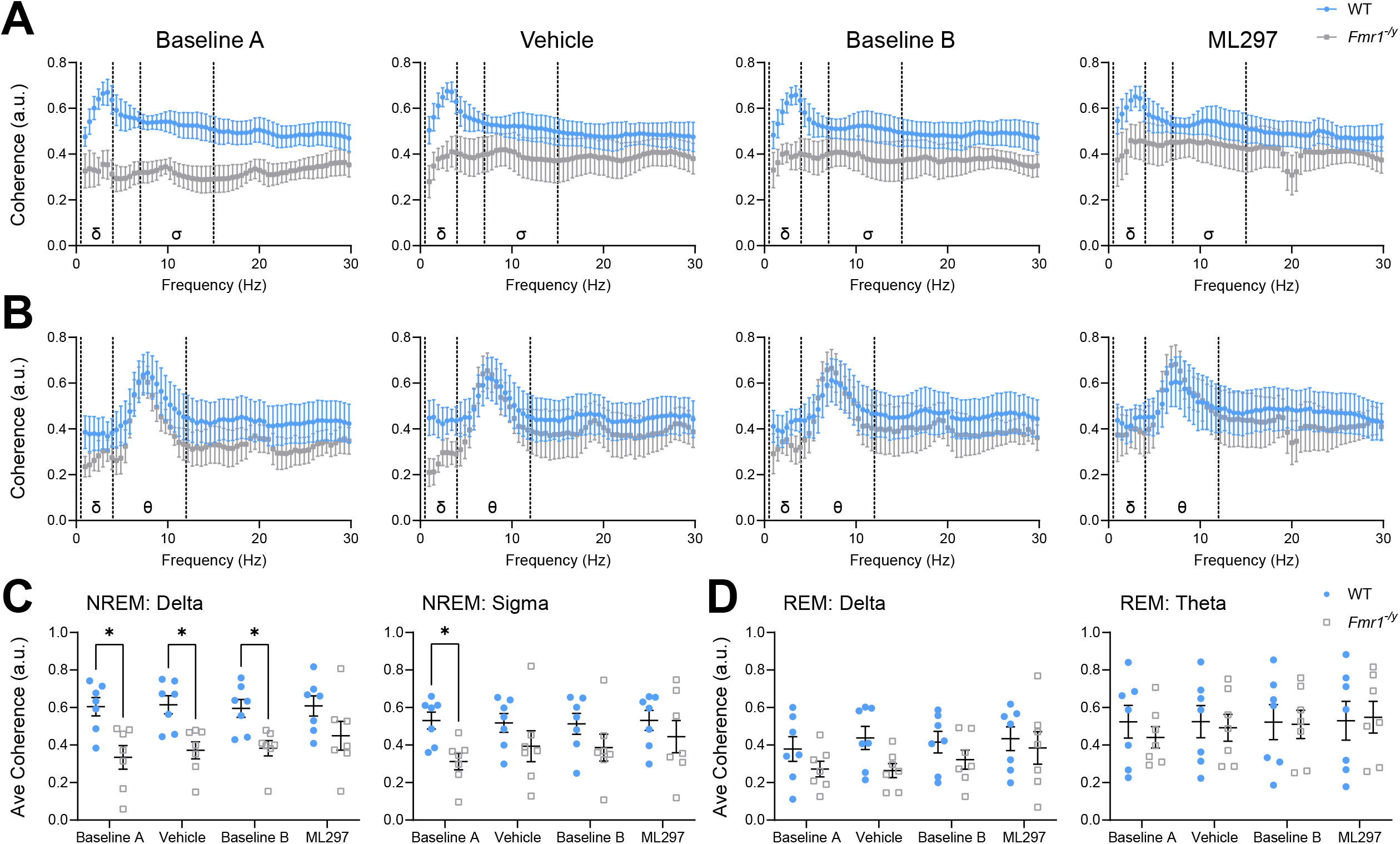
V1-PFC coherence during NREM sleep is altered in *Fmr1^-/y^* mice. **(A)** NREM-associated V1-PFC spectral coherence across baseline and treatment sessions. Two-way ANOVA, *p*(frequency) > 0.99, *p*(genotype) < 0.0001, *p*(frequency x genotype interaction) > 0.99, for baseline A-ML297 conditions. Delta (δ = 0.5-4 Hz) and sigma (σ = 7-15 Hz) frequency bands are denoted with their symbol and divided by dashed lines. **(B)** REM-associated V1-PFC spectral coherence across baseline and treatment sessions. Two-way ANOVA, *p*(frequency) = 0.0003, *p*(genotype) < 0.0001, *p*(frequency x genotype interaction) > 0.99 for baseline A. Two-way ANOVA, *p*(frequency) = 0.078, *p*(genotype) < 0.0001, *p*(frequency x genotype interaction) > 0.99 for vehicle. Two-way ANOVA, *p*(frequency) = 0.26, *p*(genotype) = 0.0004, *p*(frequency x genotype interaction) > 0.99 for baseline B. Two-way ANOVA, *p*(frequency) = 0.85, *p*(genotype) = 0.044, *p*(frequency x genotype interaction) > 0.99 for ML297. Delta (δ = 0.5-4 Hz) and theta (θ = 4-12 Hz) frequency bands are denoted with their symbol and divided by dashed lines. **(C)** Average NREM V1-PFC delta-band coherence (***left***) and sigma-band coherence (***right***). Two-way RM ANOVA for delta coherence: *p*(condition) = 0.39, *p*(genotype) = 0.0042, *p*(condition x genotype interaction) = 0.45 and sigma coherence: *p*(condition) = 0.23, *p*(genotype) = 0.11, *p*(condition x genotype interaction) = 0.20. **(D)** Average REM V1-PFC delta-band coherence (***left***) and theta-band coherence (***right***). Two-way RM ANOVA for delta coherence: *p*(condition) = 0.10, *p*(genotype) = 0.19, *p*(condition x genotype interaction) = 0.26 and theta coherence: *p*(condition) = 0.23, *p*(genotype) = 0.81, *p*(condition x genotype interaction) = 0.28. Sample size: *n* = 7 mice/genotype. * indicates *p* < 0.05, Sidak’s *post hoc* test. Data points and error bars indicate mean ± SEM.

### Post-learning ML297 administration rescues deficits in sleep-dependent memory consolidation in *Fmr1^-/y^* mice

In mice, consolidation of hippocampus-dependent contextual fear memory (CFM) and object location memory (OLM) are disrupted by brief sleep deprivation (SD) in the hours immediately following training^27, 29, 31, 32, 46, 70–72^. *Fmr1^-/y^* mice have deficits in both CFM ^15, 73, 74^ and OLM^75^, but it is unclear whether the sleep disruptions we observe in *Fmr1^-/y^* mice could mediate deficits in consolidation for these tasks. We tested whether restoring sleep with ML297 could rescue sleep-dependent memory consolidation in *Fmr1^-/y^* mice. We first carried out single-trial conditioning (CFC; placement in a novel context, paired with a foot shock) in *Fmr1^-/y^* mice and WT littermates. The two groups showed similar, very low freezing prior to shock during CFC. However, similar to prior reports^15, 73, 74^, *Fmr1^-/y^* mice showed reduced freezing during CFM recall (when returned to the shock context) 24 h later (**Figure 6A-B**).

**Figure 6:**
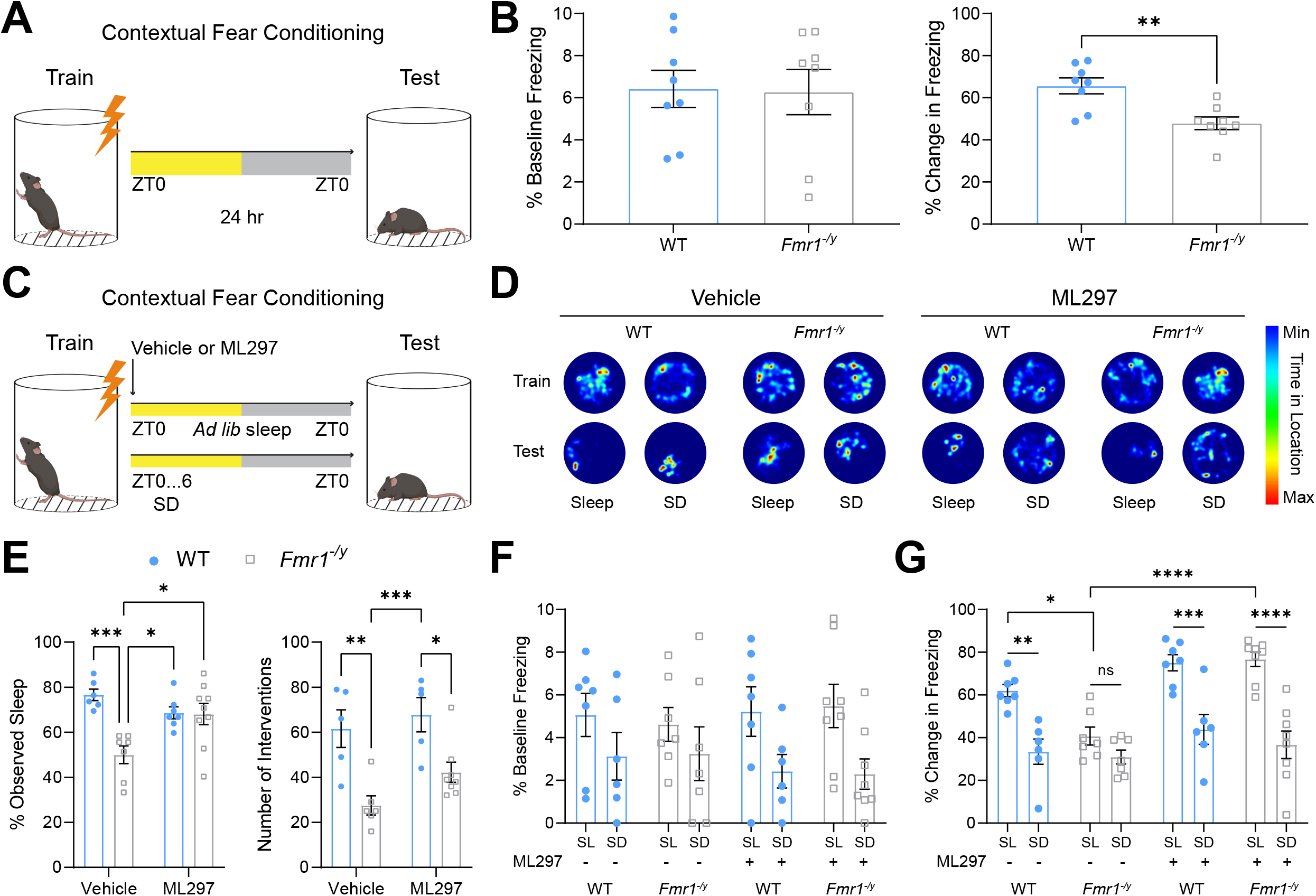
Contextual fear memory (CFM) consolidation is rescued in a sleep-dependent manner by post-conditioning ML297 administration in *Fmr1^-/y^* mice. **(A)** WT and *Fmr1^-/y^* mice underwent single-trial CFC at ZT0, and CFM testing 24 h later. **(B)** Baseline (pre-shock) freezing behavior during CFC training (***left***) between WT and *Fmr1^-/y^*mice. CFM (measured as increased freezing from pre-shock baseline during testing; ***right***) between WT and *Fmr1^-/y^* mice. Sample size: *n* = 8 mice/genotype. ** indicates *p* < 0.01, two-tailed, unpaired t-test. Data shown as mean ± SEM. **(C)** WT and *Fmr1^-/y^* mice underwent single-trial CFC at ZT0, followed immediately by either vehicle or ML297 administration and either *ad lib* sleep or 6-hour sleep deprivation (SD), and CFM testing 24 h later. **(D)** Representative heat maps showing total time-in-location for mice in the conditioning chamber during CFC (pre-shock) training and CFM testing. **(E)** Total amounts of visually observed sleep (***left***) and number of interventions during sleep deprivation (SD) (***right***) during the first 6 hours following CFC. For observed sleep, two-way ANOVA, *p*(treatment) = 0.21, *p*(genotype) = 0.0019, *p*(treatment x genotype) = 0.0028. For sleep interventions during SD, two-way ANOVA, *p*(treatment) = 0.098, *p*(genotype) < 0.0001, *p*(treatment x genotype) = 0.49. **(F)** Baseline (pre-shock) freezing behavior during CFC between WT and *Fmr1^-/y^* mice across both treatments and sleep conditions. Three-way ANOVA, *p*(treatment) = 0.82, *p*(genotype) = 0.94, *p*(sleep condition) = 0.0019, *p*(treatment x sleep condition) = 0.35, *p*(genotype x sleep condition) = 0.95, *p*(treatment x genotype) = 0.87, *p*(treatment x genotype x sleep condition) = 0.73. **(G)** Freezing behavior during CFM testing between WT and *Fmr1^-/y^* mice across both treatments and sleep conditions. Three-way ANOVA, *p*(treatment) < 0.0001, *p*(genotype) = 0.03, *p*(sleep condition) < 0.0001, *p*(treatment x sleep condition) = 0.019, *p*(genotype x sleep condition) = 0.47, *p*(treatment x genotype) = 0.18, *p*(treatment x genotype x sleep condition) = 0.047. For panels **E-G**, sample sizes for the sleep group are: *n* = 7 (WT+vehicle), *n* = 7 (*Fmr1^-/y^*+vehicle), *n* = 7 (WT+ML297), *n* = 8 (*Fmr1^-/y^*+ML297) and for the SD group are: *n* = 6 (WT+vehicle), *n* = 7 (*Fmr1^-/y^*+vehicle), *n* = 6 (WT+ML297), *n* = 8 (*Fmr1^-/y^*+ML297). *, **, ***, and **** indicate *p* < 0.05, *p* < 0.01, *p* < 0.001, *p* < 0.0001; Sidak’s *post hoc* test. Data shown as mean ± SEM. SL = sleep; SD = sleep deprived. For ML297, + indicates presence of compound, - indicates vehicle control and no ML297.

We next tested whether normalizing post-CFC sleep with ML297 could improve CFM consolidation in *Fmr1^-/y^* mice. Mice were administered either vehicle or ML297 immediately following CFC. To test for sleep-dependence of drug administration, half of the mice within each treatment group were allowed *ad lib* sleep (SL), while the second half underwent 6 h of sleep deprivation (SD) via gentle handling in their home cage (**Figure 6C**). Mice allowed *ad lib* sleep were visually monitored over the first 6 h post-CFC, to assess drug effects on sleep behavior. Consistent with our polysomnographic data, vehicle-treated *Fmr1^-/y^* mice had significantly reduced post-CFC sleep time compared to those given ML297, and compared to WT littermates with either treatment (**Figure 6E**). In addition, consistent with the interpretation that *Fmr1^-/y^* mice have reduced overall sleep drive, the number of interventions required to prevent sleep during the 6-h SD period was significantly lower in *Fmr1^-/y^* mice, regardless of treatment (**Figure 6E**). Pre-shock freezing was similarly low between the two genotypes during CFC training (**Figure 6D, F**). Vehicle-treated SL *Fmr1^-/y^* mice showed significantly reduced freezing during CFM testing compared with ML297-treated SL mice, and with SL WT littermates. Critically, CFM performance in ML297-treated *Fmr1^-/y^* SL mice was similar to that of WT SL mice, indicating a functional rescue of CFM consolidation by ML297 (**Figure 6D, G**). *Fmr1^-/y^* mice and WT littermates administered ML297 showed deficits in CFM after post-CFC SD, consistent with previous reports of SD’s disruptive effects on CFM^27, 32, 46^. This suggests that rescue of CFM consolidation in *Fmr1^-/y^* mice is sleep-dependent. For vehicle-treated WT mice, SD caused a disruption to CFM consolidation that was similar to that seen in *Fmr1^-/y^* mice (**Figure 6D, G**).

We next tested whether ML297 could rescue consolidation of other forms of hippocampus-mediated, sleep-dependent memory in *Fmr1^-/y^* mice. OLM is a form of spatial memory known to be disrupted in *Fmr1^-/y^* mice^75^. On the day prior to training on the OLM task (**Figure 7A**), mice were exposed to the OLM arena for 5 min, during which open field (OF) activity was used to measure locomotor activity and anxiety-related behavior (i.e., thigmotaxis). During the OF test (habituation period), *Fmr1^-/y^* mice and WT littermates showed similar locomotion (total travel distance) and relative time spent in outer and inner zones of the arena (**Figure 7B**). Mice underwent OLM training at ZT0, during which they explored two identical objects placed inside the arena for 10 min (**Figure 7A**). 24 h later, mice were returned to the arena where one of the two objects had been displaced and the other kept in the same (familiar) location. A discrimination index, used to measure selective interaction with the displaced vs. familiar objects, was calculated for each mouse. Consistent with prior findings^75^, discrimination indices were significantly lower during OLM testing for *Fmr1^-/y^* mice compared to WT littermates (**Figure 7C**). To test whether ML297 can rescue OLM consolidation in *Fmr1^-/y^* mice, a second cohort of mice were administered either vehicle or ML297 immediately after OLM training (**Figure 7D**). While locomotion and thigmotaxis were again comparable during habituation between genotypes (**Figure 7E**), performance during OLM testing varied by both genotype and treatment. *Fmr1^-/y^* mice treated with ML297 showed a significant improvement in discrimination between displaced and familiar object locations compared to *Fmr1^-/y^* mice treated with vehicle, suggesting at least partial rescue of OLM consolidation. However, WT littermates treated with ML297 showed no improvement in OLM consolidation, and in fact performed worse than vehicle-treated counterparts (**Figure 7F**). Together, these data indicate that restoration of post-learning sleep in *Fmr1^-/y^* mice with ML297 rescues both CFM and OLM consolidation, and further suggest that loss of sleep could contribute to cognitive impairments associated with FXS.

**Figure 7:**
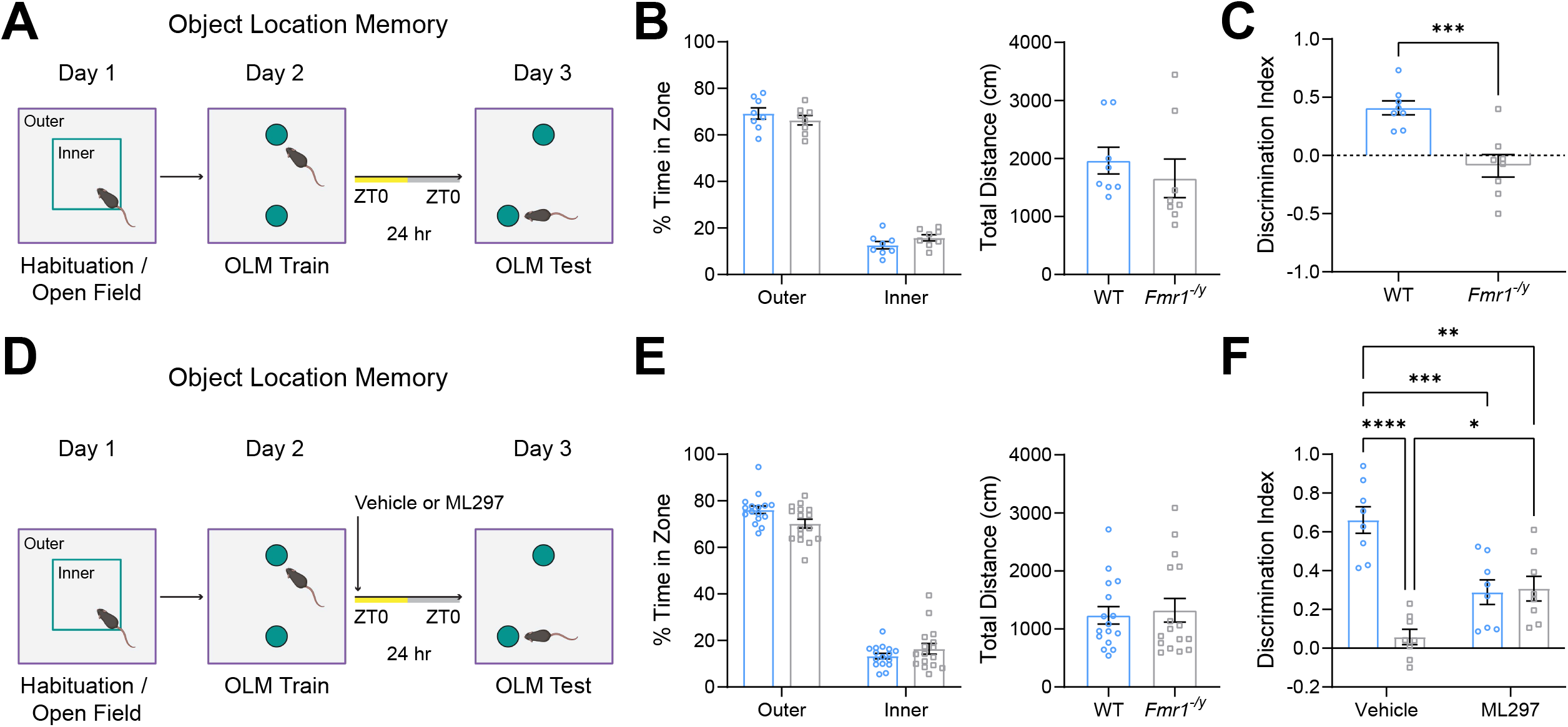
GIRK channel activation improves sleep-dependent object location memory consolidation in *Fmr1^-/y^*mice. **(A)** WT and *Fmr1^-/y^*mice (*n* = 8/genotype) underwent habituation to the object location memory (OLM) arena (combined with open field [OF] behavioral measurement) at ZT0. 24 h later, mice underwent OLM training with two identical objects in the arena. After an additional 24 h, mice were tested for OLM in the same arena with one object moved to a new location. **(B)** Proportion of time spent in inner and outer zones of the arena (***left***) between WT and *Fmr1^-/y^* mice during habituation. Two-way ANOVA, *p*(field zone) < 0.0001, *p*(genotype) = 0.94, *p*(field zone x genotype interaction) = 0.12. Total distance traveled during habituation (***right***) between genotypes. Two-tailed, unpaired t-test. Data shown as mean ± SEM. **(C)** Index for WT and *Fmr1^-/y^*mice to discriminate between displaced and non-displaced objects during OLM testing. *** indicates *p* < 0.001, two-tailed, unpaired t-test. Data shown as mean ± SEM. **(D)** WT and *Fmr1^-/y^* mice underwent habituation to the OLM arena and OLM training as described above, followed immediately by ML297 or vehicle administration and *ad lib* sleep. After an additional 24 h, mice were tested for OLM in the same arena with one object moved to a new location. **(E)** Open field activity (***left***) between WT and *Fmr1^-/y^*mice during habituation. Two-way ANOVA, *p*(field zone) < 0.0001, *p*(genotype) = 0.43, *p*(field zone x genotype interaction) = 0.013. Total distance traveled (***right***) was also similar between genotypes. Two-tailed, unpaired t-test. Data shown as mean ± SEM. **(F)** Index for WT and *Fmr1^-/y^*mice to discriminate between displaced and non-displaced objects during OLM testing with either vehicle or ML297 treatment. *n* = 8 mice/experimental condition. Two-way ANOVA, *p*(treatment) = 0.31, *p*(genotype) < 0.0001, *p*(treatment x genotype interaction) < 0.0001. *, **, ***, and **** indicate *p* < 0.05, *p* < 0.01, *p* < 0.001, *p* < 0.0001; Sidak’s *post hoc* test. Data shown as mean ± SEM.

### Memory consolidation improvements in *Fmr1^-/y^* mice are associated with ML297-mediated normalization of hippocampal activation patterns during recall

During both learning and subsequent memory retrieval, immediate early gene (IEG) expression among hippocampal dentate gyrus (DG) granule cells, and DG “engram neuron” re-activation plays a causal role in memory recall^76^. We tested whether CFM deficits in *Fmr1^-/y^* mice changes are associated with altered DG activity patterns during CFM recall, and whether ML297 has an impact on these activity patterns. *Fmr1^-/y^* mice and WT littermates underwent CFC followed by ML297 or vehicle administration. Mice were returned to their home cages after 24-h CFM testing, and perfused 90 min later to quantify cFos expression patterns associated with memory recall (**Figure 8A**). Across the entire DG region, vehicle-treated *Fmr1^-/y^* mice had significantly greater cFos+ neuron density after recall compared to WT littermate counterparts. This increase was reversed in *Fmr1^-/y^* mice treated with ML297, where cFos+ neuron density was normalized to levels observed in the DG of WT-vehicle mice. In contrast, WT littermates ML297 showed significant *increase* in DG cFos+ neuron density compared to WT-vehicle mice (**Figure 8B**), consistent with recent findings^77^. Overall density of cFos+ neurons in DG predicted successful CFM, although the direction of this relationship differed between *Fmr1^-/y^* mice and WT littermates. In WT littermates, higher density of cFos+ neurons corresponded to increased freezing behavior. *Fmr1^-/y^* mice showed the opposite relationship, where lower density of cFos+ neurons corresponded to improved CFM (**Figure 8D**). Our data suggest that overactivation of the DG network in *Fmr1^-/y^* mice disrupts CFM recall, and normalization of DG activity to WT levels with post-CFM ML297 administration improves CFM recall.

**Figure 8:**
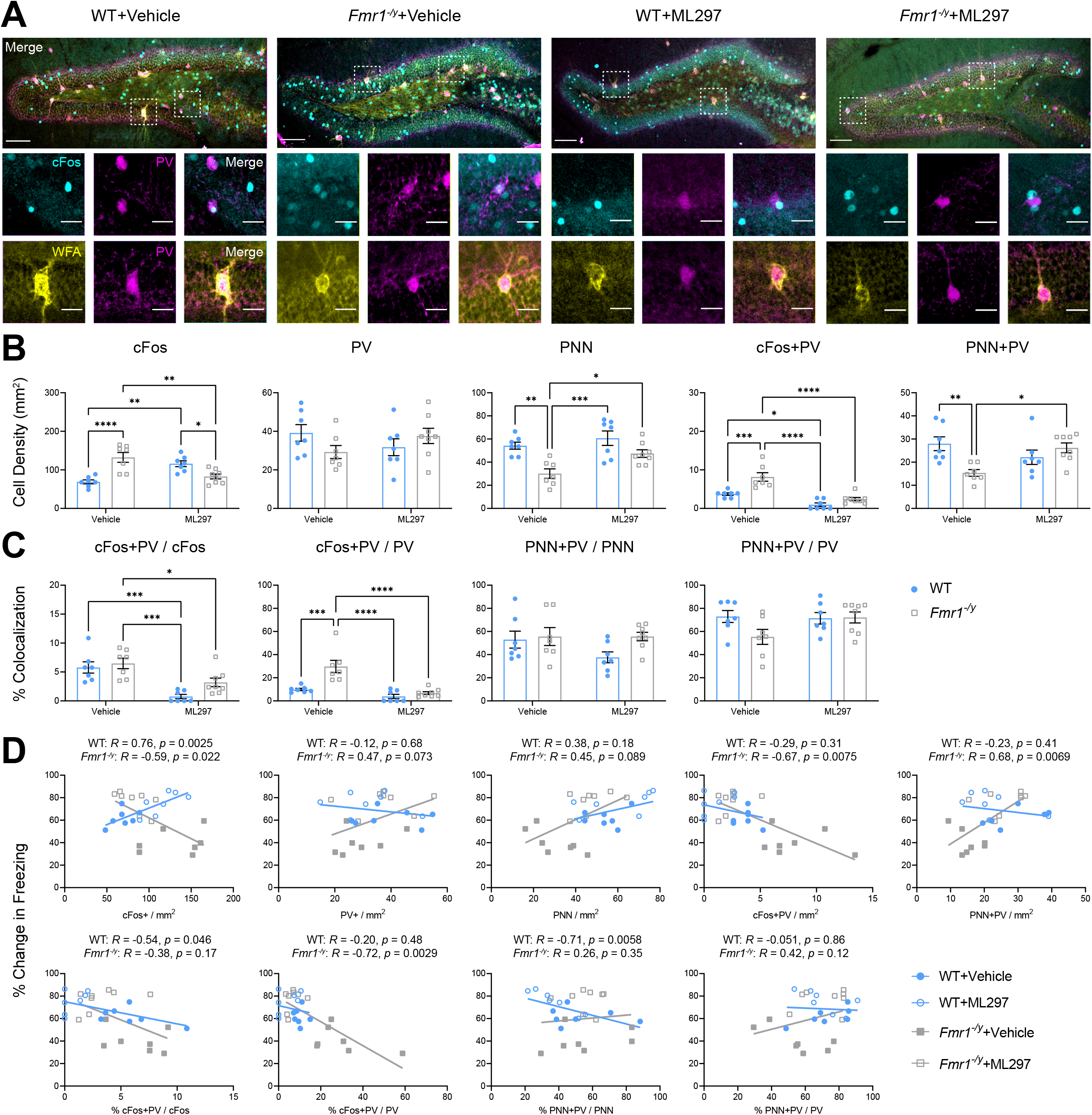
*Fmr1^-/y^* mice show increased DG network activity during memory recall, which is normalized by ML297 administration CFM consolidation. **(A)** Vehicle- and ML297-treated WT and *Fmr1^-/y^*mice with *ad lib* sleep were perfused 90 min following CFM recall to assess cFos expression, parvalbumin (PV) interneuron density, and perineuronal net (PNN) activity. Representative recall-associated cFos (cyan), PV (magenta), and wisteria floribunda agglutinin (WFA; yellow) for PNN expression in DG for the four treatment groups. Scale bar = 100 µm (***top***, whole DG); scale bar = 30 µm (***bottom***, inset of select neurons). **(B)** Quantification of DG cFos+, PV+, PNN, and co-localization neuronal density. Two-way ANOVA, for cFos+ neurons: *p*(treatment) = 0.88, *p*(genotype) = 0.082, *p*(treatment x genotype interaction) < 0.0001. For PV+ neurons: *p*(treatment) = 0.90, *p*(genotype) = 0.62, *p*(treatment x genotype interaction) = 0.06. For PNN: *p*(treatment) = 0.01, *p*(genotype) = 0.0002, *p*(treatment x genotype interaction) = 0.22. For cFos+PV co-localized neurons: *p*(treatment) < 0.0001, *p*(genotype) = 0.0001, *p*(treatment x genotype interaction) = 0.031. For PNN+PV co-localized neurons: *p*(treatment) = 0.32, *p*(genotype) = 0.098, *p*(treatment x genotype interaction) = 0.0026. **(C)** Quantification of DG percent co-localization of cFos+, PV+, and PNN over total neuronal populations. Two-way ANOVA, for % cFos+PV cells over total cFos: *p*(treatment) < 0.0001, *p*(genotype) = 0.056, *p*(treatment x genotype interaction) = 0.28. For % cFos+PV cells over total PV: *p*(treatment) < 0.0001, *p*(genotype) = 0.0005, *p*(treatment x genotype interaction) = 0.0061. For % PNN+PV cells over total PNN: *p*(treatment) = 0.21, *p*(genotype) = 0.096, *p*(treatment x genotype interaction) = 0.21. For % PNN+PV cells over total PV: *p*(treatment) = 0.16, *p*(genotype) = 0.13, *p*(treatment x genotype interaction) = 0.10. **(D)** Pearson correlation of cFos+, PV+, and PNN neuron densities and associated co-localizations in the entire DG vs. associated change in freezing behavior during CFM recall. *R* and *p* values for Pearson correlation of each data set are found alongside corresponding graph. Sample sizes: *n* = 7 (WT+vehicle), *n* = 7 (*Fmr1^-/y^*+vehicle), *n* = 7 (WT+ML297), *n* = 8 (*Fmr1^-/y^*+ML297). *, **, ***, and **** indicate *p* < 0.05, *p* < 0.01, *p* < 0.001, *p* < 0.0001; Tukey’s *post hoc* test. Data shown as mean ± SEM.

Alterations in excitatory-inhibitory (E-I) balance have been proposed as a mechanism promoting hyperactivity of neural circuits *Fmr1^-/y^* mice^78, 79^. To test whether changes in inhibitory DG networks are associated with disrupted CFM recall, we also measured parvalbumin (PV+) interneuron density and perineuronal net (PNN) expression in the DG following recall. PV+ interneuron density did not differ between the two genotypes, regardless of treatment. However, density of DG PNNs was reduced in vehicle-treated *Fmr1^-/y^* mice. PNN density in ML297-treated *Fmr1^-/y^* mice was restored to levels observed in WT littermates (**Figure 8B**). However, no significant correlations were found between CFM recall and density of PV+ interneurons or PNNs for either genotype (**Figure 8D**). We also assessed the expression of cFos among PV+ interneurons during CFM recall, as a proxy measure of their activity level. Similar to overall cFos+ neuron density, the density of cFos+PV neurons was significantly increased in vehicle-treated *Fmr1^-/y^* mice compared with WT littermates (in either treatment group; **Figure 8B**). Similarly, the proportion of all PV+ interneurons expressing cFos (cFos+PV/PV) was significantly higher in vehicle-treated *Fmr1^-/y^* mice, compared with WT littermates (**Figure 8B**). ML297 administration to *Fmr1^-/y^* mice normalized cFos+PV density (and proportion of PV+ interneurons expressing cFos) to WT levels (**Figure 8B-C**). Finally, as a proxy measure of E-I balance, we calculated the proportion of all cFos+ neurons that were also PV+ (cFos+PV/cFos). We found that despite differences in overall cFos expression, this ratio was similar in vehicle-treated WT and *Fmr1^-/y^* mice. While ML297 significantly reduced this ratio in both genotypes, the reduction was significantly more dramatic in WT mice (**Figure 8C**). Finally, we found a strong negative correlation between freezing behavior and cFos+PV neuronal density (and the proportion of all PV+ interneurons expressing cFos) in *Fmr1^-/y^* mice (**Figure 8D**). Overall, our data suggests overactivation of inhibitory and excitatory neuron activity in the DG network of *Fmr1^-/y^* mice is associated with disruption to CFM, and that this can be attenuated with ML297 administration during CFM consolidation.

To understand the impact of ML297 treatment on recall-associated neuronal activation across the rest of the dorsal hippocampus-amygdala circuit, we also measured cFos, PV, and PNN expression within CA1, CA3, and basolateral amygdala after CFM recall. Vehicle-treated *Fmr1^-/y^* mice showed dramatic reductions in cFos+ neuron density within CA1 compared with WT littermates (**Figure S7A-B**), no change in cFos+ neuron density in CA3 (**Figure S8A-B**), and modest reductions in cFos+ neuron density in amygdala (**Figure S9A-B**). ML297 treatment in *Fmr1^-/y^* mice increased cFos expression in CA1 (although expression also increased with ML297 in WT littermates), and had no effect for either genotype in CA3 or amygdala. CA1 cFos+ density was correlated with successful CFM recall in both WT and *Fmr1^-/y^* mice (**Figure S7D**), while density in amygdala and CA3 did not correlate with freezing levels. In CA1 and amygdala, PV+ interneuron and PNN density (and density of cFos+ and PNN+ PV+ interneurons) was significantly reduced in *Fmr1^-/y^* mice compared to WT littermates regardless of treatment (**Figure S7A-B; Figure S9A-B**). In CA3, PV+ interneuron and PNN densities were also modestly reduced in *Fmr1^-/y^* mice; here, ML297 treatment reversed the effects of genotype on PNN density. This suggests that alterations in the PV+ interneuron network of *Fmr1^-/y^* mice, which are present throughout the hippocampus-amygdala circuit, are largely unaffected by ML297 treatment.

Taken together, these findings suggest that *Fmr1^-/y^* mice show dramatic changes to hippocampus-amygdala network activity and interneuron function, which are evident during CFM recall. Post-CFC administration of ML297 alters network activity with the DG in *Fmr1^-/y^* mice, renormalizing activity levels in subsequent recall, but has few effects elsewhere in the circuit. The correlation of ML297 treatment’s effects on DG activation with memory performance outcomes suggest that DG activity is particularly important as a predictor of memory recall^77^, and ML297’s effects on this microcircuit are linked to rescue of CFM consolidation in *Fmr1^-/y^* mice.

## Discussion

To examine the relationship between disrupted sleep and cognitive phenotypes in FXS, we first fully characterized sleep in the *Fmr1^-/y^* FXS mouse model. While a few pieces of previous data suggested that *Fmr1^-/y^* mice could have disruptions to sleep comparable to FXS patients^20, 21, 80^, our long-term polysomnographic recordings provide the first conclusive evidence that mutant mice exhibit reduced NREM sleep, fragmented NREM architecture, and brain region-specific changes in NREM and REM sleep oscillations. These alterations are remarkably analogous to findings seen in polysomnography recordings of FXS and ASD patients^6–8^. For example, we observed increased delta power and decreased spindle density and power during NREM in *Fmr1^-/y^* mice. A similar phenotype has been reported in patients with ASD and/or FXS^13, 68, 81–85 60, 68^. Notably these genotype-driven EEG alterations are detected only in V1, not PFC. Critically, EEG recordings of ASD children and adults showed similar spatial (i.e., anteroposterior) differences^60, 61, 68^. These convergent findings suggest that the *Fmr1^-/y^* mouse model should be useful for studying sleep phenotypes related to FXS and ASD.

It is well-established that loss of *Fmr1* reduces excitatory drive onto PV+ interneurons and other GABAergic interneuron subtypes, while increasing the density of immature dendritic spines in neocortical and hippocampal pyramidal neurons ^14, 17, 19, 86–91^. These alterations are thought to lead to aberrant network excitability, which could underlie some of the sleep-associated EEG changes we observed in *Fmr1^-/y^* mice. Our data suggest that the TRN, which is populated by PV+ interneurons and essential for generation of NREM spindles ^58, 63, 92^, is likely affected by more general disruption of PV+ interneuron function in FXS. The reduction in PV+ interneuron density we observed in the TRN of *Fmr1^-/y^* mice is likely to lead to both reduced sleep spindle density, and reduced spindle coordination between cortical regions, in *Fmr1^-/y^* mice. Further study will be needed to understand the contribution of specific microcircuit-level alterations in *Fmr1^-/y^* mice to other changes in spectral power and coherence during NREM sleep.

GIRK1/2 function is thought to be selectively disrupted in the absence of FMRP^47, 48^. ML297, which activates GIRK1/2 current, has been found to promote NREM sleep without adverse effects to cognition (e.g. amnestic effects) observed with other hypnotic treatments^43, 45^. We found that acute administration of ML297 to *Fmr1^-/y^* mice was sufficient to renormalize NREM sleep amounts, sleep architecture, and EEG spectral power to parameters observed in WT littermates. In addition, we found spindle density correlations between V1 and PFC regions were normalized by ML297 treatment. Critically, available data suggest that GIRK1/2 channels are well expressed in (and modulate the physiology of) GABAergic TRN neurons^93^. Thus, ML297 activation of GIRK1/2 could be directly involved in improving synchronization of spindle oscillations.

Sleep is essential for cognitive functions, including memory consolidation^94, 95^. Given the relationship between sleep disruption and cognitive impairment, we tested whether restoration of sleep phenotypes could improve known memory consolidation deficits in *Fmr1^-/y^* mice. We found that ML297 improved consolidation of both CFM and OLM – two dorsal hippocampus-dependent, sleep-dependent forms of memory – which are normally disrupted in *Fmr1^-/y^* mice. Memory rescue of these memories by hypnotic treatment appeared to rely on sleep-dependent changes – e.g., rescue of CFM by ML297 was blocked by post-CFC SD. A parsimonious conclusion from these data is that ML297 rescues sleep-dependent memory consolidation by normalizing post-learning sleep behavior and sleep oscillations in *Fmr1^-/y^* mice.

As described above, loss of FMRP is thought to alter the structure and function of PV+ interneurons, and our present data support this conclusion. In every brain region examined in *Fmr1^-/y^* mice, we note a reduction in the density of PV+ interneurons, and in most brain regions we observed reduced PNN ensheathment of PV+ interneurons. In general, these changes were either unaffected, or only modestly changed, by ML297 administration. However, after CFM recall, we noticed changes to cFos expression in control (vehicle-treated) *Fmr1^-/y^* mice that occurred uniquely in DG – where overall expression of cFos was elevated, and cFos expression in PV+ interneurons was also elevated, relative to WT littermates. While additional studies will be needed to understand whether these changes are sufficient to cause disruption of CFM recall, we found that in ML297-treated *Fmr1^-/y^* mice – i.e. those with improved CFM consolidation – cFos expression in DG (both across DG as a whole, and in PV+ interneurons) was reduced. Moreover, the degree of reduction in cFos predicted better CFM performance. This suggests that improved sleep-dependent memory consolidation with ML297 treatment may be associated with suppression of aberrant, elevated DG activity in *Fmr1^-/y^* mice. While the role of altered E-I balance in this process is unclear, available data suggest that sleep itself can alter E-I balance in both cortical and hippocampal networks^96, 97^.

While our present findings suggest that sleep is an important potential therapeutic target for FXS, and potentially for other ASDs as well, additional work will be needed to clarify the translational potential of this work. For example, it is still unclear whether hippocampal network activity changes during recall are due to prior sleep architecture changes^77^ (or sleep-dependent oscillatory changes^98–100^) and those due to direct pharmacological effects of ML297 on hippocampal neurons themselves^101^. Future studies using hypnotic agents with distinct mechanisms of action will be needed to disentangle these potential mechanisms for hippocampal activity normalization in *Fmr1^-/y^* mice. In addition, further studies in juvenile mice will be needed to determine how relevant these findings are to treatment of pediatric FXS patients. Given the role of sleep has in the shaping of synaptic circuits early in life^30, 36, 102, 103^, treating sleep phenotypes in *Fmr1^-/y^* mice earlier in development has the potential to be even more beneficial for cognition. Nonetheless, these initial findings provide proof-of-concept for targeting sleep phenotypes as a therapeutic strategy for treating cognitive aspects of FXS.

## Acknowledgments

We thank the members of the Aton lab for useful feedback on these studies, Abbey Roelofs for coding support, and Gregg Sobocinski for microscopy support. Schematic images in Fig. 1A, 2A, 6A, 6C, 7A, and 7D were created with Biorender.com. This work was supported by a University of Michigan Graduate School Predoctoral Fellowship, Graduate Student Research Grant, and Merit Fellowship to JDM, a University of Michigan Kavli Neuroscience Innovators Magnificent Summer Fellowship to LGW and WPB, a Neuroscience Undergraduate Research Opportunity Summer Fellowship to REPT, a Brain and Behavior Research Foundation Young Investigator Award to SJA, and National Institutes of Health research grants R01 NS104776 and RF1 NS118440 to SJA.

## Author Contributions

JDM and SJA conceived and designed the study. JDM, LGW, WPB, KGP, VCG, REPT, DSP, MJD, SLMO, DT, SJ, and ZY performed the research. JDM, LGW, WPB, KGP, VCG, REPT, DSP, MJD, JJS, and BCC analyzed data. JDM and SJA wrote the manuscript. SJA supervised the study.

## STAR Methods

### Key Resources Table

See attached document.

### Resource Availability

#### Lead contact

Further information and requests for resources, code, and reagents should be directed to and will be fulfilled by the Lead Contact, Sara J. Aton.

#### Materials Availability

This study did not generate new unique reagents.

#### Code and data availability

Upon publication, code will be publicly available via **Github**. Any additional information required for data reported in this publication will be available from the lead contact, Sara J. Aton, upon request.

### Experimental Models and Subject Details

We generated male *Fmr1^-/y^* mice and wild-type (WT) littermates in-house by crossing WT male mice with *Fmr1* heterozygous (*Fmr1^+/-^*) female mice - both on a C57BL/6 background (The Jackson Laboratory, stock # 003025). Genotypes for mice were determined using standard PCR methods and primers described on the Jackson Laboratory website. For all experiments, 4-5 month old mice were used. Mice were housed under a 12:12 hour light/dark cycle (lights on at 9:00 AM), provided with compressed cotton and “Enviro-dri” paper nesting and bedding material (Shepherd Specialty Papers, TN), and provided *ad lib* access to water and food. Mice were housed with littermates until either EEG electrode implantation surgery or behavioral studies, at which point they were single housed in standard cages with extra nesting and bedding material for enrichment. All mouse husbandry, experimental, and surgical procedures were reviewed and approved by the University of Michigan Internal Animal Care and Use Committee.

### Method Details

#### EEG-EMG surgical procedures and neural data acquisition

Mice were anesthetized with 1-2% isoflurane anesthesia administered ketoprofen (0.005 mg/kg; intraperitoneal injection, i.p.) prior to surgery. Each mouse received two miniature, stainless steel screw EEG electrodes (P1 Technologies, Catalog #: E362/96/1.6/SPC) positioned over primary visual cortex (V1; 2.9 mm anterior/posterior [AP] and 2.7 mm medial/lateral [ML]) and prefrontal cortex (PFC; 1.8 mm ML). A reference screw was placed over cerebellum and a braided stainless steel wire EMG electrode was placed in the nuchal muscle. EMG electrodes were custom made from braided stainless-steel wire (Cooner Wire, Catalog No: AS636) soldered to miniature stainless steel male pins (P1 Technologies, Catalog No: 363A/PKG). The implant was secured using a pedestal (P1 Technologies, Catalog No: MS7P) bonded with super glue (Loctite). After 11 days of postoperative recovery, each mouse was moved to a new cage with an open top and habituated to tethering of flexible cables (P1 Technologies, Catalog No: 363-000) for 3 days before EEG/EMG data collection (Omniplex; Plexon Inc). Mice were recorded for four consecutive days (96 hours total). Day 1 was a 24-h baseline recording (Baseline A) starting at lights-on (ZT0). After 24 hours (Day 2), mice were injected with vehicle solution at lights-on and recorded. Day 3 was a second baseline recording (Baseline B). Day 4, mice were injected with ML297 (30 mg/kg) at lights-on and recorded for a final 24-hour period.

#### Pharmacological preparation and injection

ML297 was purchased from Tocris Bioscience (Catalog #. 5380). For all experiments, ML297 was initially dissolved in DMSO and diluted with 0.5% hydrooxypropyl cellulose aqueous solution. Mice were injected with a 30mg/kg solution of ML297, dosage based on previously published studies in rodent ^43, 45, 46^. Vehicle solutions consisted of 2% DMSO in 0.5% hydroxypropyl cellulose aqueous solution.

#### Contextual fear conditioning (CFC)

Mice underwent single-trial CFC as previously described^31, 32, 46, 70, 104, 105^. Mice were handled by the experimenter for 5 minutes daily for 3 days prior to training. Each mouse was placed in a novel cylindrical conditioning chamber made of clear Plexiglas with a black and white checkerboard pattern and metal grid floor (Med Associates). Before each individual mouse session, the arena was cleaned with a 5% Lysol solution. Mice were allowed to freely explore for 2 min and 28 s, after which they received a 0.75 mA, 2 s foot shock through the grid floor, followed by an additional 30 s in the CFC chamber. Immediately following CFC, mice were returned to their home cage. For studies on ML297, mice were given an i.p. injection of either ML297 (30 mg/kg; Tocris) or vehicle (2% DMSO in 0.5% hydrooxypropyl cellulose aqueous solution). Injections occurred within 2 min of removal from the CFC chamber. Contextual fear memory (CFM) tests were conducted 24 h later by returning mice to the CFC chamber for 5 min. Both CFC training and CFM testing began at lights-on (ZT0), and mice were video monitored continuously during both sessions.

#### Object location memory (OLM) task and open field (OF) test

Mice underwent a single trial OLM task as previously described^29, 72, 106^. Here, mice are placed in a rectangular arena made of grey PVC walls, transparent PVC bottom, and the following dimensions: length of 40 cm, width of 30 cm, and height of 30 cm. Spatial cues were placed on the opposite sides of the short walls and consisted of one black and white checkerboard pattern and one black and white striping pattern. Mice were handled by the experimenter for 5 min daily for 4 days prior to training. Prior to each session, the arena and objects were cleaned with a 10% ethanol solution. Before training, mice were habituated to the arena, which consisted of 5 minutes of free exploration and served as the OF test. The OF test is a rapid assessment of well-defined behaviors such as anxiety-related behaviors, body activity, and locomotion that require little to no prior training to the subject mouse ^107^. During training, a pair of identical objects were placed symmetrically across in the middle of the arena. Mice were placed in the arena facing the wall and were allowed to freely explore the arena and objects for 10 mins. Immediately following training, mice were returned to their home cage. For studies on ML297, mice were given an i.p. injection of either ML297 (30 mg/kg; Tocris) or vehicle (2% DMSO in 0.5% hydrooxypropyl cellulose aqueous solution). Injections occurred within 2 min of removal from the OLM arena. OLM tests were conducted 24 h later. During testing, one object was displaced (novel location) diagonally from the other object, which remained the same position (familiar location). The object displaced and novel object locations were counterbalanced to avoid confounding place and object preferences. Mice were again allowed to freely explore the arena and objects for 10 mins. Both OLM training and OLM testing began at lights-on (ZT0), and mice were video monitored continuously during both sessions and the habituation (OF test) period.

#### Sleep monitoring and sleep deprivation

Following CFC training, mice that were either allowed *ad lib* sleep (Sleep) or were sleep-deprived (SD) via gentle handing over the next 6 h (ZT0-6). This method of SD was chosen based on prior work showing that the glucocorticoid response evoked by gentle handling alone is not sufficient to disrupt consolidation of CFM (and in fact may enhance consolidation)^106, 108, 109^. These procedures included cage tapping or shaking, and/or nest disturbance and has been previously shown to ensure above 90% wake time based on EEG/EMG validation^46^. Following SD, all mice were allowed *ad lib* recovery sleep over the next 18 h prior to CFM testing. For *ad lib* sleep mice, sleep was quantified over the first 6 h post-CFC via visual monitoring. Every 5 min, individual mice were scored as awake or asleep, with sleep identification based on immobility, slow breathing, and presence of stereotyped (crouched) sleep postures, consistent with prior studies^110–112^ and has been validated with similar total sleep time in EEG/EMG implanted mice^46^.

#### Histology and immunohistochemistry

To quantify hippocampal activation patterns associated with recall, 90 min following the conclusion of CFM tests, mice were euthanized with an overdose of sodium pentobarbital and perfused with ice cold PBS, followed by ice cold 4% paraformaldehyde. Brains were dissected, post-fixed, and cryoprotected in a 30% sucrose solution. 50 µm coronal dorsal hippocampal and amygdala sections were cryosectioned. To quantify parvalbumin (PV) interneuron expression in the thalamic reticular nucleus (TRN), brains were dissected, post-fixed, and rinsed in PBS. 100 µm coronal sections containing TRN were collected via a vibratome.

For all tissue, free-floating sections were washed in PBS with 0.2% Triton X-100 (PBST) three times, each for 10 mins, and then incubated in Starting Block^TM^ blocking buffer (Thermo Scientific) for 1 hour. Hippocampal and amygdala sections were then incubated overnight in primary antibody at 4°C: rabbit-anti-cFos (1:1000; Abcam, ab190289), mouse-anti-PV (1:2000; Millipore, MAB11572), and wisteria floribunda agglutinin (lectin) tagged with fluorescein (WFA; 1:500; Vector Labs, FL-1351-2). TRN sections were only incubated with mouse-anti-PV (1:2000; Millipore, MAB11572). Sections were then washed in PBST, two times for 10 minutes and incubated with secondary antibody: Alexa Fluor 647 (1:200; Invitrogen, A11032) in 5% Starting Block blocking solution (cFos) and Alexa Fluor 594 (1:200; Invitrogen, A11034) in 5% Starting Block blocking solution (PV). Sections were washed in PBS, two times, mounted on microscope slides (Fisherbrand Superfrost Plus) and coverslipped with Prolong Gold antifade reagent with DAPI (Invitrogen, P36931).

### Quantification and Statistical Analysis

#### Sleep state and power spectra analysis

EEG/EMG signals (0.5-300 Hz) were amplified at 20 ×, digitized, further digitally amplified at 20-100 ×, and continuously recorded (with a 60-Hz notch filter applied to remove environmental noise) using Plexon Omniplex software and hardware (Plexon Inc., TX) as previously described Baseline and post-treatment recordings were scored manually in 10-second epochs as wake, NREM, or REM sleep using custom MATLAB software and previously published studies^31, 32, 46, 53, 104, 113^. Scorers were blind to genotype and recording day. EEG and EMG data were band-pass filtered at 0-60 Hz and 150-250 Hz, respectively, for viewing during scoring. NREM sleep was defined as synchronized, high amplitude, low frequency oscillations within the EEG and low EMG activity. REM sleep was defined as reduced, low frequency oscillations and a theta rhythm pattern (4-12 Hz) with lack of EMG activity. Wakefulness was defined as de-synchronized EEG activity with active EMG activity. For power spectral analysis, raw EEG data (0.5-300 Hz) was first filtered to 0.5-60 Hz using custom MATLAB scripts, Afterwards, using fast-Fourier transform (FFT) of EEG data was carried out Neuroexplorer 5 software (Nex Technologies).

#### Sleep spindle identification and spectral coherence between brain regions

An automated spindle detection algorithm (custom MATLAB software) was used to identify sleep spindles in band-pass filtered EEG data (7-15 Hz), as intervals containing ≥ 6 successive deviations (i.e., peaks or troughs) of signal that surpassed mean signal amplitude by 1.5 standard deviations, lasting between 0.25-1.75 seconds ^46, 53^. The coherence of raw EEG data at given frequencies was analyzed using Neuroexplorer 5 software (Nex Technologies) using previously published methods^114, 115^. Coherence was calculated using a Welch taper with a 50% window overlap. Coherence generated values are expressed from 0 to 1.0, with 0 indicating no relationship between two signals (two brain regions) at a given frequency and 1.0 indicating a perfect, linear relationship. Band coherence for frequency bands of interest was assessed by averaging the coherence values with each designated frequency band.

#### Behavior quantification

All behavioral measurements were assessed using Ethovision XT 16 (Noldus) software with files coded for blind scoring. For CFC behavior, freezing was measured using in a semi-automated manner. Freezing was first scored based on transient periods of immobility, as described previously^46, 116^ and was verified offline based on the assessment of characteristic freezing-associated posture^32, 113^. CFM-associated freezing behavior was quantified by subtracting each mouse’s % freezing time during pre-shock baseline from % freezing time across the entire CFM test, as described previously^32, 46, 70, 104^. For OF test (habituation of OLM task), the total ambulatory distance was measured to account for differences in locomotor ability. Thigmotaxis (or wall-hugging behavior), an anxiety-related behavior, was measured by comparing percentage of time spend in the outer (wall) zone vs. the inner (center) zone^107^. Both locomotor ability and thigmotaxis were measured to ensure further analyses in OLM were not skewed due to inactivity instead of genotype or treatment effects. For OLM behavior, measurements were based on time mice spent exploring the familiar and displaced object. Engagement with object included directing nose to the object at no more than 1 cm and/or touching the object. A discrimination index was used a relative measure of discrimination corrected for total exploration time^29^.

#### Microscopy and image analysis

Images were obtained using a Leica SP8 confocal microscope with a 10× objective, to obtain z-stack images (10 µm steps) for maximum projection of fluorescence signals. Identical image acquisition settings (e.g. exposure times, frame average, pixel size) were used for all sections. For analysis of hippocampal and amygdala activation patterns, three images per section of dorsal hippocampus and amygdala were taken per mouse and areas for regions of interest (ROI) were determined using previously established methods^70, 110^. cFos+, PV+, and PNN density was quantified in subregions across these ROIs (DG, CA1, and CA3). For analysis of PV+ interneuron expression within the TRN, cell density was measured using the number of PV+ cells counted within the area made via the ROI surrounding the TRN. Counts were made by two scorers blinded to genotype and treatment using Fiji image analysis software, with the final number computed using the average of counts made by the two scorers.

#### Statistical analysis

Data analyses were carried out in a blind manner; in some cases (e.g., EEG recordings and cell counts), data was consensus scored by two individuals to reduce variability. Exclusion criteria for EEG recordings were based on lack of signal in one cortical region or faulty reference electrode that prevented confirmation of recording signals. For CFC behavior, exclusion criteria consisted of freezing levels above 15% before shock administration during CFC training sessions. For OLM behavior, exclusion criteria included mice that spend less than 10 seconds interacting with objects during 10-minute training sessions or exhibited impaired locomotor ability and/or thigmotaxis. Statistical analyses were carried out using GraphPad Prism software (Version 9.5). For each specific data set, the statistical tests and *p*-values are listed within the appropriate corresponding figure legend. Data sets were tested for normality to determine use of parametric or non-parametric tests.

---

**Figure S1:**
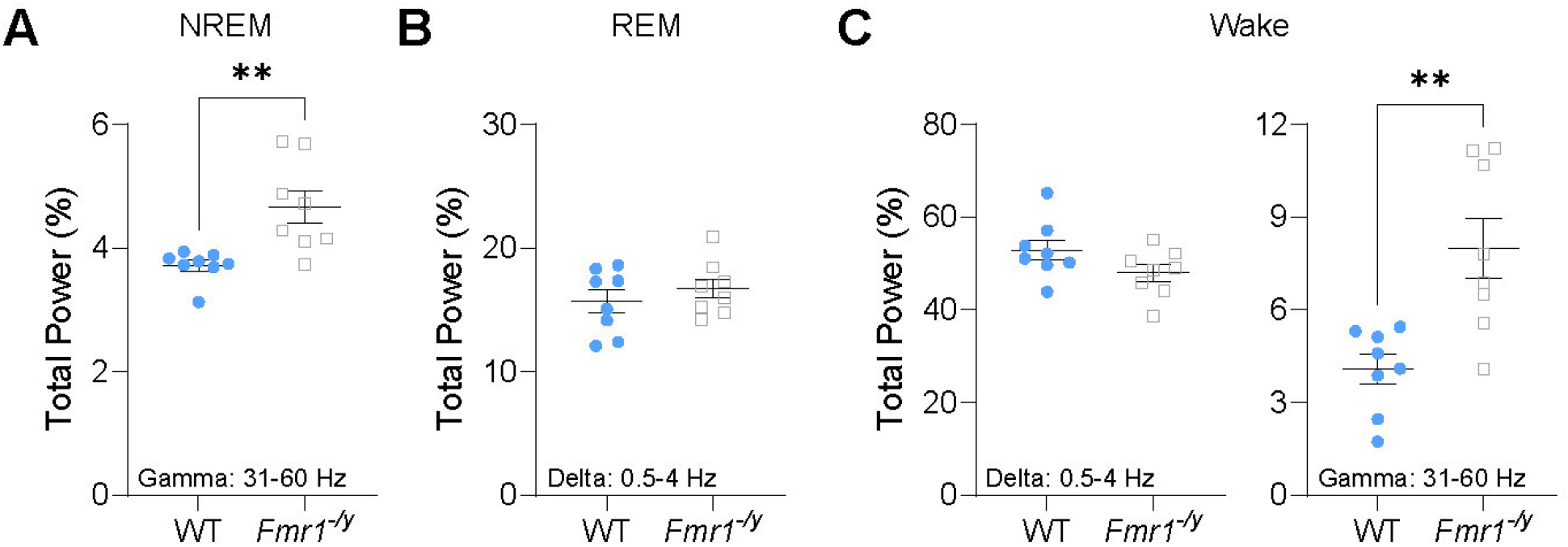
*Fmr1^-/y^*mice NREM gamma, REM delta, and waking spectral power during the rest phase. **(A)** Total gamma-band (31-100 Hz) power was increased in *Fmr1^-/y^* mice during NREM sleep across the 12-h light phase. **(B)** Total delta-band (0.5-4 Hz) power during REM sleep across the 12-h light phase. **(C)** Total delta-band (***left***) and gamma-band (***right***) power were reduced and increased, respectively, in *Fmr1^-/y^* mice during wake across the 12-h light phase. Sample size: *n* = 8 mice/genotype. ** indicates *p* < 0.01, two-tailed, unpaired t-test (**E-H**). Data points and error bars indicate mean ± SEM. (Related to Fig. 1)

**Figure S2:**
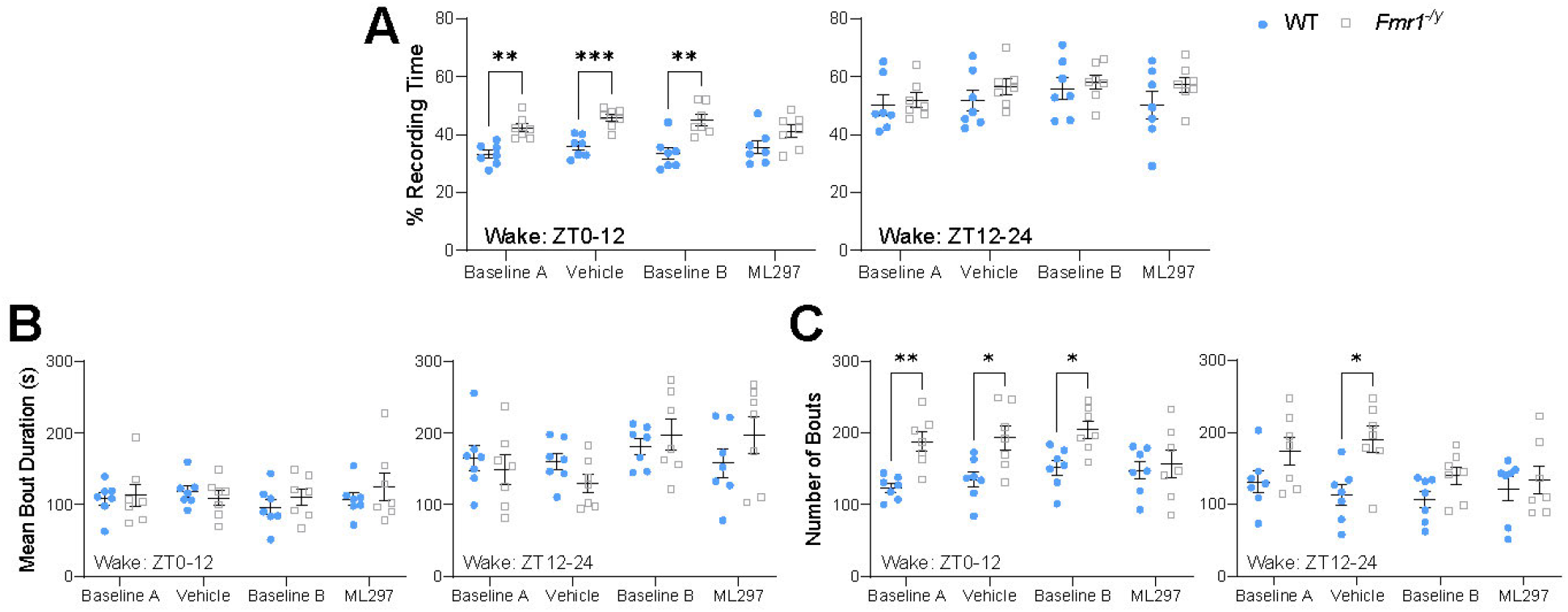
ML297 normalizes wake architecture differences in *Fmr1^-/y^* mice. **(A)** Waking percent recording time for the 0-12 h light phase (***left***) and 12-24 h dark phase (***right***) across all four days for WT and *Fmr1^-/y^* mice. Two-way RM ANOVA for ZT0-12: *p*(condition) = 0.24, *p*(genotype) = 0.0002, *p*(condition x genotype interaction) = 0.27 and ZT12-24: *p*(condition) = 0.035, *p*(genotype) = 0.35, *p*(condition x genotype interaction) = 0.41. **(B)** Waking bout duration for the 0-12 h light phase (***left***) and 12-24 h dark phase (***right***) across all four days for WT and *Fmr1^-/y^* mice. Two-way RM ANOVA for ZT0-12: *p*(condition) = 0.35, *p*(genotype) = 0.64, *p*(condition x genotype interaction) = 0.31 and ZT12-24: *p*(condition) = 0.0037, *p*(genotype) = 0.93, *p*(condition x genotype interaction) = 0.025. **(C)** Number of wakefulness bouts for the 0-12 h light phase (***left***) and 12-24 h dark phase (***right***) across all four days for WT and *Fmr1^-/y^* mice. Two-way RM ANOVA for ZT0-12: *p*(condition) = 0.039, *p*(genotype) = 0.0097, *p*(condition x genotype interaction) = 0.011 and ZT12-24: *p*(condition) = 0.052, *p*(genotype) = 0.023, *p*(condition x genotype interaction) = 0.027. Sample size: *n* = 7 mice/genotype. *, ** and *** indicate *p* < 0.05, *p* < 0.01, and *p* < 0.001, Sidak’s *post hoc* test. Data points and error bars indicate mean ± SEM. (Related to Fig. 2)

**Figure S3:**
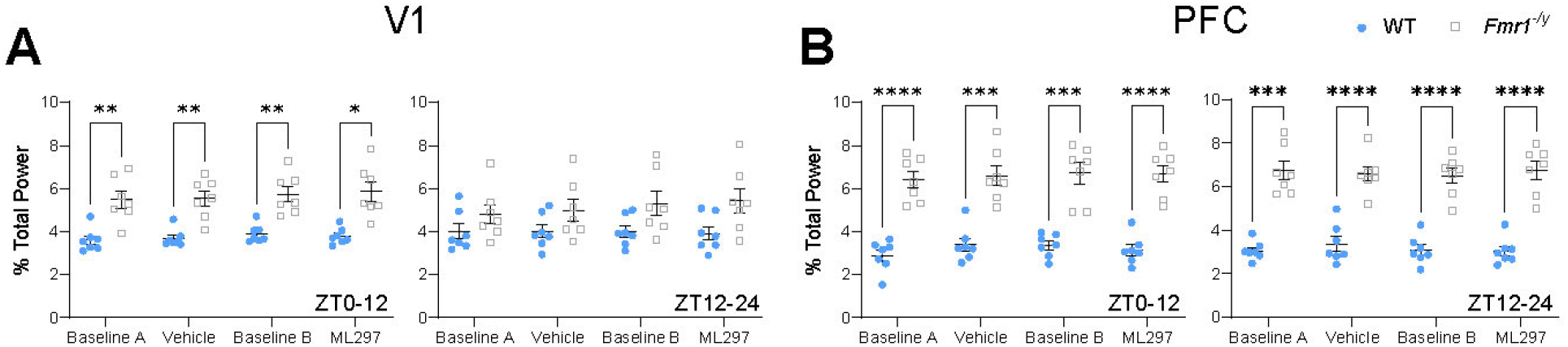
NREM gamma power is increased in V1 and PFC in WT and *Fmr1^-/y^* mice. (A) Total NREM gamma-band power (31-60 Hz) during the light phase (ZT0-12) (***left***) and dark phase (ZT12-24) (***right***) in V1. Two-way RM ANOVA for ZT0-12: *p*(condition) = 0.12, *p*(genotype) = 0.0004, *p*(condition x genotype interaction) = 0.79 and ZT12-24: *p*(condition) = 0.19, *p*(genotype) = 0.073, *p*(condition x genotype interaction) = 0.054. (B) Total NREM gamma-band power (31-60 Hz) during the light phase (ZT0-12) (***left***) and dark phase (ZT12-24) (***right***) in PFC. Two-way RM ANOVA for ZT0-12: *p*(condition) = 0.23, *p*(genotype) < 0.0001, *p*(condition x genotype interaction) = 0.79 and ZT12-24: *p*(condition) = 0.80, *p*(genotype) < 0.0001, *p*(condition x genotype interaction) = 0.58. Sample size: *n* = 7 mice/genotype. *, **, *** and **** indicate *p* < 0.05, *p* < 0.01, *p* < 0.001, and *p* < 0.0001, Sidak’s *post hoc* test. Data points and error bars indicate mean ± SEM. (Related to Fig. 3)

**Figure S4:**
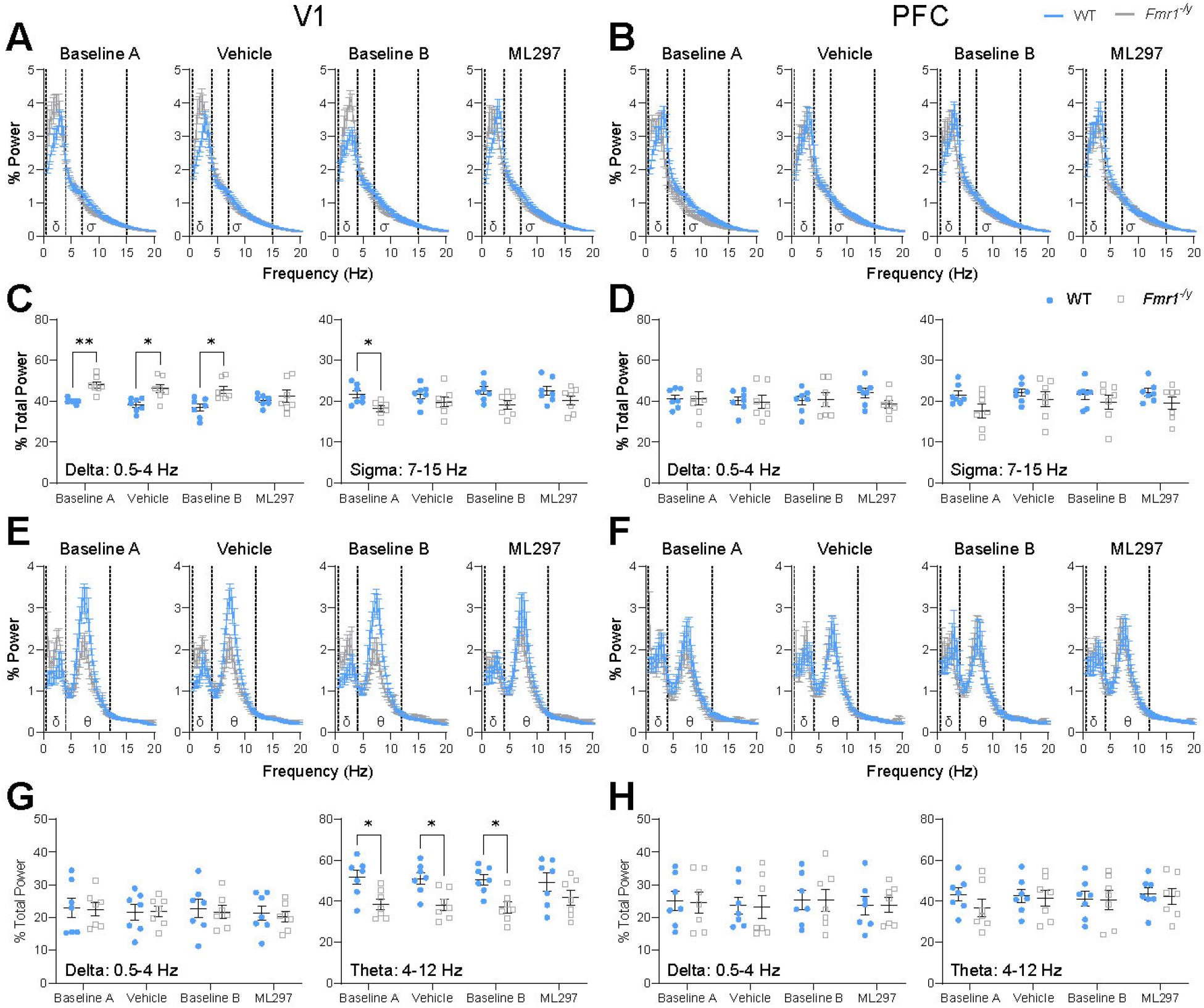
NREM and REM spectral power in V1, but not PFC, is altered in *Fmr1^-/y^* mice and normalized with ML297 during the dark phase. (A) NREM EEG power spectra in V1 for baselines and treatment conditions during ZT12-24 (dark phase) with delta (δ; 0.5-4 Hz) and sigma (σ; 7-15 Hz) frequency bands highlighted via dashed lines. Two-way RM ANOVA, *p*(frequency) < 0.0001, *p*(frequency x genotype interaction) < 0.0001, for baseline A-ML297 conditions and *p*(genotype) = 0.11 (baseline A), = 0.002 (vehicle), = 0.072 (baseline B), and = 0.26 (ML297). (B) NREM EEG power spectra in PFC for baselines and treatment conditions during ZT12-24 (dark phase) with delta (δ; 0.5-4 Hz) and sigma (σ; 7-15 Hz) frequency bands highlighted via dashed lines. Two-way RM ANOVA, *p*(frequency) < 0.0001, *p*(frequency x genotype interaction) < 0.0001, for baseline A-ML297 conditions and *p*(genotype) = 0.091 (baseline A), = 0.33 (vehicle), = 0.42 (baseline B), and = 0.001 (ML297). (C) Total NREM delta-band power (***left***) and total sigma-band power for spindles (***right***) in V1 during ZT12-24. Two-way RM ANOVA for total delta power: *p*(condition) = 0.048, *p*(genotype) = 0.0082, *p*(condition x genotype interaction) = 0.0098 and total sigma power: *p*(condition) = 0.35, *p*(genotype) = 0.019, *p*(condition x genotype interaction) = 0.64. (D) Total NREM delta-band power (***left***) and total sigma-band power for spindles (***right***) in PFC during ZT12-24. Two-way RM ANOVA for total delta power: *p*(condition) = 0.69, *p*(genotype) = 0.65, *p*(condition x genotype interaction) = 0.29 and total sigma power: *p*(condition) = 0.17, *p*(genotype) = 0.17, *p*(condition x genotype interaction) = 0.43. (E) REM EEG power spectra in V1 for baselines and treatment conditions during ZT12-24 (dark phase) with delta (δ; 0.5-4 Hz) and theta (θ; 4-12 Hz) frequency bands highlighted via dashed lines. Two-way RM ANOVA, *p*(frequency) < 0.0001, *p*(frequency x genotype interaction) < 0.0001, for baseline A, vehicle, and baseline B conditions; ML297 condition, *p*(frequency) < 0.0001, *p*(frequency x genotype interaction) = 0.13 and *p*(genotype) = 0.037 (baseline A), = 0.053 (vehicle), = 0.077 (baseline B), and = 0.26 (ML297). (F) REM EEG power spectra in PFC for baselines and treatment conditions during ZT12-24 (dark phase) with delta (δ; 0.5-4 Hz) and theta (θ; 4-12 Hz) frequency bands highlighted via dashed lines. Two-way RM ANOVA, *p*(frequency) < 0.0001, *p*(frequency x genotype interaction) > 0.99, for baseline A-ML297 conditions and *p*(genotype) = 0.14 (baseline A), = 0.84 (vehicle), = 0.85 (baseline B), and = 0.76 (ML297). (G) Total REM delta-band power (***left***) and total theta-band power (***right***) in V1 during ZT12-24. Two-way RM ANOVA for total delta power: *p*(condition) = 0.40, *p*(genotype) = 0.84, *p*(condition x genotype interaction) = 0.93 and total theta power: *p*(condition) = 0.56, *p*(genotype) = 0.011, *p*(condition x genotype interaction) = 0.24. (H) Total REM delta-band power (***left***) and total theta-band power (***right***) in PFC during ZT12-24. Two-way RM ANOVA for total delta power: *p*(condition) = 0.45, *p*(genotype) = 0.97, *p*(condition x genotype interaction) = 0.99 and total theta power: *p*(condition) = 0.24, *p*(genotype) = 0.64, *p*(condition x genotype interaction) = 0.13. Sample size: *n* = 7 mice/genotype. * and ** indicate *p* < 0.05 and *p* < 0.01, Sidak’s *post hoc* test. Data points and error bars indicate mean ± SEM. (Related to Fig. 3)

**Figure S5:**
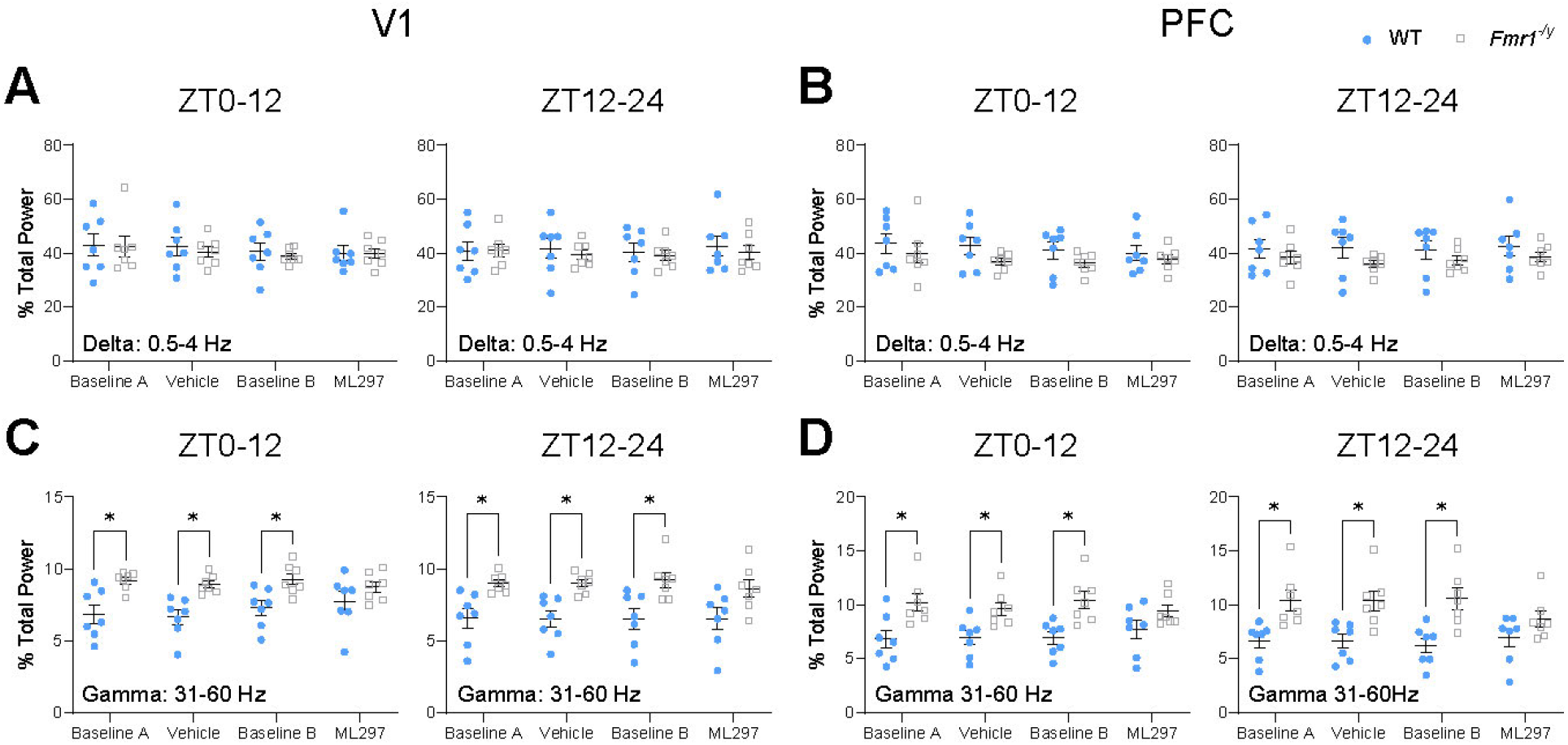
Gamma power during wake is altered in *Fmr1^-/y^* mice, and normalized by ML297. (A) Total delta-band power (0.5-4 Hz) during wakefulness for ZT0-12 (***left***) and ZT12-24 (***right***) in V1. Two-way RM ANOVA for ZT0-12: *p*(condition) = 0.26, *p*(genotype) = 0.78, *p*(condition x genotype interaction) = 0.93 and ZT12-24: *p*(condition) = 0.70, *p*(genotype) = 0.71, *p*(condition x genotype interaction) = 0.83. (B) Total delta-band power (0.5-4 Hz) during wakefulness for ZT0-12 (***left***) and ZT12-24 (***right***) in PFC. Two-way RM ANOVA for ZT0-12: *p*(condition) = 0.26, *p*(genotype) = 0.24, *p*(condition x genotype interaction) = 0.72 and ZT12-24: *p*(condition) = 0.66, *p*(genotype) = 0.25, *p*(condition x genotype interaction) = 0.82. (C) Total gamma-band power (31-60 Hz) during wakefulness for ZT0-12 (***left***) and ZT12-24 (***right***) in V1. Two-way RM ANOVA for ZT0-12: *p*(condition) = 0.37, *p*(genotype) = 0.0043, *p*(condition x genotype interaction) = 0.20 and ZT12-24: *p*(condition) = 0.90, *p*(genotype) = 0.0007, *p*(condition x genotype interaction) = 0.95. (D) Total gamma-band power (31-60 Hz) during wakefulness for ZT0-12 (***left***) and ZT12-24 (***right***) in V1. Two-way RM ANOVA for ZT0-12: *p*(condition) = 0.51, *p*(genotype) = 0.012, *p*(condition x genotype interaction) = 0.017 and ZT12-24: *p*(condition) = 0.24, *p*(genotype) = 0.0065, *p*(condition x genotype interaction) = 0.013. Sample size: *n* = 7 mice/genotype. * indicates *p* < 0.05, Sidak’s *post hoc* test. Data points and error bars indicate mean ± SEM. (Related to Fig. 3)

**Figure S6:**
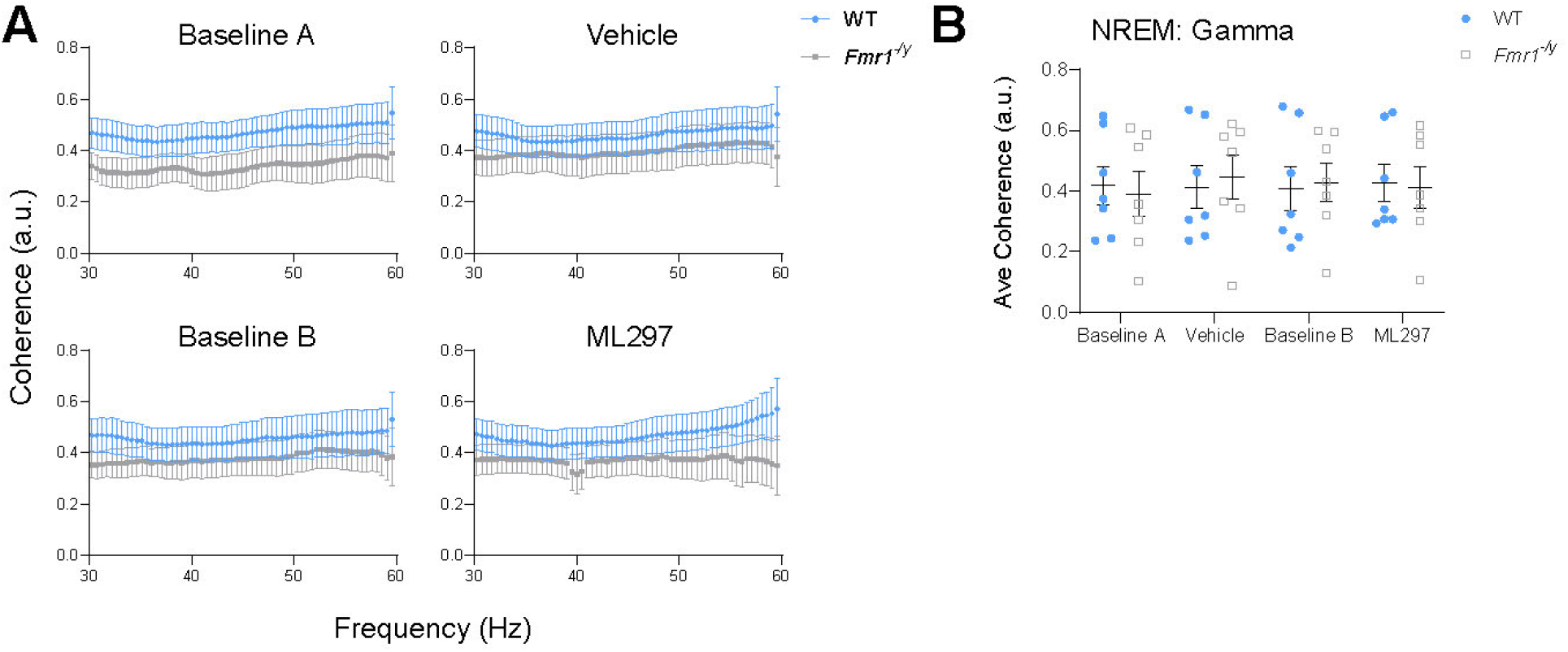
NREM gamma coherence between WT and *Fmr1^-/y^* mice. **(A)** NREM-associated V1-PFC spectral coherence for 31-60 Hz across baseline and treatment sessions. Two-way ANOVA, *p*(frequency) > 0.99, *p*(genotype) < 0.0001, *p*(frequency x genotype interaction) > 0.99, for baseline A-ML297 conditions. **(B)** Average NREM V1-PFC gamma-band (31-60 Hz) coherence. Two-way RM ANOVA: *p*(condition) = 0.49, *p*(genotype) = 0.98, *p*(condition x genotype interaction) = 0.34. Sample size: *n* = 7 mice/genotype. Data points and error bars indicate mean ± SEM. (Related to Fig. 5)

**Figure S7:**
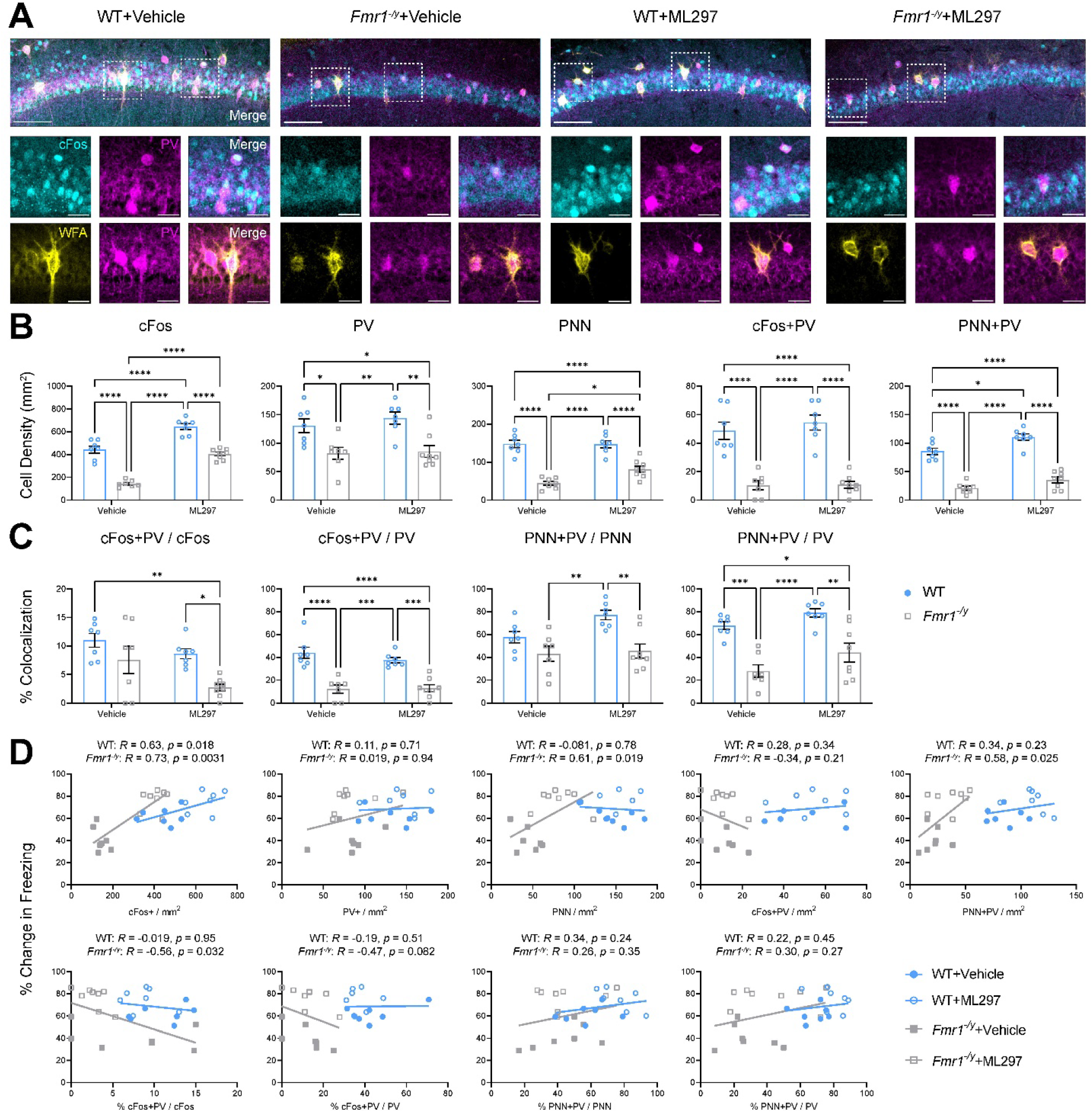
*Fmr1^-/y^*mice show reduced CA1 network activity during memory recall, which is not rescued by ML297 administration. **(A)** Representative fear memory recall-associated cFos (cyan), PV (magenta), and wisteria floribunda agglutinin (WFA; yellow) for PNN expression in CA1 for the four treatment groups. Scale bar = 250 µm (***top***, whole CA1); scale bar = 30 µm (***bottom***, inset of select neurons). **(B)** Quantification of CA1 cFos+, PV+, PNN, and co-localization neuronal density. Two-way ANOVA, for cFos+ neurons: *p*(treatment) < 0.0001, *p*(genotype) < 0.0001, *p*(treatment x genotype interaction) = 0.21. For PV+ neurons: *p*(treatment) = 0.44, *p*(genotype) < 0.0001, *p*(treatment x genotype interaction) = 0.66. For PNN: *p*(treatment) = 0.039, *p*(genotype) < 0.0001, *p*(treatment x genotype interaction) = 0.033. For cFos+PV co-localized neurons: *p*(treatment) = 0.48, *p*(genotype) < 0.0001, *p*(treatment x genotype interaction) = 0.54. For PNN+PV co-localized neurons: *p*(treatment) = 0.0008, *p*(genotype) < 0.0001, *p*(treatment x genotype interaction) = 0.30. **(C)** Quantification of CA1 percent co-localization of cFos+, PV+, and PNN over total neuronal populations. Two-way ANOVA, for % cFos+PV cells over total cFos: *p*(treatment) = 0.018, *p*(genotype) = 0.0029, *p*(treatment x genotype interaction) = 0.38. For % cFos+PV cells over total PV: *p*(treatment) = 0.42, *p*(genotype) < 0.0001, *p*(treatment x genotype interaction) = 0.32. For % PNN+PV cells over total PNN: *p*(treatment) = 0.065, *p*(genotype) = 0.0004, *p*(treatment x genotype interaction) = 0.15. For % PNN+PV cells over total PV: *p*(treatment) = 0.026, *p*(genotype) < 0.0001, *p*(treatment x genotype interaction) = 0.67. **(D)** Pearson correlation of cFos+, PV+, and PNN neuron densities and associated co-localizations in CA1 vs. associated change in freezing behavior during CFM recall. *R* and *p* values for Pearson correlation of each data set are found alongside corresponding graph. Sample sizes: *n* = 7 (WT+vehicle), *n* = 7 (*Fmr1^-/y^*+vehicle), *n* = 7 (WT+ML297), *n* = 8 (*Fmr1^-/y^*+ML297). *, **, ***, and **** indicate *p* < 0.05, *p* < 0.01, *p* < 0.001, *p* < 0.0001; Tukey’s *post hoc* test. Data shown as mean ± SEM. (Related to Fig. 8)

**Figure S8:**
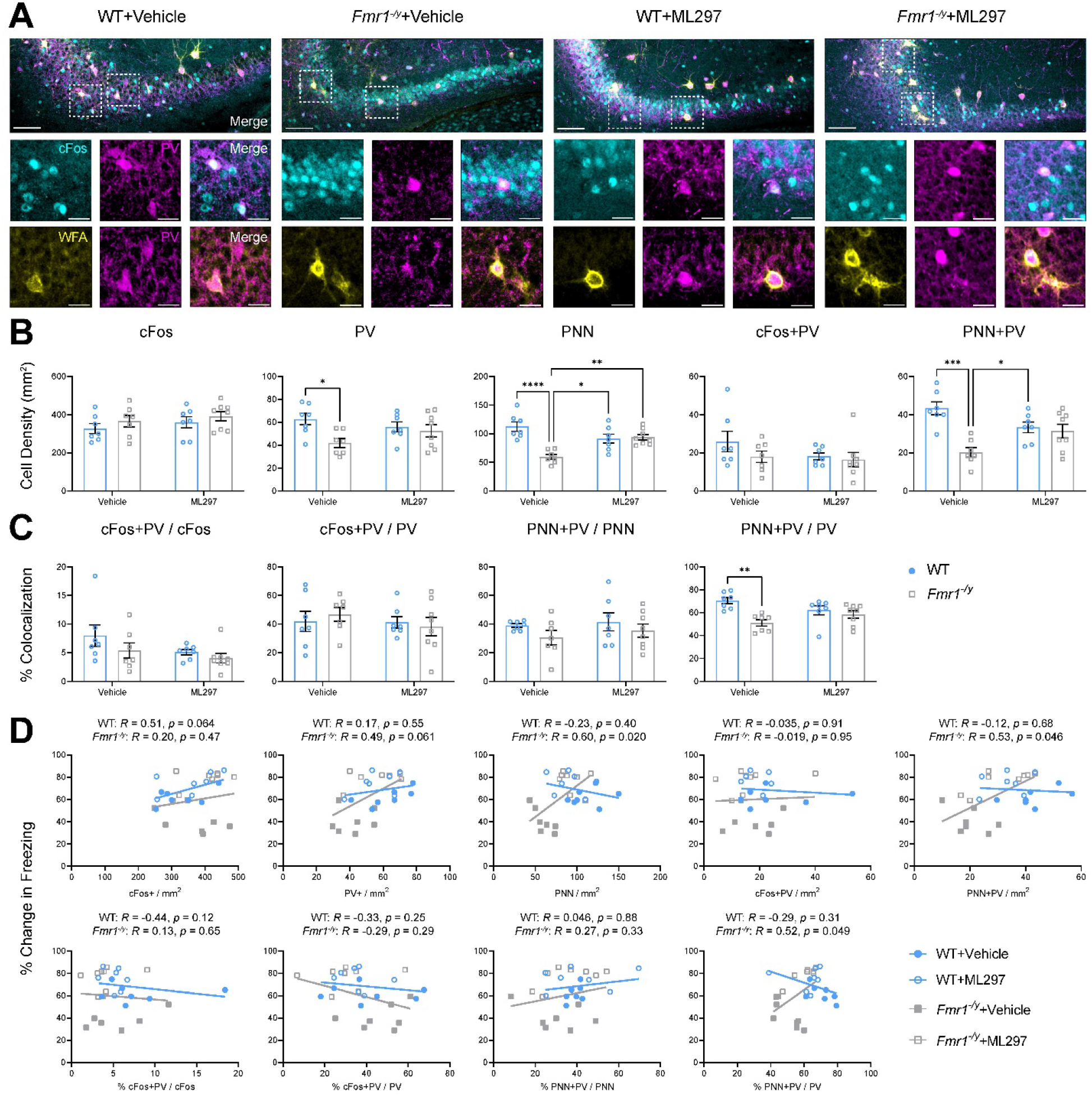
*Fmr1^-/y^* mice show alterations to CA3 PV+ interneurons and PNNs that are normalized by ML297. **(A)** Representative fear memory recall-associated cFos (cyan), PV (magenta), and wisteria floribunda agglutinin (WFA; yellow) for PNN expression in CA3 for the four treatment groups. Scale bar = 250 µm (***top***, whole CA3); scale bar = 30 µm (***bottom***, inset of select neurons). **(B)** Quantification of CA3 cFos+, PV+, PNN, and co-localization neuronal density. Two-way ANOVA, for cFos+ neurons: *p*(treatment) = 0.30, *p*(genotype) = 0.21, *p*(treatment x genotype interaction) = 0.91. For PV+ neurons: *p*(treatment) = 0.68, *p*(genotype) = 0.018, *p*(treatment x genotype interaction) = 0.083. For PNN: *p*(treatment) = 0.32, *p*(genotype) = 0.0006, *p*(treatment x genotype interaction) = 0.0002. For cFos+PV co-localized neurons: *p*(treatment) = 0.23, *p*(genotype) = 0.21, *p*(treatment x genotype interaction) = 0.41. For PNN+PV co-localized neurons: *p*(treatment) = 0.86, *p*(genotype) = 0.0005, *p*(treatment x genotype interaction) = 0.0026. **(C)** Quantification of CA3 percent co-localization of cFos+, PV+, and PNN over total neuronal populations. Two-way ANOVA, for % cFos+PV cells over total cFos: *p*(treatment) = 0.093, *p*(genotype) = 0.14, *p*(treatment x genotype interaction) = 0.52. For % cFos+PV cells over total PV: *p*(treatment) = 0.044, *p*(genotype) = 0.95, *p*(treatment x genotype interaction) = 0.33. For % PNN+PV cells over total PNN: *p*(treatment) = 0.43, *p*(genotype) = 0.13, *p*(treatment x genotype interaction) = 0.79. For % PNN+PV cells over total PV: *p*(treatment) = 0.87, *p*(genotype) = 0.0014, *p*(treatment x genotype interaction) = 0.022. **(D)** Pearson correlation of cFos+, PV+, and PNN neuron densities and associated co-localizations in CA3 vs. associated change in freezing behavior during CFM recall. *R* and *p* values for Pearson correlation of each data set are found alongside corresponding graph. Sample sizes: *n* = 7 (WT+vehicle), *n* = 7 (*Fmr1^-/y^*+vehicle), *n* = 7 (WT+ML297), *n* = 8 (*Fmr1^-/y^*+ML297). *, **, ***, and **** indicate *p* < 0.05, *p* < 0.01, *p* < 0.001, *p* < 0.0001; Tukey’s *post hoc* test. Data shown as mean ± SEM. (Related to Fig. 8)

**Figure S9:**
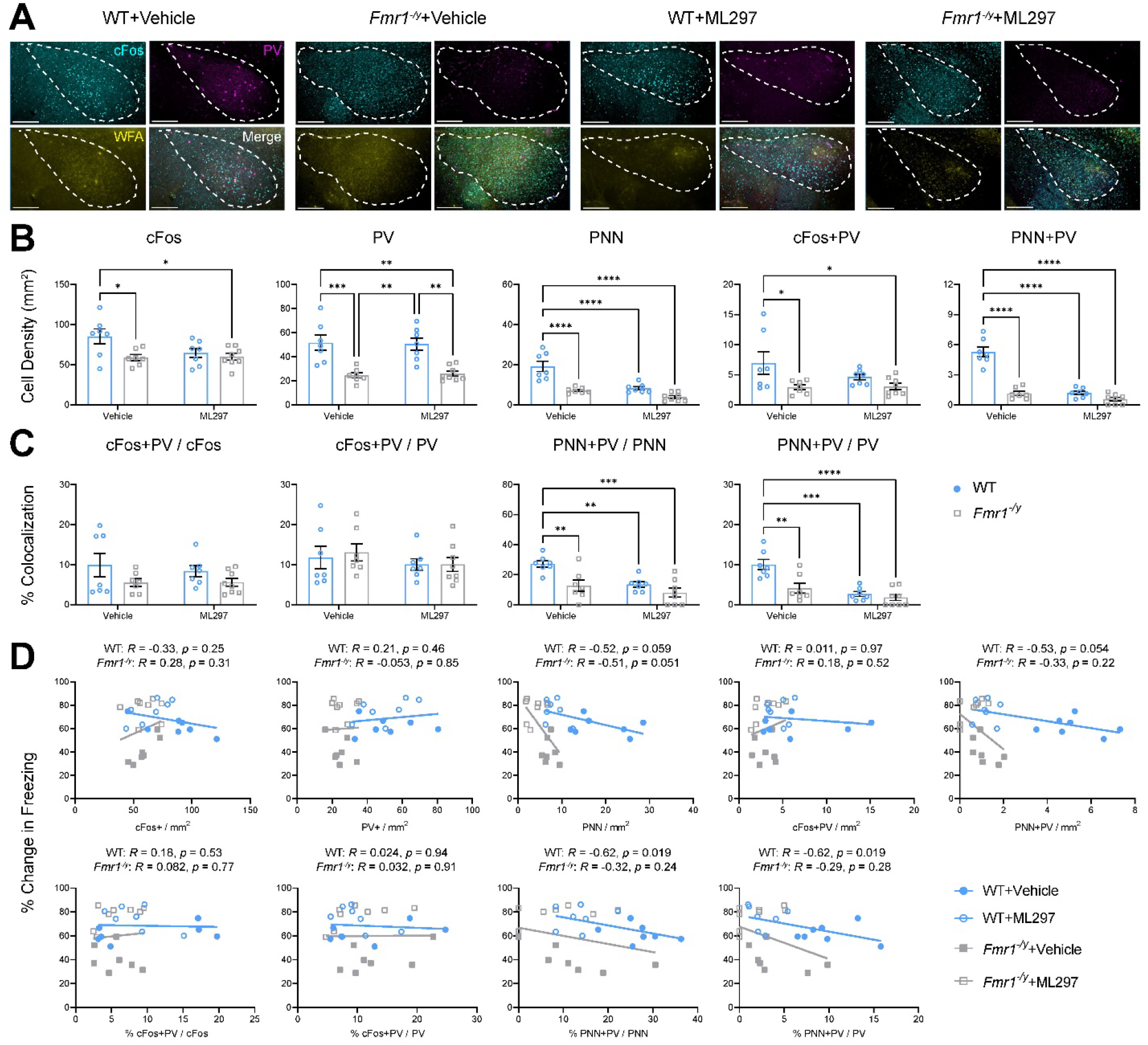
*Fmr1^-/y^* mice show altered amygdala activity and PV+ interneuron density, which are not rescued by ML297 administration. **(A)** Representative fear memory recall-associated cFos (cyan), PV (magenta), and wisteria floribunda agglutinin (WFA; yellow) for PNN expression in the amygdala for the four treatment groups. Scale bar = 250 µm (***top***, amygdala); scale bar = 30 µm (***bottom***, inset of select neurons). **(B)** Quantification of cFos+, PV+, PNN, and co-localization neuronal density. Two-way ANOVA, for cFos+ neurons: *p*(treatment) = 0.12, *p*(genotype) = 0.018, *p*(treatment x genotype interaction) = 0.085. For PV+ neurons: *p*(treatment) = 0.98, *p*(genotype) < 0.0001, *p*(treatment x genotype interaction) = 0.77. For PNN: *p*(treatment) < 0.0001, *p*(genotype) < 0.0001, *p*(treatment x genotype interaction) = 0.01. For cFos+PV co-localized neurons: *p*(treatment) = 0.26, *p*(genotype) = 0.0088, *p*(treatment x genotype interaction) = 0.23. For PNN+PV co-localized neurons: *p*(treatment) < 0.0001, *p*(genotype) < 0.0001, *p*(treatment x genotype interaction) < 0.0001. **(C)** Quantification of percent co-localization of cFos+, PV+, and PNN over total neuronal populations. Two-way ANOVA, for % cFos+PV cells over total cFos: *p*(treatment) = 0.67, *p*(genotype) = 0.048, *p*(treatment x genotype interaction) = 0.66. For % cFos+PV cells over total PV: *p*(treatment) = 0.27, *p*(genotype) = 0.76, *p*(treatment x genotype interaction) = 0.77. For % PNN+PV cells over total PNN: *p*(treatment) = 0.0027, *p*(genotype) = 0.0013, *p*(treatment x genotype interaction) = 0.11. For % PNN+PV cells over total PV: *p*(treatment) < 0.0001, *p*(genotype) = 0.0026, *p*(treatment x genotype interaction) = 0.018. **(D)** Pearson correlation of cFos+, PV+, and PNN neuron densities and associated co-localizations in amygdala vs. associated change in freezing behavior during CFM recall. *R* and *p* values for Pearson correlation of each data set are found alongside corresponding graph. Sample sizes: *n* = 7 (WT+vehicle), *n* = 7 (*Fmr1^-/y^*+vehicle), *n* = 7 (WT+ML297), *n* = 8 (*Fmr1^-/y^*+ML297). *, **, ***, and **** indicate *p* < 0.05, *p* < 0.01, *p* < 0.001, *p* < 0.0001; Tukey’s *post hoc* test. Data shown as mean ± SEM. (Related to Fig. 8)

## References

1 Rogers, S. J., Wehner, D. E. & Hagerman, R. The behavioral phenotype in fragile X: symptoms of autism in very young children with fragile X syndrome, idiopathic autism, and other developmental disorders. J Dev Behav Pediatr 22, 409–417 (2001). https://doi.org:10.1097/00004703-200112000-00008

2 Dahlhaus, R. in Frontiers in Molecular Neuroscience Vol. 11 41–41 (Frontiers Media S.A., 2018).

3 Baumgardner, T. L., Reiss, A. L., Freund, L. S. & Abrams, M. T. Specification of the neurobehavioral phenotype in males with fragile X syndrome. Pediatrics 95, 744–752 (1995).

4 Garber, K. B., Visootsak, J. & Warren, S. T. Fragile X syndrome. Eur J Hum Genet 16, 666–672 (2008). https://doi.org:10.1038/ejhg.2008.61

5 Hartley, S. L. et al. Exploring the adult life of men and women with fragile X syndrome: results from a national survey. Am J Intellect Dev Disabil 116, 16–35 (2011). https://doi.org:10.1352/1944-7558-116.1.16

6 Kronk, R. et al. Prevalence, nature, and correlates of sleep problems among children with fragile X syndrome based on a large scale parent survey. Sleep 33, 679–687 (2010). https://doi.org:10.1093/sleep/33.5.679

7 Kronk, R., Dahl, R. & Noll, R. Caregiver reports of sleep problems on a convenience sample of children with fragile X syndrome. Am J Intellect Dev Disabil 114, 383–392 (2009). https://doi.org:10.1352/1944-7588-114.6.383

8 Kidd, S. A. et al. Fragile X syndrome: a review of associated medical problems. Pediatrics 134, 995–1005 (2014). https://doi.org:10.1542/peds.2013-4301

9 Weiskop, S., Richdale, A. & Matthews, J. Behavioural treatment to reduce sleep problems in children with autism or fragile X syndrome. Dev Med Child Neurol 47, 94–104 (2005). https://doi.org:10.1017/s0012162205000186

10 Colten, H. R. & Altevogt, B. M. Sleep disorders and sleep deprivation: An unmet public health problem. (National Academies Press, 2006).

11 Liu, X., Hubbard, J. A., Fabes, R. A. & Adam, J. B. Sleep disturbances and correlates of children with autism spectrum disorders. Child Psychiatry Hum Dev 37, 179–191 (2006). https://doi.org:10.1007/s10578-006-0028-3

12 Sateia, M. J. International classification of sleep disorders-third edition: highlights and modifications. Chest 146, 1387–1394 (2014). https://doi.org:10.1378/chest.14-0970

13 Blackmer, A. B. & Feinstein, J. A. Management of Sleep Disorders in Children With Neurodevelopmental Disorders: A Review. Pharmacotherapy 36, 84–98 (2016). https://doi.org:10.1002/phar.1686

14 Hays, S. A., Huber, K. M. & Gibson, J. R. Altered neocortical rhythmic activity states in Fmr1 KO mice are due to enhanced mGluR5 signaling and involve changes in excitatory circuitry. J Neurosci 31, 14223–14234 (2011). https://doi.org:10.1523/JNEUROSCI.3157-11.2011

15 Ding, Q., Sethna, F. & Wang, H. Behavioral analysis of male and female Fmr1 knockout mice on C57BL/6 background. Behav Brain Res 271, 72–78 (2014). https://doi.org:10.1016/j.bbr.2014.05.046

16 van der Molen, M. J., Stam, C. J. & van der Molen, M. W. Resting-state EEG oscillatory dynamics in fragile X syndrome: abnormal functional connectivity and brain network organization. PLoS One 9, e88451 (2014). https://doi.org:10.1371/journal.pone.0088451

17 Dolen, G. et al. Correction of fragile X syndrome in mice. Neuron 56, 955–962 (2007). https://doi.org:10.1016/j.neuron.2007.12.001

18 Dölen, G. & Bear, M. F. Role for metabotropic glutamate receptor 5 (mGluR5) in the pathogenesis of fragile X syndrome. The Journal of Physiology 586, 1503–1508 (2008). https://doi.org:10.1113/jphysiol.2008.150722

19 Pan, F., Aldridge, G. M., Greenough, W. T. & Gan, W. B. Dendritic spine instability and insensitivity to modulation by sensory experience in a mouse model of fragile X syndrome. Proc Natl Acad Sci U S A 107, 17768–17773 (2010). https://doi.org:10.1073/pnas.1012496107

20 Sare, R. M. et al. Deficient Sleep in Mouse Models of Fragile X Syndrome. Front Mol Neurosci 10, 280 (2017). https://doi.org:10.3389/fnmol.2017.00280

21 Sare, R. M., Lemons, A. & Smith, C. B. Effects of Treatment With Hypnotics on Reduced Sleep Duration and Behavior Abnormalities in a Mouse Model of Fragile X Syndrome. Front Neurosci 16, 811528 (2022). https://doi.org:10.3389/fnins.2022.811528

22 Puentes-Mestril, C. & Aton, S. J. Linking network activity to synaptic plasticity during sleep: hypotheses and recent data. Frontiers in Neural Circuits 11, doi: 10.3389/fncir.2017.00061 (2017).

23 Puentes-Mestril, C., Roach, J., Niethard, N., Zochowski, M. & Aton, S. J. How rhythms of the sleeping brain tune memory and synaptic plasticity. Sleep 42, pii: zsz095 (2019). https://doi.org:10.1093/sleep/zsz095

24 Walker, M. P., Brakefield, T., Hobson, J. A. & Stickgold, R. Dissociable stages of human memory consolidation and reconsolidation. Nature 425, 616–620 (2003). https://doi.org:10.1038/nature01930

25 Stickgold, R. & Walker, M. P. Memory consolidation and reconsolidation: what is the role of sleep? Trends Neurosci 28, 408–415 (2005). https://doi.org:10.1016/j.tins.2005.06.004

26 Stickgold, R. & Walker, M. P. Sleep-dependent memory consolidation and reconsolidation. Sleep Med 8, 331–343 (2007). https://doi.org:10.1016/j.sleep.2007.03.011

27 Graves, L. A., Heller, E. A., Pack, A. I. & Abel, T. Sleep deprivation selectively impairs memory consolidation for contextual fear conditioning. Learn Mem 10, 168–176 (2003). https://doi.org:10.1101/lm.48803

28 Havekes, R. & Abel, T. The tired hippocampus: the molecular impact of sleep deprivation on hippocampal function. Curr Opin Neurobiol 44, 13–19 (2017). https://doi.org:10.1016/j.conb.2017.02.005

29 Heckman, P. R. A., Roig Kuhn, F., Meerlo, P. & Havekes, R. A brief period of sleep deprivation negatively impacts the acquisition, consolidation, and retrieval of object-location memories. Neurobiol Learn Mem 175, 107326 (2020). https://doi.org:10.1016/j.nlm.2020.107326

30 Aton, S. J. et al. Mechanisms of sleep-dependent consolidation of cortical plasticity. Neuron 61, 454–466 (2009). https://doi.org:10.1016/j.neuron.2009.01.007

31 Ognjanovski, N., Maruyama, D., Lashner, N., Zochowski, M. & Aton, S. J. CA1 hippocampal network activity changes during sleep-dependent memory consolidation. Front Syst Neurosci 8, 61 (2014). https://doi.org:10.3389/fnsys.2014.00061

32 Ognjanovski, N., Broussard, C., Zochowski, M. & Aton, S. J. Hippocampal Network Oscillations Rescue Memory Consolidation Deficits Caused by Sleep Loss. Cereb Cortex 28, 3711–3723 (2018). https://doi.org:10.1093/cercor/bhy174

33 Basta, M., Chrousos, G. P., Vela-Bueno, A. & Vgontzas, A. N. Chronic Insomnia and Stress System. Sleep Med Clin 2, 279–291 (2007). https://doi.org:10.1016/j.jsmc.2007.04.002

34 Owens, J., Adolescent Sleep Working, G. & Committee on, A. Insufficient sleep in adolescents and young adults: an update on causes and consequences. Pediatrics 134, e921–932 (2014). https://doi.org:10.1542/peds.2014-1696

35 Kotagal, S. & Broomall, E. Sleep in children with autism spectrum disorder. Pediatr Neurol 47, 242–251 (2012). https://doi.org:10.1016/j.pediatrneurol.2012.05.007

36 Wintler, T., Schoch, H., Frank, M. G. & Peixoto, L. Sleep, brain development, and autism spectrum disorders: Insights from animal models. J Neurosci Res 98, 1137–1149 (2020). https://doi.org:10.1002/jnr.24619

37 Szabadi, E. Selective targets for arousal-modifying drugs: implications for the treatment of sleep disorders. Drug Discov Today 19, 701–708 (2014). https://doi.org:10.1016/j.drudis.2014.01.001

38 Inanobe, A. et al. Characterization of G-protein-gated K+ channels composed of Kir3.2 subunits in dopaminergic neurons of the substantia nigra. J Neurosci 19, 1006–1017 (1999). https://doi.org:10.1523/JNEUROSCI.19-03-01006.1999

39 Walsh, K. B. Targeting GIRK Channels for the Development of New Therapeutic Agents. Front Pharmacol 2, 64 (2011). https://doi.org:10.3389/fphar.2011.00064

40 Hibino, H., et al. in Physiological Reviews Vol. 90 291–366 (American Physiological Society Bethesda, MD, 2010).

41 Luscher, C., Jan, L. Y., Stoffel, M., Malenka, R. C. & Nicoll, R. A. G protein-coupled inwardly rectifying K+ channels (GIRKs) mediate postsynaptic but not presynaptic transmitter actions in hippocampal neurons. Neuron 19, 687–695 (1997). https://doi.org:10.1016/s0896-6273(00)80381-5

42 Lüscher, C. & Slesinger, P. A. in Nature Reviews Neuroscience Vol. 11 301–315 (Nature Publishing Group, 2010).

43 Kaufmann, K. et al. ML297 (VU0456810), the first potent and selective activator of the GIRK potassium channel, displays antiepileptic properties in mice. ACS Chem Neurosci 4, 1278–1286 (2013). https://doi.org:10.1021/cn400062a

44 Wydeven, N. et al. Mechanisms underlying the activation of G-protein-gated inwardly rectifying K+ (GIRK) channels by the novel anxiolytic drug, ML297. Proc Natl Acad Sci U S A 111, 10755–10760 (2014). https://doi.org:10.1073/pnas.1405190111

45 Zou, B. et al. Direct activation of G-protein-gated inward rectifying K+ channels promotes nonrapid eye movement sleep. Sleep 42, 1–16 (2019). https://doi.org:10.1093/sleep/zsy244

46 Martinez, J. D., Brancaleone, W. P., Peterson, K. G., Wilson, L. G. & Aton, S. J. Atypical hypnotic compound ML297 restores sleep architecture immediately following emotionally valenced learning, to promote memory consolidation and hippocampal network activation during recall. Sleep 46 (2023). https://doi.org:10.1093/sleep/zsac301

47 Wolfe, S. A. et al. FMRP regulates an ethanol-dependent shift in GABABR function and expression with rapid antidepressant properties. Nat Communications 7, 12867 (2016).

48 Pacey, L. K. K. et al. Genetic deletion of regulator of G-protein signaling 4 (RGS4) rescues a subset of fragile X related phenotypes in the FMR1 knockout mouse. Mol Cell Neurosci 46, 563–572 (2011).

49 Trachsel, L., Tobler, I. & Borbely, A. A. Electroencephalogram analysis of non-rapid eye movement sleep in rats. Am J Physiol 255, R27–37 (1988). https://doi.org:10.1152/ajpregu.1988.255.1.R27

50 Bandarabadi, M. et al. A role for spindles in the onset of rapid eye movement sleep. Nat Commun 11, 5247 (2020). https://doi.org:10.1038/s41467-020-19076-2

51 Andrillon, T. et al. Sleep spindles in humans: insights from intracranial EEG and unit recordings. J Neurosci 31, 17821–17834 (2011). https://doi.org:10.1523/JNEUROSCI.2604-11.2011

52 Kim, A. et al. Optogenetically induced sleep spindle rhythms alter sleep architectures in mice. Proc Natl Acad Sci U S A 109, 20673–20678 (2012). https://doi.org:10.1073/pnas.1217897109

53 Durkin, J. et al. Cortically coordinated NREM thalamocortical oscillations play an essential, instructive role in visual system plasticity. Proc Natl Acad Sci U S A 114, 10485–10490 (2017). https://doi.org:10.1073/pnas.1710613114

54 Kozono, N., Okamura, A., Honda, S., Matsumoto, M. & Mihara, T. Gamma power abnormalities in a Fmr1-targeted transgenic rat model of fragile X syndrome. Sci Rep 10, 18799 (2020). https://doi.org:10.1038/s41598-020-75893-x

55 Lovelace, J. W., Ethell, I. M., Binder, D. K. & Razak, K. A. Translation-relevant EEG phenotypes in a mouse model of Fragile X Syndrome. Neurobiol Dis 115, 39–48 (2018). https://doi.org:10.1016/j.nbd.2018.03.012

56 Jonak, C. R., Lovelace, J. W., Ethell, I. M., Razak, K. A. & Binder, D. K. Multielectrode array analysis of EEG biomarkers in a mouse model of Fragile X Syndrome. Neurobiol Dis 138, 104794 (2020). https://doi.org:10.1016/j.nbd.2020.104794

57 Puentes-Mestril, C. & Aton, S. J. Linking Network Activity to Synaptic Plasticity during Sleep: Hypotheses and Recent Data. Front Neural Circuits 11, 61 (2017). https://doi.org:10.3389/fncir.2017.00061

58 Puentes-Mestril, C., Roach, J., Niethard, N., Zochowski, M. & Aton, S. J. How rhythms of the sleeping brain tune memory and synaptic plasticity. Sleep 42 (2019). https://doi.org:10.1093/sleep/zsz095

59 Lehoux, T., Carrier, J. & Godbout, R. NREM sleep EEG slow waves in autistic and typically developing children: Morphological characteristics and scalp distribution. J Sleep Res 28, e12775 (2019). https://doi.org:10.1111/jsr.12775

60 Rochette, A. C., Soulieres, I., Berthiaume, C. & Godbout, R. NREM sleep EEG activity and procedural memory: A comparison between young neurotypical and autistic adults without sleep complaints. Autism Research 11, 613–623 (2018). https://doi.org:10.1002/aur.1933

61 Gagnon, K., Bolduc, C., Bastien, L. & Godbout, R. REM Sleep EEG Activity and Clinical Correlates in Adults With Autism. Front Psychiatry 12 (2021). https://doi.org:10.3389/fpsyt.2021.659006

62 Clawson, B. C., Durkin, J. & Aton, S. J. Form and function of sleep spindles across the lifespan. Neural Plast. 2016, 6936381 (2016).

63 Clawson, B. C., Durkin, J. & Aton, S. J. Form and Function of Sleep Spindles across the Lifespan. Neural Plast 2016, 6936381 (2016). https://doi.org:10.1155/2016/6936381

64 Durkin, J. M. & Aton, S. J. How Sleep Shapes Thalamocortical Circuit Function in the Visual System. Annu Rev Vis Sci 5, 295–315 (2019). https://doi.org:10.1146/annurev-vision-091718-014715

65 Aton, S. J. et al. Visual experience and subsequent sleep induce sequential plastic changes in putative inhibitory and excitatory cortical neurons. Proc Natl Acad Sci U S A 110, 3101–3106 (2013).

66 Wamsley, E. J. et al. The effects of eszopiclone on sleep spindles and memory consolidation in schizophrenia: a randomized placebo-controlled trial. Sleep 36, 1369–1376 (2013).

67 Blumberg, M. S., Dooley, J. C. & Sokoloff, G. The developing brain revealed during sleep. Curr Opin Physiol 15, 14–22 (2020).

68 Lambert, A. et al. Poor sleep affects daytime functioning in typically developing and autistic children not complaining of sleep problems: A questionnaire-based and polysomnographic study. Research in Autism Spectrum Disorders 23, 94–106 (2016).

69 Tessier, S. et al. Intelligence measures and stage 2 sleep in typically-developing and autistic children. Int J Psychophysiol 97, 58–65 (2015). https://doi.org:10.1016/j.ijpsycho.2015.05.003

70 Delorme, J. et al. Sleep loss drives acetylcholine- and somatostatin interneuron-mediated gating of hippocampal activity to inhibit memory consolidation. Proc Natl Acad Sci U S A 118 (2021). https://doi.org:10.1073/pnas.2019318118

71 Vecsey, C. G. et al. Sleep deprivation impairs cAMP signalling in the hippocampus. Nature 461, 1122–1125 (2009). https://doi.org:10.1038/nature08488

72 Havekes, R. et al. Sleep deprivation causes memory deficits by negatively impacting neuronal connectivity in hippocampal area CA1. Elife 5, e13424–e13424 (2016). https://doi.org:10.7554/eLife.13424

73 Dam, D. V. et al. in Behavioural Brain Research Vol. 117 127–136 (2000).

74 Li, J. et al. Defective memory engram reactivation underlies impaired fear memory recall in Fragile X syndrome. Elife 9 (2020). https://doi.org:10.7554/eLife.61882

75 Seese, R. R., Wang, K., Yao, Y. Q., Lynch, G. & Gall, C. M. Spaced training rescues memory and ERK1/2 signaling in fragile X syndrome model mice. Proc Natl Acad Sci U S A 111, 16907–16912 (2014). https://doi.org:10.1073/pnas.1413335111

76 Liu, X. et al. Optogenetic stimulation of a hippocampal engram activates fear memory recall. Nature 484, 381–385 (2012). https://doi.org:10.1038/nature11028

77 Martinez, J. D., et al. Hypnotic treatment restores sleep architecture and sleep-dependent memory consolidation in a mouse model of Fragile X syndrome. . BioRxiv (2023).

78 Lee, F. H. F., Lai, T. K. Y., Su, P. & Liu, F. Altered cortical Cytoarchitecture in the Fmr1 knockout mouse. Mol Brain 12, 56 (2019). https://doi.org:10.1186/s13041-019-0478-8

79 Kalinowska, M. et al. Deletion of Fmr1 in parvalbumin-expressing neurons results in dysregulated translation and selective behavioral deficits associated with fragile X syndrome. Mol Autism 13, 29 (2022). https://doi.org:10.1186/s13229-022-00509-2

80 Boone, C. E., Davoudi, H., Harrold, J. B. & Foster, D. J. Abnormal Sleep Architecture and Hippocampal Circuit Dysfunction in a Mouse Model of Fragile X Syndrome. Neuroscience 384, 275–289 (2018). https://doi.org:10.1016/j.neuroscience.2018.05.012

81 Elia, M. et al. Sleep in subjects with autistic disorder: a neurophysiological and psychological study. Brain Dev 22, 88–92 (2000). https://doi.org:10.1016/s0387-7604(99)00119-9

82 Limoges, E., Mottron, L., Bolduc, C., Berthiaume, C. & Godbout, R. Atypical sleep architecture and the autism phenotype. Brain 128, 1049–1061 (2005). https://doi.org:10.1093/brain/awh425

83 Johnson, K. P., Giannotti, F. & Cortesi, F. Sleep patterns in autism spectrum disorders. Child Adolesc Psychiatr Clin N Am 18, 917–928 (2009). https://doi.org:10.1016/j.chc.2009.04.001

84 Picchioni, D., Reith, R. M., Nadel, J. L. & Smith, C. B. Sleep, plasticity and the pathophysiology of neurodevelopmental disorders: the potential roles of protein synthesis and other cellular processes. Brain Sci 4, 150–201 (2014). https://doi.org:10.3390/brainsci4010150

85 Accardo, J. A. & Malow, B. A. Sleep, epilepsy, and autism. Epilepsy Behav 47, 202–206 (2015). https://doi.org:10.1016/j.yebeh.2014.09.081

86 Goel, A. et al. Impaired perceptual learning in a mouse model of Fragile X syndrome is mediated by parvalbumin neuron dysfunction and is reversible. Nat Neurosci 21, 1404–1411 (2018). https://doi.org:10.1038/s41593-018-0231-0

87 Sabanov, V. et al. Impaired GABAergic inhibition in the hippocampus of Fmr1 knockout mice. Neuropharmacology 116, 71–81 (2017). https://doi.org:10.1016/j.neuropharm.2016.12.010

88 Angriman, M., Caravale, B., Novelli, L., Ferri, R. & Bruni, O. Sleep in children with neurodevelopmental disabilities. Neuropediatrics 46, 199–210 (2015). https://doi.org:10.1055/s-0035-1550151

89 Contractor, A., Klyachko, V. A. & Portera-Cailliau, C. Altered Neuronal and Circuit Excitability in Fragile X Syndrome. Neuron 87, 699–715 (2015). https://doi.org:10.1016/j.neuron.2015.06.017

90 Paluszkiewicz, S. M., Martin, B. S. & Huntsman, M. M. Fragile X syndrome: the GABAergic system and circuit dysfunction. Dev Neurosci 33, 349–364 (2011). https://doi.org:10.1159/000329420

91 Contractor, A., Ethell, I. M. & Portera-Cailliau, C. Cortical interneurons in autism. Nat Neurosci 24, 1648–1659 (2021). https://doi.org:10.1038/s41593-021-00967-6

92 Vantomme, G., Osorio-Forero, A., Luthi, A. & Fernandez, L. M. J. Regulation of Local Sleep by the Thalamic Reticular Nucleus. Front Neurosci 13, 576 (2019). https://doi.org:10.3389/fnins.2019.00576

93 Sun, Q.-Q., Huguenard, J. R. & Prince, D. A. Somatostatin Inhibits Thalamic Network Oscillations In Vitro: Actions on the GABAergic Neurons of the Reticular Nucleus. J.Neurosci. 22, 5374–5386 (2002).

94 Diekelmann, S. & Born, J. in Nature Reviews Neuroscience Vol. 11 114–126 (2010).

95 Rasch, B. & Born, J. About sleep’s role in memory. Physiol Rev 93, 681–766 (2013). https://doi.org:10.1152/physrev.00032.2012

96 Bridi, M. C. D. et al. Daily Oscillation of the Excitation-Inhibition Balance in Visual Cortical Circuits. Neuron 105, 621–629 (2020).

97 Pantazopoulos, H. et al. Circadian Rhythms of Perineuronal Net Composition. eNeuro 7 (2020). https://doi.org:10.1523/ENEURO.0034-19.2020

98 Ognjanovski, N., Maruyama, D., Lashner, N., Zochowski, M. & Aton, S. J. CA1 hippocampal network activity changes during sleep-dependent memory consolidation. Front Syst Neurosci 8, 61 (2014).

99 Ognjanovski, N. et al. Parvalbumin-expressing interneurons coordinate hippocampal network dynamics required for memory consolidation. Nature Communications 8 (2017).

100 Ognjanovski, N., Broussard, C., Zochowski, M. & Aton, S. J. Hippocampal Network Oscillations Rescue Memory Consolidation Deficits Caused by Sleep Loss. Cereb. Cortex 28, 3711–3723 (2018). https://doi.org:10.1093/cercor/bhy174

101 Hill, E., Hickman, C., Diez, R. & Wall, M. Role of A1 receptor-activated GIRK channels in the suppression of hippocampal seizure activity. Neuropharmacology 164, 107904 (2020). https://doi.org:10.1016/j.neuropharm.2019.107904

102 Frank, M. G. Sleep and developmental plasticity not just for kids. Prog Brain Res 193, 221–232 (2011). https://doi.org:10.1016/B978-0-444-53839-0.00014-4

103 Frank, M. G., Issa, N. P. & Stryker, M. P. Sleep enhances plasticity in the developing visual cortex. Neuron 30, 275–287 (2001). https://doi.org:10.1016/s0896-6273(01)00279-3

104 Ognjanovski, N. et al. Parvalbumin-expressing interneurons coordinate hippocampal network dynamics required for memory consolidation. Nat Commun 8, 15039 (2017). https://doi.org:10.1038/ncomms15039

105 Delorme, J. et al. Hippocampal neurons’ cytosolic and membrane-bound ribosomal transcript profiles are differentially regulated by learning and subsequent sleep. Proc Natl Acad Sci U S A 118 (2021). https://doi.org:10.1073/pnas.2108534118

106 Raven, F., Heckman, P. R. A., Havekes, R. & Meerlo, P. Sleep deprivation-induced impairment of memory consolidation is not mediated by glucocorticoid stress hormones. J Sleep Res 29, e12972 (2020). https://doi.org:10.1111/jsr.12972

107 Seibenhener, M. L. & Wooten, M. C. Use of the Open Field Maze to measure locomotor and anxiety-like behavior in mice. J Vis Exp, e52434 (2015). https://doi.org:10.3791/52434

108 Hui, G. K. et al. Memory enhancement of classical fear conditioning by post-training injections of corticosterone in rats. Neurobiol Learn Mem 81, 67–74 (2004). https://doi.org:10.1016/j.nlm.2003.09.002

109 Roozendaal, B., Okuda, S., Van der Zee, E. A. & McGaugh, J. L. Glucocorticoid enhancement of memory requires arousal-induced noradrenergic activation in the basolateral amygdala. Proc Natl Acad Sci U S A 103, 6741–6746 (2006). https://doi.org:10.1073/pnas.0601874103

110 Delorme, J. E., Kodoth, V. & Aton, S. J. Sleep loss disrupts Arc expression in dentate gyrus neurons. Neurobiol Learn Mem 160, 73–82 (2019). https://doi.org:10.1016/j.nlm.2018.04.006

111 Fisher, S. P. et al. Rapid assessment of sleep-wake behavior in mice. J Biol Rhythms 27, 48–58 (2012). https://doi.org:10.1177/0748730411431550

112 Puentes-Mestril, C. et al. Sleep Loss Drives Brain Region-Specific and Cell Type-Specific Alterations in Ribosome-Associated Transcripts Involved in Synaptic Plasticity and Cellular Timekeeping. J Neurosci 41, 5386–5398 (2021). https://doi.org:10.1523/JNEUROSCI.1883-20.2021

113 Clawson, B. C. et al. Causal role for sleep-dependent reactivation of learning-activated sensory ensembles for fear memory consolidation. Nat Commun 12, 1200 (2021). https://doi.org:10.1038/s41467-021-21471-2

114 Kattla, S. & Lowery, M. M. Fatigue related changes in electromyographic coherence between synergistic hand muscles. Exp Brain Res 202, 89–99 (2010). https://doi.org:10.1007/s00221-009-2110-0

115 Swift, K. M. et al. Sex differences within sleep in gonadally intact rats. Sleep 43, 1–14 (2020). https://doi.org:10.1093/sleep/zsz289

116 Pham, J., Cabrera, S. M., Sanchis-Segura, C. & Wood, M. A. Automated scoring of fear-related behavior using EthoVision software. J Neurosci Methods 178, 323–326 (2009). https://doi.org:10.1016/j.jneumeth.2008.12.021

