## Supplemental material for "Hypnotic treatment reverses NREM sleep disruption and EEG desynchronization in a mouse model of Fragile X syndrome to rescue memory consolidation deficits"

### Title:

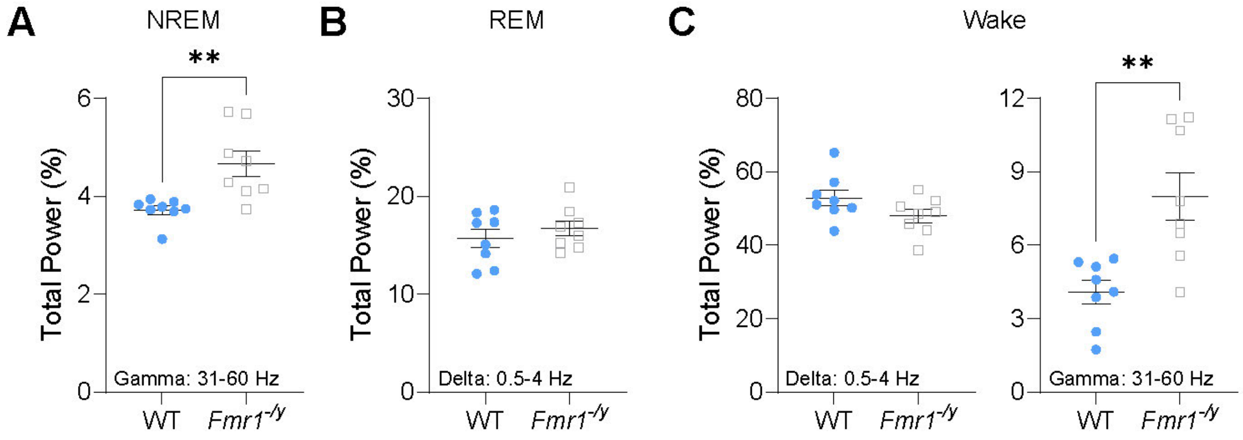


**Figure S1: *Fmr1$^{-/y}$* mice NREM gamma, REM delta, and waking spectral power during the rest phase.**

**(A)** Total gamma-band (31-100 Hz) power was increased in *Fmr1$^{-/y}$* mice during NREM sleep across the 12-h light phase.

**(B)** Total delta-band (0.5-4 Hz) power during REM sleep across the 12-h light phase.

**(C)** Total delta-band (***left***) and gamma-band (***right***) power were reduced and increased, respectively, in *Fmr1$^{-/y}$* mice during wake across the 12-h light phase.

Sample size: $n$ = 8 mice/genotype. ** indicates $p < 0.01$, two-tailed, unpaired t-test (**E-H**). Data points and error bars indicate mean ± SEM.

(Related to Fig. 1)

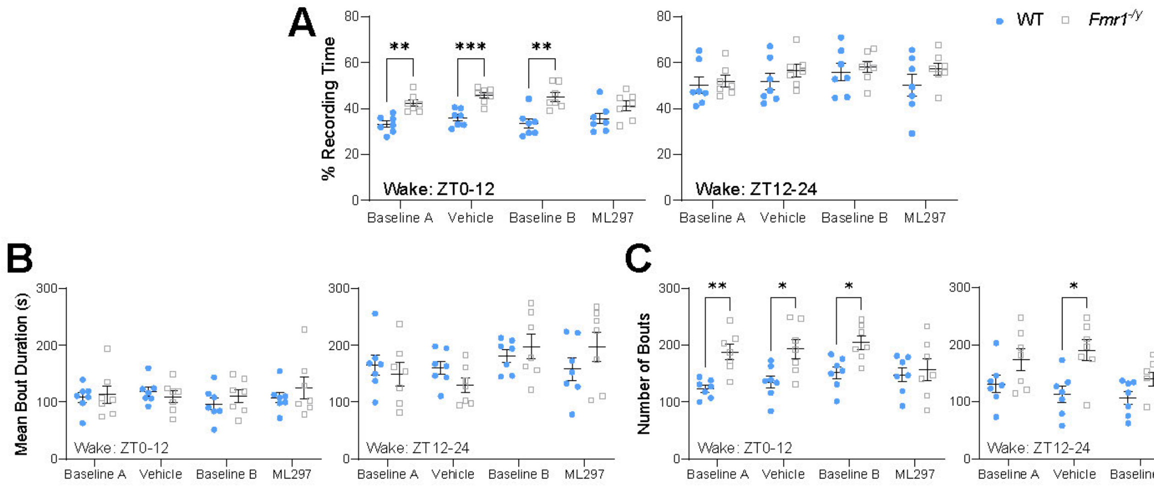


**Figure S2: ML297 normalizes wake architecture differences in *Fmr1$^{-/y}$* mice.**

**(A)** Waking percent recording time for the 0-12 h light phase (***left***) and 12-24 h dark phase (***right***) across all four days for WT and *Fmr1$^{-/y}$* mice. Two-way RM ANOVA for ZT0-12: *p*(condition) = 0.24, *p*(genotype) = 0.0002, *p*(condition x genotype interaction) = 0.27 and ZT12-24: *p*(condition) = 0.035, *p*(genotype) = 0.35, *p*(condition x genotype interaction) = 0.41.

**(B)** Waking bout duration for the 0-12 h light phase (***left***) and 12-24 h dark phase (***right***) across all four days for WT and *Fmr1$^{-/y}$* mice. Two-way RM ANOVA for ZT0-12: *p*(condition) = 0.35, *p*(genotype) = 0.64, *p*(condition x genotype interaction) = 0.31 and ZT12-24: *p*(condition) = 0.0037, *p*(genotype) = 0.93, *p*(condition x genotype interaction) = 0.025.

**(C)** Number of wakefulness bouts for the 0-12 h light phase (***left***) and 12-24 h dark phase (***right***) across all four days for WT and *Fmr1$^{-/y}$* mice. Two-way RM ANOVA for ZT0-12: *p*(condition) = 0.039, *p*(genotype) = 0.0097, *p*(condition x genotype interaction) = 0.011 and ZT12-24: *p*(condition) = 0.052, *p*(genotype) = 0.023, *p*(condition x genotype interaction) = 0.027.

Sample size: *n* = 7 mice/genotype. *, ** and *** indicate $p < 0.05$, $p < 0.01$, and $p < 0.001$, Sidak's *post hoc* test. Data points and error bars indicate mean ± SEM.

(Related to Fig. 2)

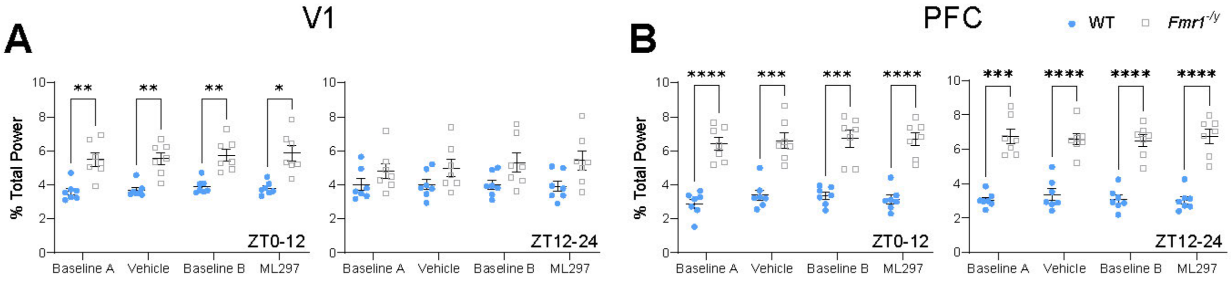


**Figure S3: NREM gamma power is increased in V1 and PFC in WT and *Fmr1$^{-/y}$* mice.**

(**A**) Total NREM gamma-band power (31-60 Hz) during the light phase (ZT0-12) (***left***) and dark phase (ZT12-24) (***right***) in V1. Two-way RM ANOVA for ZT0-12: $p$(condition) = 0.12, $p$(genotype) = 0.0004, $p$(condition x genotype interaction) = 0.79 and ZT12-24: $p$(condition) = 0.19, $p$(genotype) = 0.073, $p$(condition x genotype interaction) = 0.054.

(**B**) Total NREM gamma-band power (31-60 Hz) during the light phase (ZT0-12) (***left***) and dark phase (ZT12-24) (***right***) in PFC. Two-way RM ANOVA for ZT0-12: $p$(condition) = 0.23, $p$(genotype) < 0.0001, $p$(condition x genotype interaction) = 0.79 and ZT12-24: $p$(condition) = 0.80, $p$(genotype) < 0.0001, $p$(condition x genotype interaction) = 0.58.

Sample size: $n$ = 7 mice/genotype. *, **, *** and **** indicate $p < 0.05$, $p < 0.01$, $p < 0.001$, and $p < 0.0001$, Sidak's *post hoc* test. Data points and error bars indicate mean ± SEM.

(Related to Fig. 3)

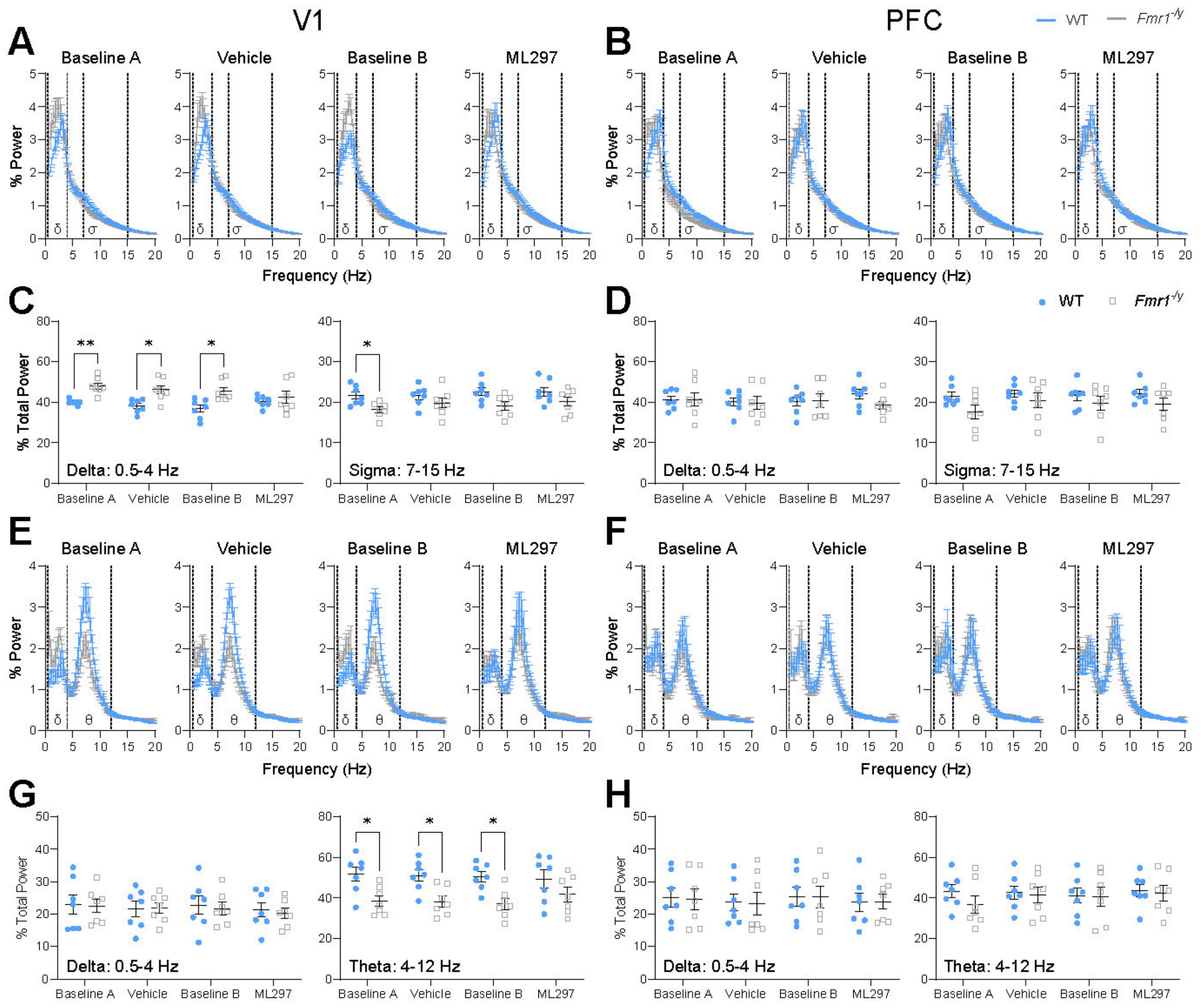


**Figure S4: NREM and REM spectral power in V1, but not PFC, is altered in *Fmr1*$^{-/y}$ mice and normalized with ML297 during the dark phase.**

**(A)** NREM EEG power spectra in V1 for baselines and treatment conditions during ZT12-24 (dark phase) with delta (δ; 0.5-4 Hz) and sigma (σ; 7-15 Hz) frequency bands highlighted via dashed lines. Two-way RM ANOVA, *p*(frequency) < 0.0001, *p*(frequency x genotype interaction) < 0.0001, for baseline A-ML297 conditions and *p*(genotype) = 0.11 (baseline A), = 0.002 (vehicle), = 0.072 (baseline B), and = 0.26 (ML297).

**(B)** NREM EEG power spectra in PFC for baselines and treatment conditions during ZT12-24 (dark phase) with delta (δ; 0.5-4 Hz) and sigma (σ; 7-15 Hz) frequency bands highlighted via dashed lines. Two-way RM ANOVA, *p*(frequency) < 0.0001, *p*(frequency x genotype interaction) < 0.0001, for baseline A-ML297 conditions and *p*(genotype) = 0.091 (baseline A), = 0.33 (vehicle), = 0.42 (baseline B), and = 0.001 (ML297).

Sample size: *n* = 7 mice/genotype. * and ** indicate *p* < 0.05 and *p* < 0.01, Sidak's *post hoc* test. Data points and error bars indicate mean ± SEM.

(Related to Fig. 3)

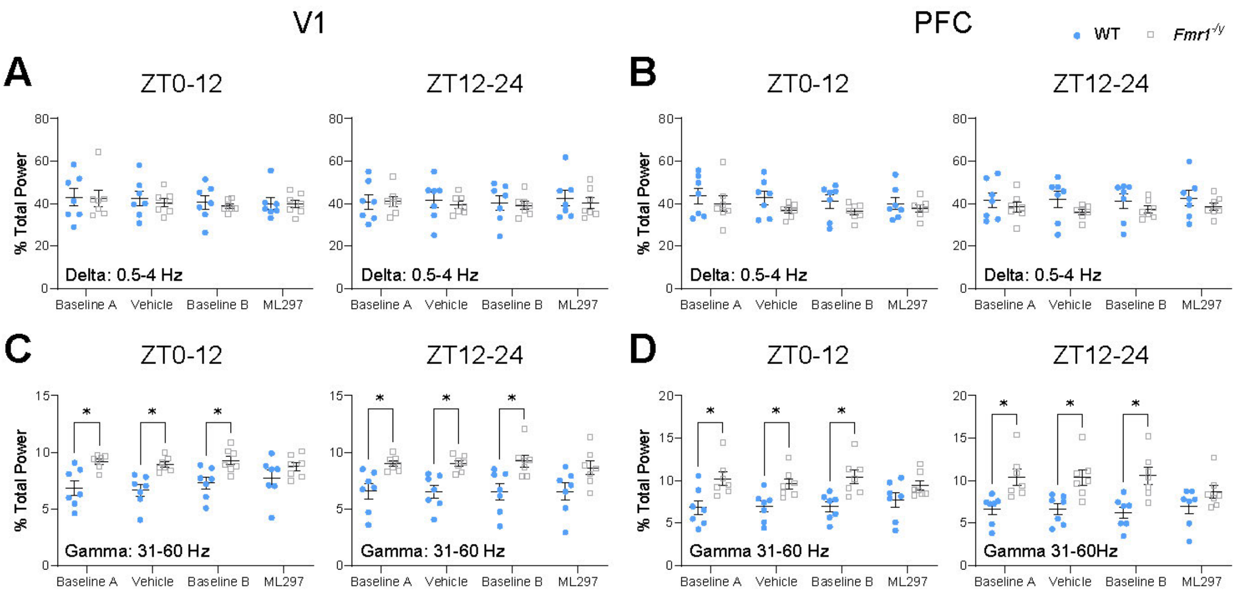


**Figure S5: Gamma power during wake is altered in *Fmr1$^{-/y}$* mice, and normalized by ML297.**

(**A**) Total delta-band power (0.5-4 Hz) during wakefulness for ZT0-12 (***left***) and ZT12-24 (***right***) in V1. Two-way RM ANOVA for ZT0-12: *p*(condition) = 0.26, *p*(genotype) = 0.78, *p*(condition x genotype interaction) = 0.93 and ZT12-24: *p*(condition) = 0.70, *p*(genotype) = 0.71, *p*(condition x genotype interaction) = 0.83.

(**B**) Total delta-band power (0.5-4 Hz) during wakefulness for ZT0-12 (***left***) and ZT12-24 (***right***) in PFC. Two-way RM ANOVA for ZT0-12: *p*(condition) = 0.26, *p*(genotype) = 0.24, *p*(condition x genotype interaction) = 0.72 and ZT12-24: *p*(condition) = 0.66, *p*(genotype) = 0.25, *p*(condition x genotype interaction) = 0.82.

(**C**) Total gamma-band power (31-60 Hz) during wakefulness for ZT0-12 (***left***) and ZT12-24 (***right***) in V1. Two-way RM ANOVA for ZT0-12: *p*(condition) = 0.37, *p*(genotype) = 0.0043, *p*(condition x genotype interaction) = 0.20 and ZT12-24: *p*(condition) = 0.90, *p*(genotype) = 0.0007, *p*(condition x genotype interaction) = 0.95.

(**D**) Total gamma-band power (31-60 Hz) during wakefulness for ZT0-12 (***left***) and ZT12-24 (***right***) in V1. Two-way RM ANOVA for ZT0-12: *p*(condition) = 0.51, *p*(genotype) = 0.012, *p*(condition x genotype interaction) = 0.017 and ZT12-24: *p*(condition) = 0.24, *p*(genotype) = 0.0065, *p*(condition x genotype interaction) = 0.013.

Sample size: *n* = 7 mice/genotype. * indicates $p < 0.05$, Sidak's *post hoc* test. Data points and error bars indicate mean ± SEM.

(Related to Fig. 3)

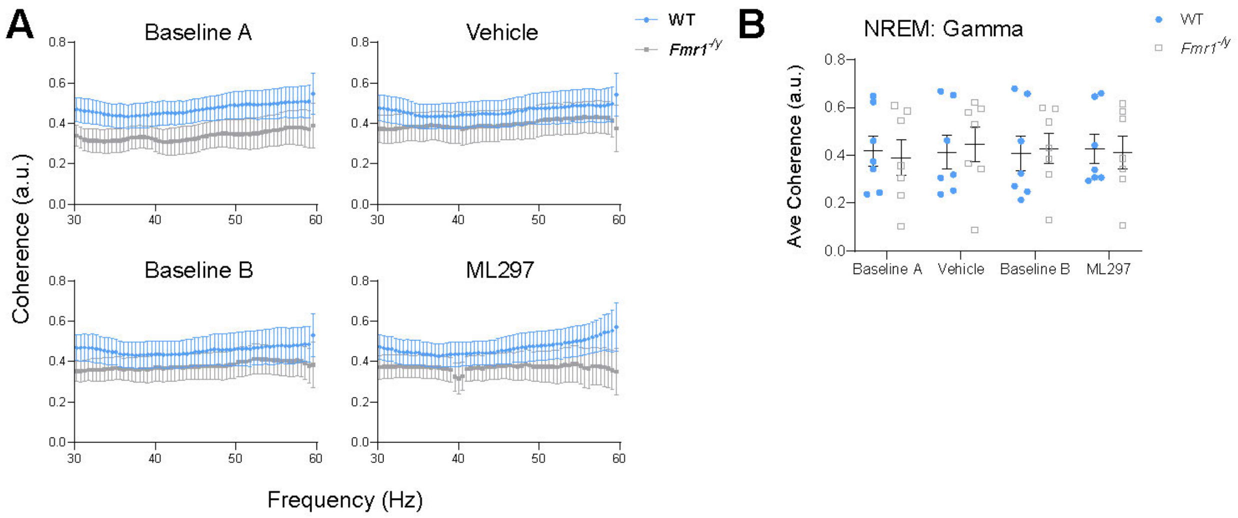


**Figure S6: NREM gamma coherence between WT and *Fmr1*$^{-/y}$ mice.**

**(A)** NREM-associated V1-PFC spectral coherence for 31-60 Hz across baseline and treatment sessions. Two-way ANOVA, $p$(frequency) > 0.99, $p$(genotype) < 0.0001, $p$(frequency x genotype interaction) > 0.99, for baseline A-ML297 conditions.

**(B)** Average NREM V1-PFC gamma-band (31-60 Hz) coherence. Two-way RM ANOVA: $p$(condition) = 0.49, $p$(genotype) = 0.98, $p$(condition x genotype interaction) = 0.34.

Sample size: $n$ = 7 mice/genotype. Data points and error bars indicate mean ± SEM.

(Related to Fig. 5)

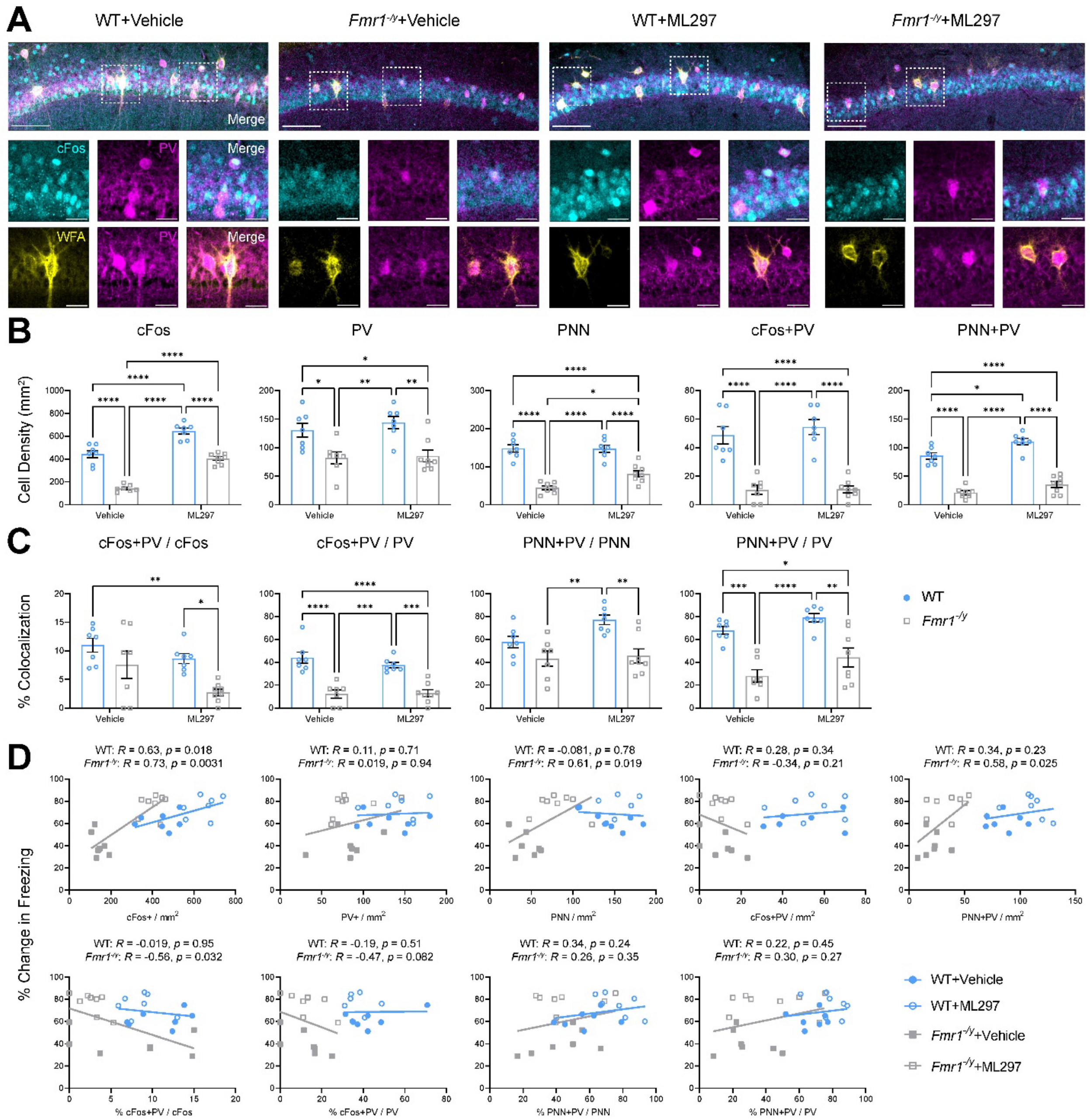

**Figure S7: *Fmr1*$^{-/y}$ mice show reduced CA1 network activity during memory recall, which is not rescued by ML297 administration.**

**(A)** Representative fear memory recall-associated cFos (cyan), PV (magenta), and wisteria floribunda agglutinin (WFA; yellow) for PNN expression in CA1 for the four treatment groups. Scale bar = 250 μm (***top***, whole CA1); scale bar = 30 μm (***bottom***, inset of select neurons).

**(B)** Quantification of CA1 cFos+, PV+, PNN, and co-localization neuronal density. Two-way ANOVA, for cFos+ neurons: $p$(treatment) < 0.0001, $p$(genotype) < 0.0001, $p$(treatment x genotype interaction) = 0.21. For PV+ neurons: $p$(treatment) = 0.44, $p$(genotype) < 0.0001, $p$(treatment x genotype interaction) = 0.66. For PNN: $p$(treatment) = 0.039, $p$(genotype) < 0.0001, $p$(treatment x genotype interaction) = 0.033. For cFos+PV co-localized neurons: *p*(treatment) = 0.48, *p*(genotype) < 0.0001, *p*(treatment x genotype interaction) = 0.54. For PNN+PV co-localized neurons: *p*(treatment) = 0.0008, *p*(genotype) < 0.0001, *p*(treatment x genotype interaction) = 0.30.

**(C)** Quantification of CA1 percent co-localization of cFos+, PV+, and PNN over total neuronal populations. Two-way ANOVA, for % cFos+PV cells over total cFos: *p*(treatment) = 0.018, *p*(genotype) = 0.0029, *p*(treatment x genotype interaction) = 0.38. For % cFos+PV cells over total PV: *p*(treatment) = 0.42, *p*(genotype) < 0.0001, *p*(treatment x genotype interaction) = 0.32. For % PNN+PV cells over total PNN: *p*(treatment) = 0.065, *p*(genotype) = 0.0004, *p*(treatment x genotype interaction) = 0.15. For % PNN+PV cells over total PV: *p*(treatment) = 0.026, *p*(genotype) < 0.0001, *p*(treatment x genotype interaction) = 0.67.

**(D)** Pearson correlation of cFos+, PV+, and PNN neuron densities and associated co-localizations in CA1 vs. associated change in freezing behavior during CFM recall. *R* and *p* values for Pearson correlation of each data set are found alongside corresponding graph.

Sample sizes: *n* = 7 (WT+vehicle), *n* = 7 ($Fmr1^{-/y}$+vehicle), *n* = 7 (WT+ML297), *n* = 8 ($Fmr1^{-/y}$+ML297). *, **, ***, and **** indicate *p* < 0.05, *p* < 0.01, *p* < 0.001, *p* < 0.0001; Tukey's *post hoc* test. Data shown as mean ± SEM.

(Related to Fig. 8)

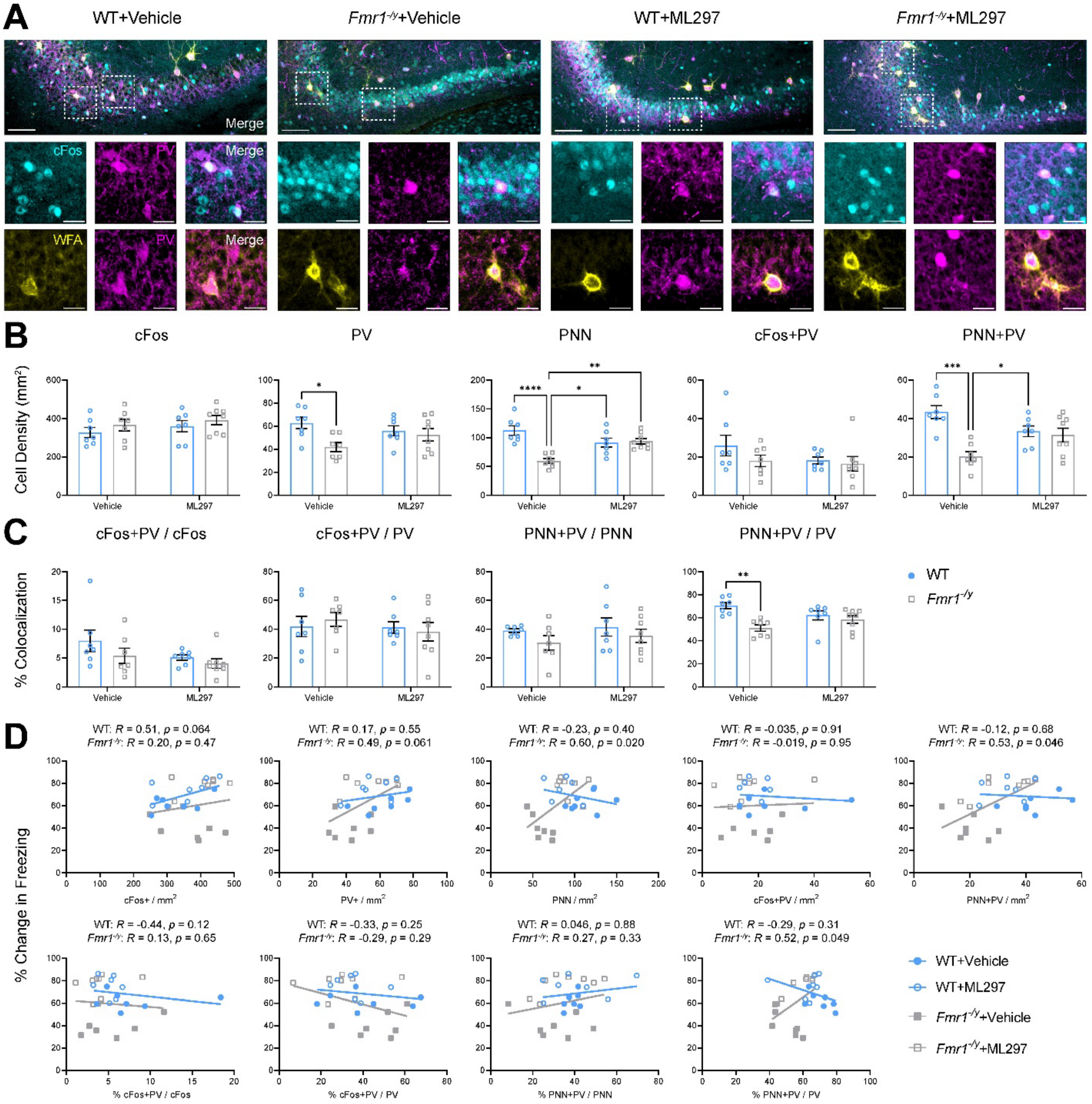

**Figure S8: *Fmr1$^{-/y}$* mice show alterations to CA3 PV+ interneurons and PNNs that are normalized by ML297.**

**(A)** Representative fear memory recall-associated cFos (cyan), PV (magenta), and wisteria floribunda agglutinin (WFA; yellow) for PNN expression in CA3 for the four treatment groups. Scale bar = 250 µm (***top***, whole CA3); scale bar = 30 µm (***bottom***, inset of select neurons).

**(B)** Quantification of CA3 cFos+, PV+, PNN, and co-localization neuronal density. Two-way ANOVA, for cFos+ neurons: *p*(treatment) = 0.30, *p*(genotype) = 0.21, *p*(treatment x genotype interaction) = 0.91. For PV+ neurons: *p*(treatment) = 0.68, *p*(genotype) = 0.018, *p*(treatment x genotype interaction) = 0.083. For PNN: $p$(treatment) = 0.32, $p$(genotype) = 0.0006, $p$(treatment x genotype interaction) = 0.0002. For cFos+PV co-localized neurons: $p$(treatment) = 0.23, $p$(genotype) = 0.21, $p$(treatment x genotype interaction) = 0.41. For PNN+PV co-localized neurons: $p$(treatment) = 0.86, $p$(genotype) = 0.0005, $p$(treatment x genotype interaction) = 0.0026.

**(C)** Quantification of CA3 percent co-localization of cFos+, PV+, and PNN over total neuronal populations. Two-way ANOVA, for % cFos+PV cells over total cFos: $p$(treatment) = 0.093, $p$(genotype) = 0.14, $p$(treatment x genotype interaction) = 0.52. For % cFos+PV cells over total PV: $p$(treatment) = 0.044, $p$(genotype) = 0.95, $p$(treatment x genotype interaction) = 0.33. For % PNN+PV cells over total PNN: $p$(treatment) = 0.43, $p$(genotype) = 0.13, $p$(treatment x genotype interaction) = 0.79. For % PNN+PV cells over total PV: $p$(treatment) = 0.87, $p$(genotype) = 0.0014, $p$(treatment x genotype interaction) = 0.022.

**(D)** Pearson correlation of cFos+, PV+, and PNN neuron densities and associated co-localizations in CA3 vs. associated change in freezing behavior during CFM recall. $R$ and $p$ values for Pearson correlation of each data set are found alongside corresponding graph.

Sample sizes: $n$ = 7 (WT+vehicle), $n$ = 7 (*Fmr1*$^{-/y}$+vehicle), $n$ = 7 (WT+ML297), $n$ = 8 (*Fmr1*$^{-/y}$+ML297). *, **, ***, and **** indicate $p < 0.05$, $p < 0.01$, $p < 0.001$, $p < 0.0001$; Tukey's *post hoc* test. Data shown as mean ± SEM.

(Related to Fig. 8)

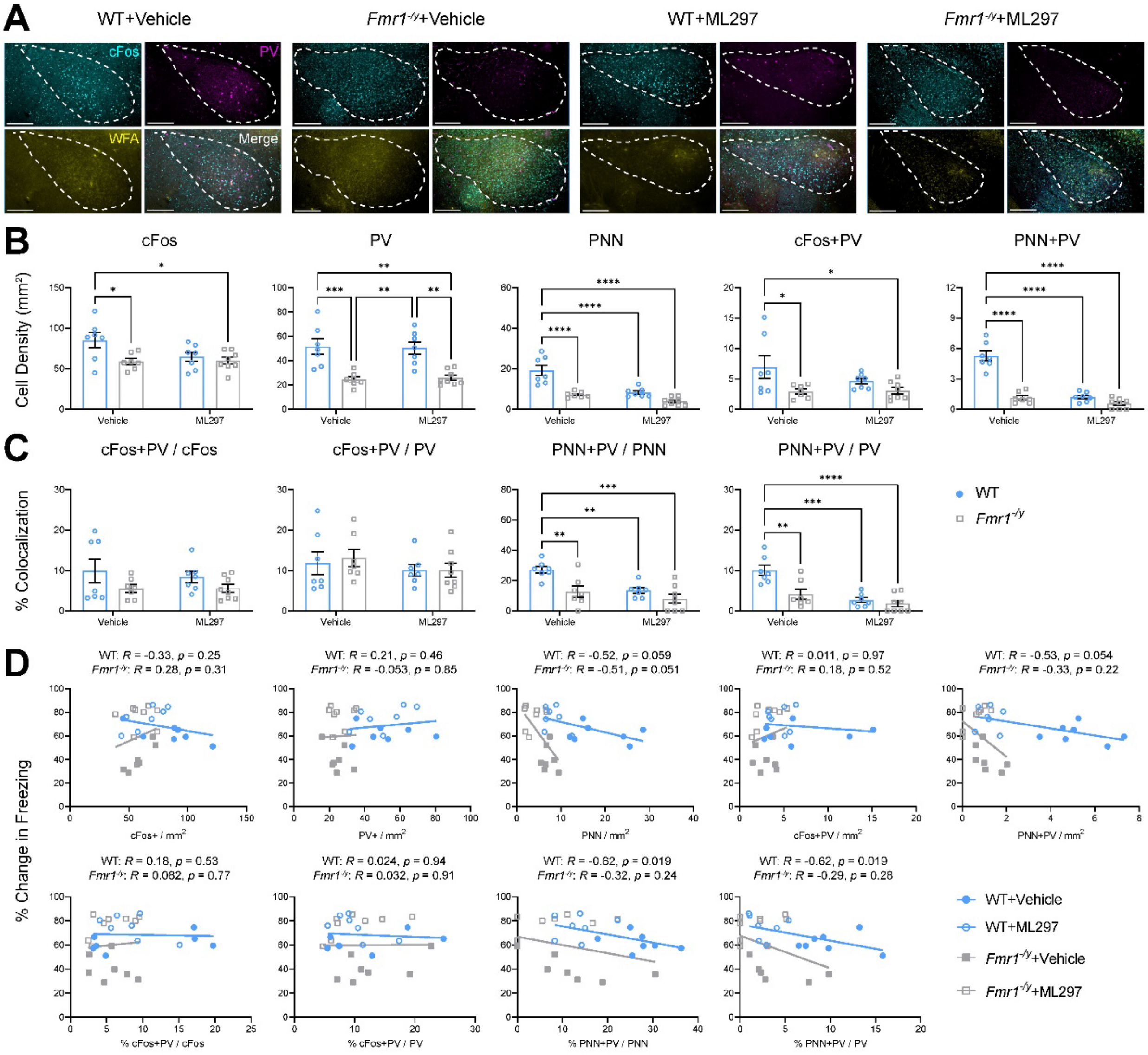


**Figure S9: *Fmr1$^{-/y}$* mice show altered amygdala activity and PV+ interneuron density, which are not rescued by ML297 administration.**

**(A)** Representative fear memory recall-associated cFos (cyan), PV (magenta), and wisteria floribunda agglutinin (WFA; yellow) for PNN expression in the amygdala for the four treatment groups. Scale bar = 250 µm (***top***, amygdala); scale bar = 30 µm (***bottom***, inset of select neurons).

**(B)** Quantification of cFos+, PV+, PNN, and co-localization neuronal density. Two-way ANOVA, for cFos+ neurons: $p$(treatment) = 0.12, $p$(genotype) = 0.018, $p$(treatment x genotype interaction) = 0.085. For PV+ neurons: $p$(treatment) = 0.98, $p$(genotype) < 0.0001, $p$(treatment x genotype interaction) = 0.77. For PNN: $p$(treatment) < 0.0001, $p$(genotype) < 0.0001, $p$(treatment x genotype interaction) = 0.01. For cFos+PV co-localized neurons: $p$(treatment) = 0.26, $p$(genotype)

= 0.0088, $p$(treatment x genotype interaction) = 0.23. For PNN+PV co-localized neurons: $p$(treatment) < 0.0001, $p$(genotype) < 0.0001, $p$(treatment x genotype interaction) < 0.0001.

**(C)** Quantification of percent co-localization of cFos+, PV+, and PNN over total neuronal populations. Two-way ANOVA, for % cFos+PV cells over total cFos: $p$(treatment) = 0.67, $p$(genotype) = 0.048, $p$(treatment x genotype interaction) = 0.66. For % cFos+PV cells over total PV: $p$(treatment) = 0.27, $p$(genotype) = 0.76, $p$(treatment x genotype interaction) = 0.77. For % PNN+PV cells over total PNN: $p$(treatment) = 0.0027, $p$(genotype) = 0.0013, $p$(treatment x genotype interaction) = 0.11. For % PNN+PV cells over total PV: $p$(treatment) < 0.0001, $p$(genotype) = 0.0026, $p$(treatment x genotype interaction) = 0.018.

**(D)** Pearson correlation of cFos+, PV+, and PNN neuron densities and associated co-localizations in amygdala vs. associated change in freezing behavior during CFM recall. $R$ and $p$ values for Pearson correlation of each data set are found alongside corresponding graph.

Sample sizes: $n$ = 7 (WT+vehicle), $n$ = 7 (*Fmr1*$^{-/y}$+vehicle), $n$ = 7 (WT+ML297), $n$ = 8 (*Fmr1*$^{-/y}$+ML297). *, **, ***, and **** indicate $p < 0.05$, $p < 0.01$, $p < 0.001$, $p < 0.0001$; Tukey's *post hoc* test. Data shown as mean ± SEM.

(Related to Fig. 8)
